# Mechanotherapeutic Potential of Survivin in Glioblastoma

**DOI:** 10.64898/2026.03.18.712467

**Authors:** Gabrielle Inserra, Sarah Balghonaim, Jessica Jong, Rhonda Drewes, Briana A Santo, Bat-Ider Tumenbayar, Khanh Pham, Sefunmi Babatunde, John E Tomaszewski, Tracey A Ignatowski, Ruogang Zhao, Jaims Lim, Sunghan Kim, Adnan H Siddiqui, Bhaskar C Das, Vincent M Tutino, Yongho Bae

## Abstract

Glioblastoma Multiforme (GBM) is a highly aggressive brain cancer characterized by rapid proliferation and extensive remodeling of the extracellular matrix (ECM), leading to progressive tissue stiffening. Although ECM stiffness is known to promote GBM progression, the molecular mechanisms linking mechanical cues to tumor growth remain insufficiently defined. In this study, transcriptomic comparison of GBM tumors and non-neoplastic brain tissue revealed coordinated upregulation of cell cycle regulators and matrisome-associated genes, with survivin (BIRC5) identified as a central node linking proliferative signaling and ECM remodeling networks. Analysis of GBM patient specimens further showed strong nuclear survivin expression in regions with elevated collagen deposition. To directly evaluate stiffness-dependent regulation of survivin, GBM cells were cultured on fibronectin-infused hydrogels with tunable stiffness. Stiff matrices increased survivin expression along with cyclin D1 and cyclin A, consistent with increased cell cycle progression. Pharmacologic inhibition or siRNA-mediated suppression of survivin reduced stiffness-induced proliferation and attenuated expression of matrisome components, including collagens and lysyl oxidase. These findings indicate that survivin functions as a mechanosensitive regulator that coordinates cell cycle progression with ECM production in stiff tumor microenvironments. Collectively, this study identifies survivin as a key mediator linking ECM stiffness to GBM growth and matrisome remodeling. Targeting survivin and its effectors may offer a mechanosensitive strategy to limit GBM growth.

## INTRODUCTION

Glioblastoma Multiforme (GBM) is an isocitrate dehydrogenase (IDH) wildtype World Health Organization (WHO) grade IV astrocytoma and the most aggressive primary brain tumor in adults [1–3]. GBM is characterized by rapid proliferation, diffuse infiltration into healthy brain tissue, active remodeling of the tumor microenvironment (TME), and therapy resistance [4–6]. Current treatment relies on a multimodal approach combining maximal surgical resection, radiotherapy, and temozolomide chemotherapy [7]. Despite the aggressive regimen, prognosis remains poor, with an average survival of only 15 months [1, 2, 8]. These limitations underscore the urgent need to identify molecular targets that could improve patient outcomes. Accordingly, extensive efforts have sought to identify key regulators of tumor progression *in vitro* and *in vivo* to elucidate pathways driving GBM aggressiveness. In this study, we aim to investigate the cellular and transcriptional changes within GBM tumors and their association with the TME to define mechanisms that contribute to tumor progression.

GBM exhibits extensive inter- and intratumoral heterogeneity, giving rise to tumor cell subpopulations with distinct proliferative capacities, metabolic states, and therapeutic responses [9]. A defining feature of these subpopulations can transition between invasive and proliferative phenotypes, highlighting proliferative signaling pathways as potential therapeutic targets [9, 10]. In many GBM cases, abnormal activation of the CDK-Rb-E2F axis drives unchecked G1/S transition, leading to uncontrolled proliferation. Additionally, activation of the PI3K−AKT pathway through PTEN loss or EGFR amplification promotes survival, growth, and transcription of mitotic regulators [9]. GBM tumor cells commonly overexpress cell cycle regulators including *CDK1*, *CCND1*, *CCNA2*, *PBK*, *PLK1*, *BUB1*, *CHEK1*, and *BIRC5* (survivin), sustaining high mitotic activity while suppressing apoptotic signaling [11, 12]. The coexistence of multiple proliferative and survival programs within individual tumors and across patient populations underscores the importance of identifying therapeutic targets that can effectively reduce cell cycle progression and proliferation.

During GBM progression, the extracellular matrix (ECM) within and surrounding the tumor undergoes extensive remodeling, generating a pathological TME that actively promotes tumor aggressiveness [13–16]. Non-neoplastic brain tissue is composed of non-fibrillar components such as hyaluronic acid, proteoglycans, and tenascins, which form a flexible scaffold. In contrast, the GBM-associated ECM is remolded into a stiff pathological matrix enriched in collagens and glycoproteins, including fibronectin, which are minimally expressed in normal brain tissue. Differential expression of ECM components contributes to GBM progression by promoting tumor cell proliferation and survival [4]. A defining feature of GBM is increased ECM stiffness within the tumor region, which activates mechanosensitive signaling pathways that accelerate tumor cell migration, proliferation, and therapeutic resistance [17]. While non-neoplastic brain tissue typically exhibits stiffness values of 0.2–1.2 kPa (kilopascal), GBM tumors can reach levels as high as 35 kPa [17]. These values illustrate the dramatic difference in tissue stiffness between GBM and normal brain, emphasizing the mechanosensitive behavior of tumor cells. Pathological ECM stiffness regulates tumor cell behavior through mechanotransduction, a process during which mechanical cues from the ECM are sensed by integrins on the cell surface and transmitted into intracellular signaling cascades [18]. Previous studies have shown that glioma cells enhance mechanosensing by upregulating ECM proteins, adhesion molecules, and cytoskeletal regulators, which facilitate stiffness sensing and cellular responses [19, 20]. The dynamic and reciprocal interactions between GBM cells and the ECM therefore create a feedforward loop that promotes tumor progression, highlighting the ECM as both a mediator of disease and a prospective therapeutic target in GBM [21].

Among the molecular drivers of GBM, survivin, a member of the inhibitor of apoptosis protein family encoded by baculoviral inhibitor of apoptosis repeat-containing 5 (BIRC5), has emerged as a critical regulator of tumor progression [22, 23]. Survivin has dual functions: it regulates the cell cycle by facilitating mitotic spindle assembly and chromosomal segregation, and it suppresses apoptosis through caspase inhibition. Although largely absent from most differentiated adult tissues, survivin is highly upregulated in various human cancers, where its expression correlates with increased tumor cell proliferation and migration, therapeutic resistance, and poor clinical outcomes [24]. In glioma, survivin levels positively correlate with pathologic grade, with WHO Grade IV glioblastoma exhibiting the highest expression [12], and elevated survivin levels predict therapeutic resistance [25]. Despite its established role as a clinical biomarker and regulator of cell cycle progression, the potential involvement of survivin in ECM regulation and stiffness-mediated mechanotransduction remain unclear.

In this study, we investigate the regulatory mechanisms underlying aberrant cellular behaviors in GBM, with a focus on dysregulated proliferation and ECM production driven by pathological tissue stiffness. Using an integrated approach combining bioinformatic analysis, *in vivo* tissue staining, and *in vitro* assays, we identified survivin as a critical mediator of stiffness-induced signaling, modulating both cell cycle progression and ECM synthesis. Our findings reveal a coordinated interplay between ECM stiffness and survivin expression that promotes a hyperproliferative and ECM-producing phenotype characteristic of GBM. By elucidating how mechanical cues converge with oncogenic cellular behaviors, this study advances our understanding of the molecular drivers of GBM and highlights the potential of targeting mechanosensitive pathways as a therapeutic strategy.

## RESULTS

### GBM transcriptome reveals altered proliferation, ECM remodeling, and neuronal function

Whole-transcriptome analysis was performed using the publicly available Gene Expression Omnibus (GEO) dataset GSE10878, generated by Tayrac *et al*., to identify transcriptomic changes between 19 histologically defined GBM tumors and 4 non-neoplastic brain tissues [26]. The distribution and relationships among samples are shown in a UMAP plot (Fig. 1A). Samples clustered distinctly according to GBM and non-neoplastic groups, demonstrating clear separation between the two conditions, while also revealing substantial heterogeneity among GBM patient samples (Fig. 1A). From 8887 genes in the dataset, filtering based on significance thresholds (q values ≤ 0.05, log_2_(fold change) > 0.75) identified 1858 differentially expressed genes (DEGs), including 593 upregulated and 1265 downregulated genes in GBM tumor samples. A volcano plot displayed the distribution of DEGs relative to total number of genes by plotting −log_10_(q value) against log_2_(fold change) for each gene, with red denoting significantly upregulated genes and blue denoting significantly downregulated genes (Fig. 1B).

**Figure 1.**
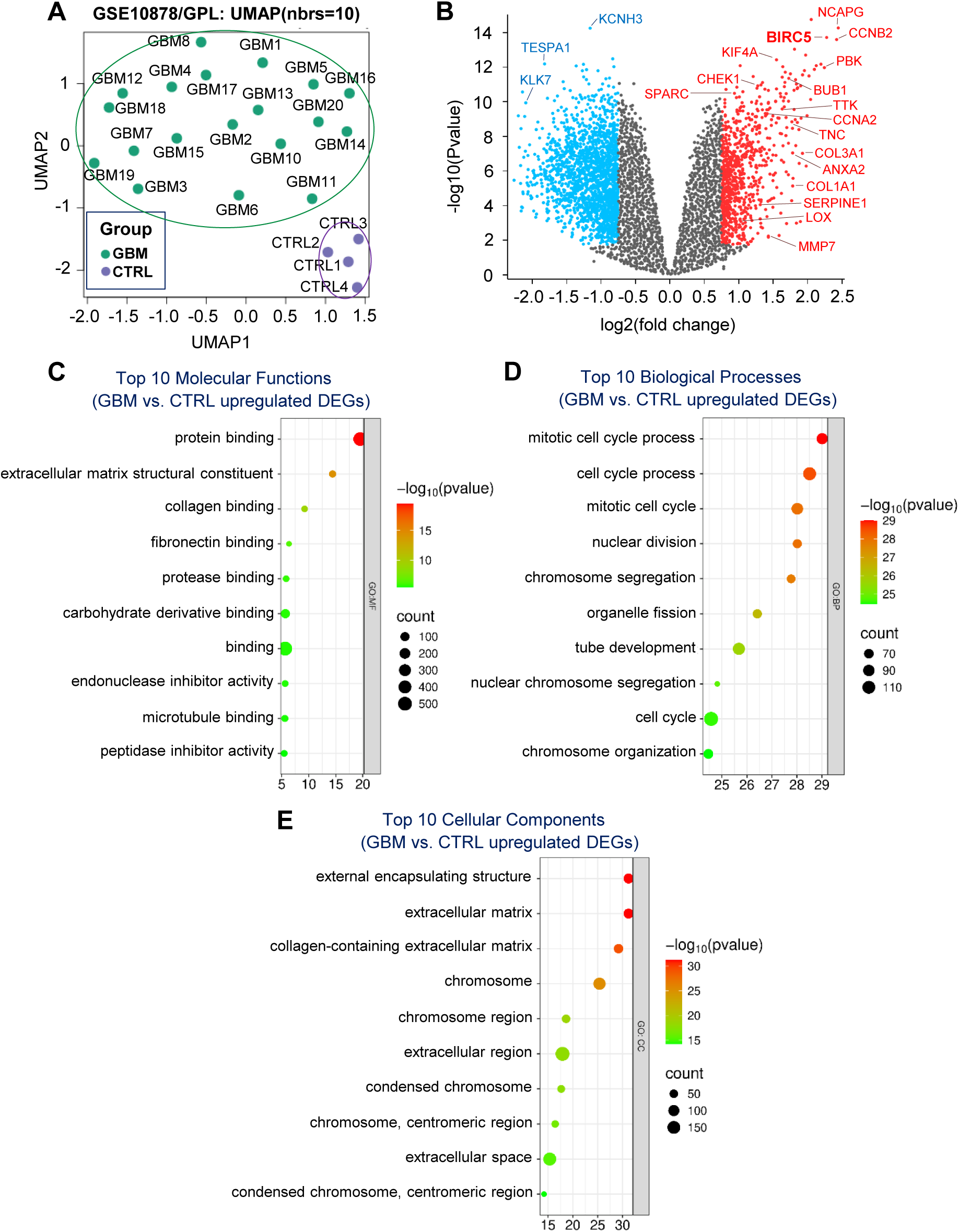
Transcriptomic analysis reveals DEGs in GBM tumors. DEGs in human GBM tumors versus non-neoplastic brain tissue were identified using thresholds (q ≤ 0.05; log2FC ≥ 0.75). Principle component analysis shows individual sample variance (**A**). Volcano plot depicts significantly upregulated (red) or downregulated (blue) genes relative to all identified genes (**B**). Bubble plots show the top 10 enriched molecular functions (**C**), biological processes (**D**), and cellular components (**E**) for upregulated DEGs.

To explore the biological significance of these transcriptional changes in GBM, functional enrichment analysis of upregulated and downregulated DEGs was performed using g:Profiler. Enriched molecular functions (MFs) among upregulated DEGs were primarily associated with ECM structural integrity and protein interactions, particularly “extracellular matrix structural constituent,” “collagen binding,” and “fibronectin binding” (Fig. 1C), reflecting reinforcement of ECM organization and remodeling in GBM pathology. In contrast, downregulated DEGs for MFs were enriched for ion transport–related molecular functions, including “monoatomic ion channel activity,” “gated channel activity,” and “passive transmembrane transporter activity,” revealing impaired ion transport and electrochemical regulation in GBM compared with non-neoplastic brain tissue (Supplementary Fig. 1A).

Analysis of enriched biological processes (BPs) further revealed that upregulated DEGs were dominated by proliferation and cell cycle-related terms, including “mitotic cell cycle process,” “nuclear division,” “chromosome segregation,” “tube development,” and “vasculature development” (Fig. 1D), collectively indicating increased proliferative capacity and tissue remodeling in GBM. In contrast, downregulated DEGs were enriched for neuronal signaling pathways, including “trans-synaptic signaling,” and “chemical synaptic transmission,” as well as processes critical for nervous system development and neuron differentiation (Supplementary Fig. 1B), collectively indicating broad disruption of neuronal communication and developmental programs in GBM tissue.

Upregulated DEGs were enriched in cellular components (CCs) related to both the ECM and chromosomes, including “extracellular matrix,” “collagen-containing extracellular matrix,” “extracellular region,” “chromosome,” “chromosomal region,” and “condensed chromosome,” indicating enrichment of ECM remodeling and cell cycle–associated nuclear components (Fig. 1e). In contrast, downregulated DEGs were enriched in neuronal components, including “synapse,” “synaptic junction,” “postsynapse,” “presynapse,” “neuron projection,” and “axon” (Supplementary Fig. 1C), indicating a marked reduction of genes maintaining neuronal connectivity and signaling structures within GBM.

Finally, clustering analysis of upregulated DEGs revealed three major interconnected clusters associated with ECM and mitotic cell cycle processes, along with a smaller cluster related to Hox gene activation, highlighting coordinated regulation of proliferation and matrix remodeling during GBM progression (Supplementary Fig. 2). In contrast, downregulated DEGs formed two distinct clusters associated with “GPCR ligand binding” and “synapse”, indicating impaired neuronal signaling during GBM progression (Supplementary Fig. 3).

### Heatmap and network analyses identify key regulators of cell cycle and proliferation in GBM

To visualize gene expression patterns related to cell cycle progression and proliferation, we analyzed the expression of cell cycle- and proliferation-associated genes in GBM tissues compared with non-neoplastic brain tissues by annotating DEGs using the KEGG *Cell Cycle* gene list. Heatmap analysis revealed broad upregulation of genes involved in cell cycle regulation and mitotic progression, including *CCNB2*, *CCNA2*, *BIRC5*, *PBK*, *TTK*, *BUB1B*, and *PLK1*, along with reduced expression of the cell cycle inhibitor *CDKN2D* (Fig. 2A). Several checkpoint regulators, such as *CHEK1*, *WEE1*, and *TP53*, were also dysregulated, indicating disruption of cell cycle control.

**Figure 2.**
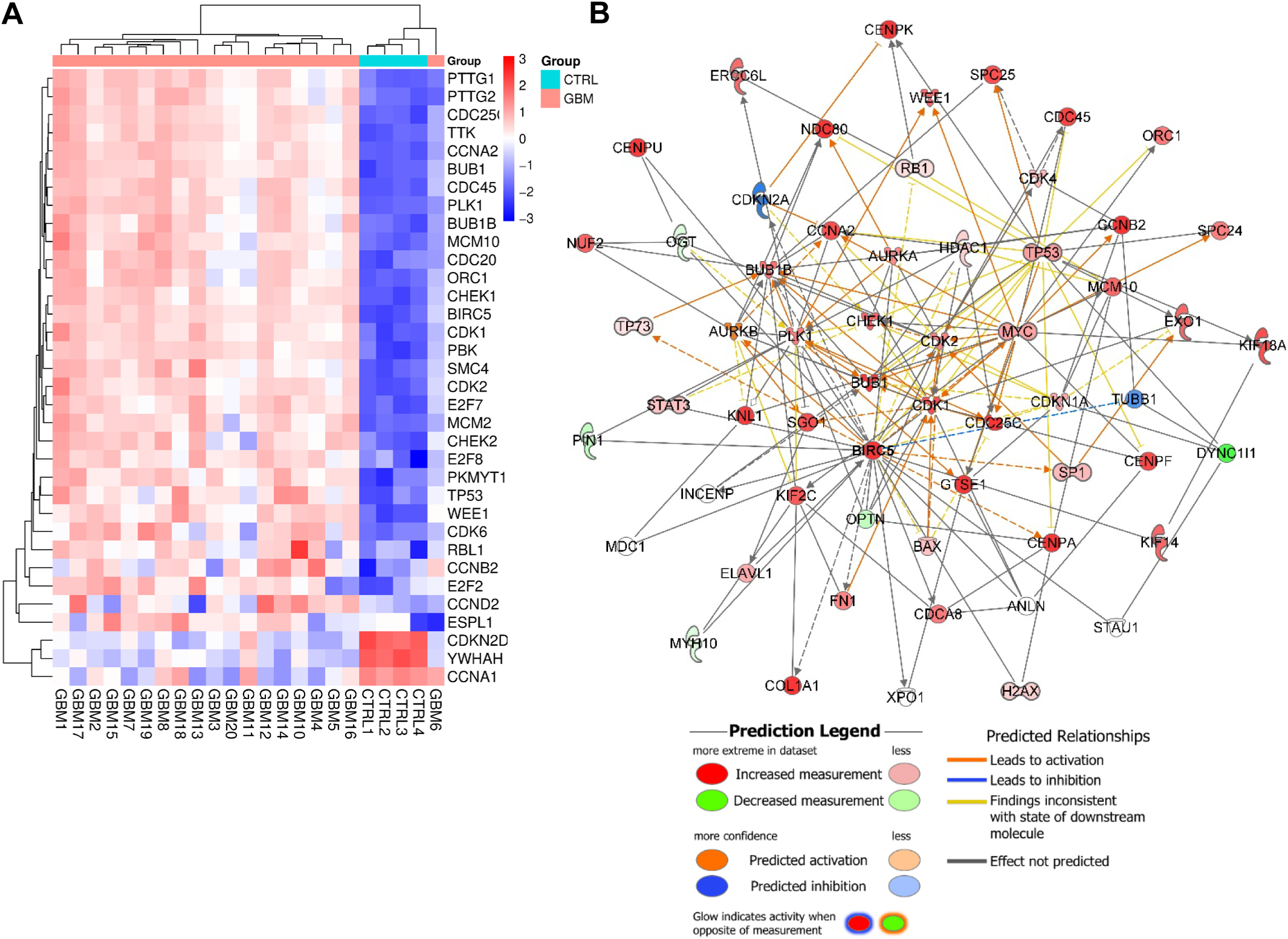
GBM tumors show enrichment of cell cycle genes and survivin. DEGs annotated using the KEGG *Cell Cycle* gene list are visualized with SRplot in a heatmap (**A**). Mechanistic network predicting BIRC5 regulation of cell cycle DEGs identified from IPA’s Canonical pathway analysis (Z-score=2.124) (**B**).

Survivin is commonly upregulated in cancer, where it promotes cell cycle progression, proliferation, and therapeutic resistance. In our dataset, *BIRC5* (survivin) ranked as the third most upregulated cell cycle- and proliferation-associated gene (Supplementary Table 1). Network analysis using Ingenuity Pathway Analysis (IPA) revealed that *BIRC5* is interconnected with multiple upregulated DEGs driving cell cycle progression and proliferation (Z-score = 2.124; significant if ≥ 2), including *CCNA2*, *CCNB2*, *CDK1*, *CDK2*, *CHEK1*, *BUB1B*, and *PLK1* (Fig. 2B). These findings identify survivin as a potential upstream regulator of cell cycle-associated genes in GBM.

### Matrisome profiling reveals distinct ECM signatures in GBM

To characterize the ECM profile in GBM, we performed a targeted analysis of matrisome genes, comprising core ECM genes (collagens, proteoglycans, and glycoproteins) and ECM-associated genes (ECM-affiliated proteins, ECM regulators, and secreted factors) [27, 28], enriched in tumors using the Matrisome Database [29]. Upregulated and downregulated DEGs were annotated according to the Matrisome Project classification, assigning each gene to a matrisome division (core vs. associated) and a functional category (e.g., collagens, proteoglycans, glycoproteins, ECM-affiliated proteins, ECM regulators, secreted factors). Heatmaps of matrisome gene expression (Fig. 3A−B; Supplementary Fig. 4) revealed heterogenous but distinct ECM profiles between non-neoplastic and GBM samples. GBM tumors exhibited broad upregulation of core matrisome genes, including multiple collagens (*COL1A1*, *COL3A1*, *COL4A1*, *COL4A2*), glycoproteins (*FN1*, *SPARC*, *TNC*) and proteoglycans (*VCAN*, *BGN*). In parallel, several matrisome-associated genes were elevated in GBM, particularly ECM regulators (*MMP2*, *MMP7*, *LOX*), ECM-affiliated proteins (*LGALS1*, *ANXA2*, *MFGE8*), and secreted factors (*SERPINE1*, *IGFBP3*). These patterns suggest coordinated activation of ECM synthesis and remodeling pathways, indicating that both structural and regulatory components of the ECM are extensively reprogrammed in GBM.

**Figure 3.**
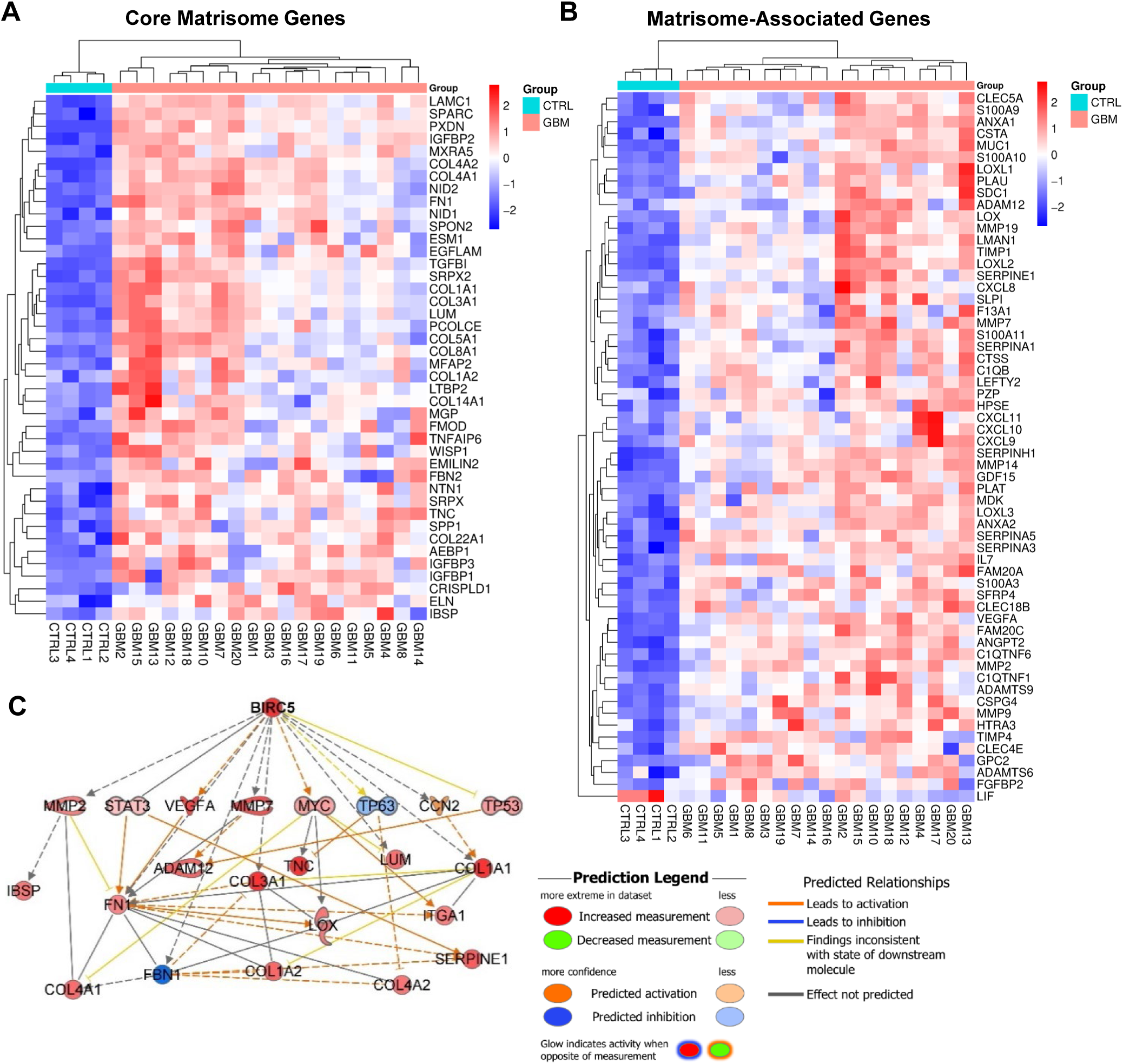
Matrisome profiling reveals distinct ECM signatures in GBM tumors. Upregulated DEGs were annotated via the Matrisome database and classified into core matrisome (**A**) and matrisome-associated (**B**) categories, visualized per sample in a heatmap. Mechanistic network predicting BIRC5 regulation of DEGs from ‘Extracellular matrix organization’ identified from IPA’s Canonical pathway analysis (Z-score=2.124) (**C**).

To explore survivin’s association with ECM remodeling, we identified DEGs interacting with *BIRC5* and linked to ECM regulation (Z-score = 2.124). Network analysis revealed predicted interactions between *BIRC5* and key ECM components, including collagens (*COL1A1*, *COL3A1*, *COL4A1*, *COL4A2*), fibronectin (*FN1*), and matrix remodeling enzymes (*MMP2*, *MMP7*, *LOX*) (Fig. 3C). As shown in the network, survivin acts as an upstream regulator influencing genes involved in ECM. These findings imply that survivin contributes to pathological ECM remodeling and may facilitate the invasive and proliferative behavior characteristic of GBM.

### Survivin and ECM proteins are upregulated in GBM tumors *in vivo*

To evaluate survivin expression and its spatial association with ECM components in GBM, immunostaining analyses were performed in both U87-implanted nude rats and human GBM tissue sections. For *in vivo* tumor engraftment, 100,000 U87 human GBM cells in 5μL of PBS were implanted into the cortex of T-cell deficient nude rats at a depth of 7 mm. Tumor progression was monitored daily by weight, neurological assessments, and pre-euthanasia MRI. Following confirmation of tumor growth, U87-implanted rats were euthanized and tissues were formalin-fixed alongside saline-injected and healthy controls. STEM121 staining, which detects engrafted human U87 cells after transplantation, confirmed well-defined tumor boundaries in all U87-implanted rats, indicating successful engraftment of U87-implanted cells (Fig. 4A). Immunohistochemical (IHC) analysis showed strong nuclear survivin staining in tumor regions of U87-implanted rat brains compared with non-tumor areas, whereas saline-injected and healthy controls displayed markedly lower, diffuse expression (Fig. 4B−C; Supplementary Fig. 6). Similarly, human GBM sections exhibited significant survivin expression within tumor regions, while non-neoplastic brain tissue sections showed markedly lower staining (Fig. 4F−H; Supplementary Fig. 7). Tumor-associated ECM remodeling accompanied survivin upregulation in both rat and human GBM samples. Collagen 1a1 (Col1a1) expression was elevated in tumor regions of U87-implanted rats relative to adjacent non-tumor tissue (Fig. 4D−E). Consistently, human GBM sections showed strong Col1a1 staining, particularly within tumor-associated vasculature (Fig. 4I). Together, these results support the transcriptomic findings by demonstrating increased survivin expression and ECM protein accumulation in GBM tumor regions *in vivo*.

**Figure 4.**
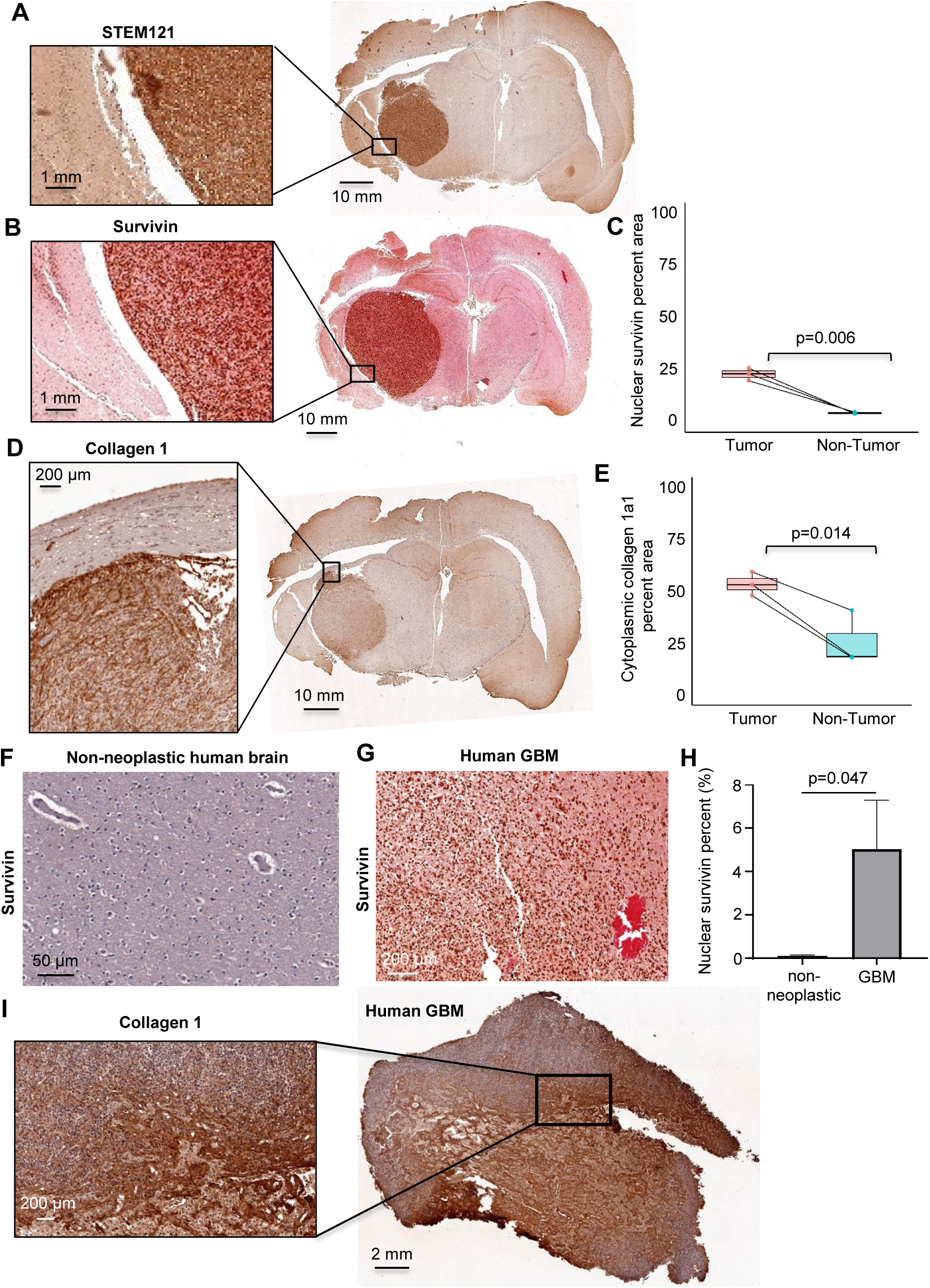
U87-implanted nude rats and human GBM tissue show survivin and collagen upregulation. U87 GBM cells (100,000) were implanted into the cortex of T-cell deficient nude rats (*n=3*). Tumor progression was monitored and confirmed by MRI. Brain sections were STEM121 stained (**A**) to confirm tumor margins. IHC shows strong nuclear survivin in tumor regions versus peri-tumor areas (**B−C**) and increased collagen staining (**D−E**). Human GBM resections (*n=3*) show strong nuclear survivin (**G**–**H**) compared to non-neoplastic controls (*n=3*) (**F**). Human GBM resections displayed significant collagen 1 (**I**) staining within the tumor borders. Whole-slide imaging at 40x.

### Pathological ECM stiffness upregulates survivin, cyclins, and ECM proteins in GBM cells

Our bioinformatic and *in vivo* analyses revealed upregulation of survivin together with cell cycle-and ECM-related genes in GBM tumors. To determine whether these changes could be recapitulated *in vitro*, U87 and A1207 human GBM cells were cultured on fibronectin-infused polyacrylamide hydrogels with defined stiffness levels representing normal brain tissue (low; ∼0.90 kPa), early-stage tumor stiffening (medium; ∼3.64 kPa), and advanced tumor stiffness (high; ∼24.36 kPa) [17, 30]. For these studies, GBM cells were synchronized in G0 by serum starvation and plated on the hydrogels in medium containing 10% fetal bovine serum (FBS). After 24 hours, cell lysates were collected for immunoblotting. Survivin expression was markedly upregulated in both U87 (Fig. 5A−B) and A1207 (Supplementary Fig. 8A−B) cells cultured on high-stiffness hydrogels compared with those on low- and medium-stiffness conditions. GBM cells grown on high-stiffness hydrogels also exhibited increased expression of the cell-cycle related proteins cyclin D1 (Fig. 5C−D; Supplementary Fig. 8C−D) and cyclin A (Fig. 5C, E; Supplementary Fig. 8E−F), as well as the ECM-associated proteins collagen (Fig. 5F−G) and lysyl oxidase (Fig. 5H−I; Supplementary Fig. 8G−H). Although medium stiffness showed a trend toward increased protein levels, these changes were not statistically significant compared with low stiffness. Because both cell lines showed similar expression patterns and no significant differences were observed between low- and medium-stiffness conditions, subsequent experiments were performed using U87 cells cultured on low- and high-stiffness hydrogels. Taken together, these results demonstrate that stiffness-tunable hydrogels recapitulate the expression patterns of survivin, cell cycle-related, and ECM-associated genes and proteins observed in GBM tumors.

**Figure 5.**
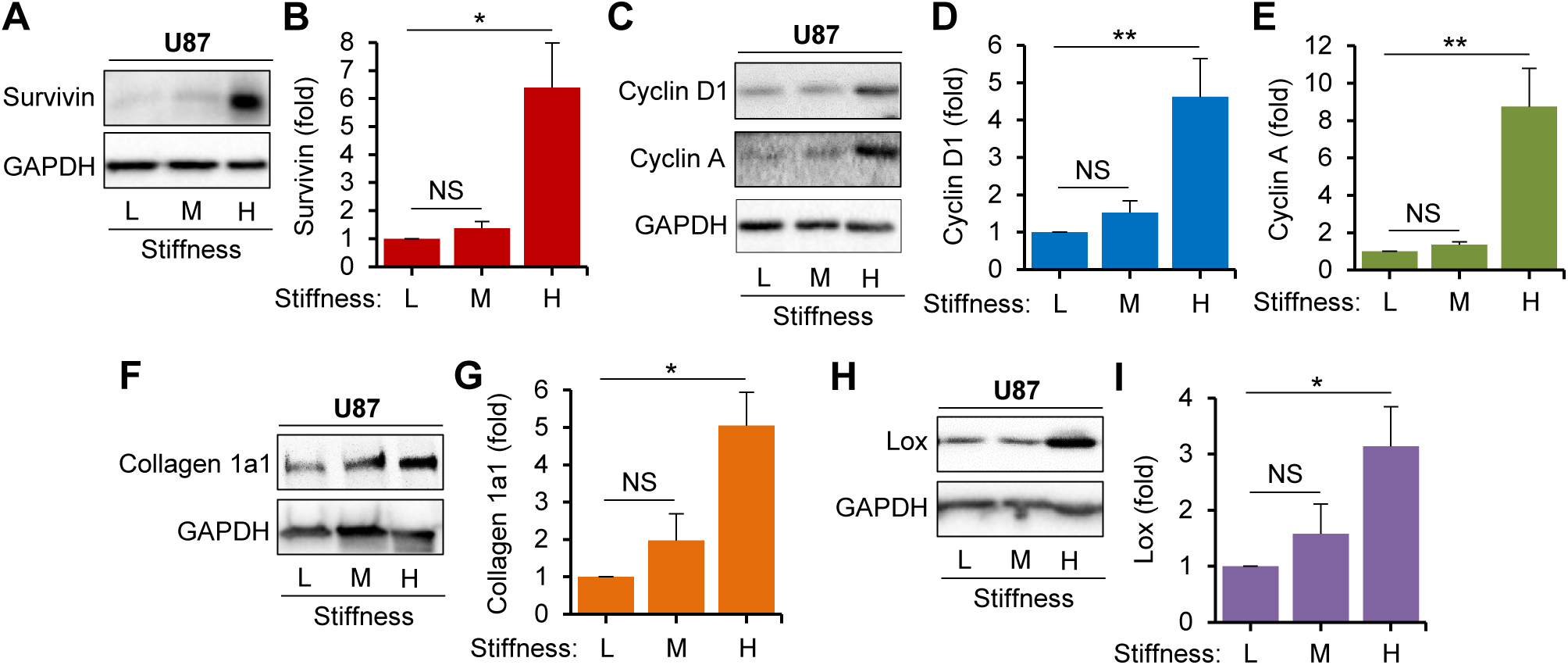
Stiffness induces expression of survivin, cell cycle- and ECM-associated proteins in U87 cells. Western blot analysis shows stiffness-induced changes in survivin (**A−B**), cyclin D1 (**C−D)**, cyclin A (**C, E**), collagen 1a1 (**F−G**), and lysyl oxidase (Lox) (**H−I**) after 24 hours on low (L), medium (M), and high (H) stiffness hydrogels. Protein levels were normalized to GAPDH, displayed relative to low stiffness. Representative blot images are shown next to quantification. *n=3-6;* Significance: *\*p < 0.05*, *\*\*p < 0.01*, and *NS*: not significant.

### Survivin regulates stiffness-induced cell cycle progression in GBM cells

To explore the role of survivin in stiffness-induced cell cycle progression, U87 cells were cultured on low- and high-stiffness hydrogels and treated with sepantronium bromide (YM155), a small-molecule inhibitor that suppresses survivin transcription by blocking SP1 binding to its promoter [31]. Treatment with 0.1 μM YM155 for 24 hours effectively reduced survivin protein levels (Fig. 6A–B) and decreased protein expression of cyclin D1 (Fig. 6C−D) and cyclin A (Fig. 6C, E), indicating that survivin is required for stiffness-induced cell cycling. GBM cells were also transfected with survivin-targeting siRNA or control siRNA (200 nM) for 48 hours prior to seeding on low- and high-stiffness hydrogels. After 24-hour incubation, total cell lysates were collected for immunoblotting. Survivin knockdown (Fig. 6F−G) attenuated stiffness-induced upregulation of cyclin D1 (Fig. 6F, H) and cyclin A (Fig. 6F, I) compared with control siRNA. Finally, EdU incorporation assays demonstrated reduced S-phase entry in U87 cells treated with YM155 (Supplementary Fig. 9A) or survivin siRNA (Supplementary Fig. 9B), indicating that survivin contributes to stiffness-induced proliferation in GBM cells.

**Figure 6.**
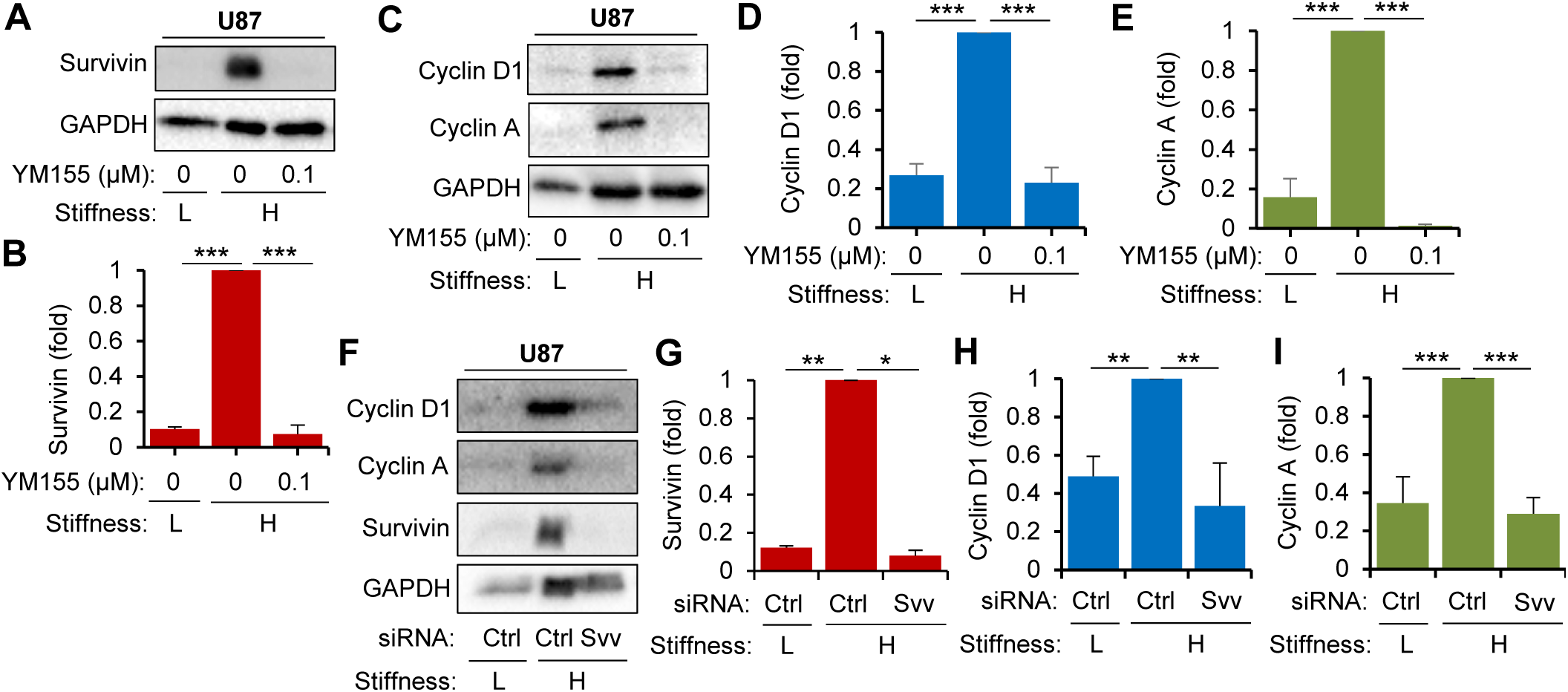
Survivin inhibition reduces stiffness-induced proliferation in U87 cells. Serum-starved cells plated on low and high stiffness hydrogels were treated with DMSO (Ctrl) or 0.1𝜇M YM155 for 24 hours. Western blot analysis shows reduced survivin (**A−B**), cyclin D1 (**C−D**), and cyclin A (**C, E**) expression on high stiffness hydrogels in U87 cells treated with YM155. For siRNA knockdown, U87 cells were transfected with 200 nM control or survivin siRNA. Western blot analysis confirms survivin knockdown (**F−G**) and effects on Cyclin D1 (**F**, **H**) and Cyclin A (**F**, **I**) expression. *n=3-6*; Significance: *\*p < 0.05*, *\*\*p < 0.01*, *\*\*\*p < 0.001*, and *NS*: not significant.

### Survivin regulates stiffness-induced expression of matrisome-related proteins

To confirm the role of survivin in ECM regulation predicted by IPA analysis, U87 cells were cultured on low- and high-stiffness hydrogels and treated with YM155. Survivin inhibition significantly reduced collagen 1 expression (Fig. 7A−B), indicating that survivin promotes pathological ECM production. Furthermore, survivin inhibition attenuated stiffness-induced Lox expression (Fig. 7C−D), suggesting survivin is required for stiffness-mediated ECM remodeling. Consistent with these findings, siRNA-mediated knockdown of survivin reduced stiffness-dependent expression of collagen 1 (Fig. 7E–F) and Lox (Fig. 7E, G), confirming survivin is required for ECM production and remodeling.

**Figure 7.**
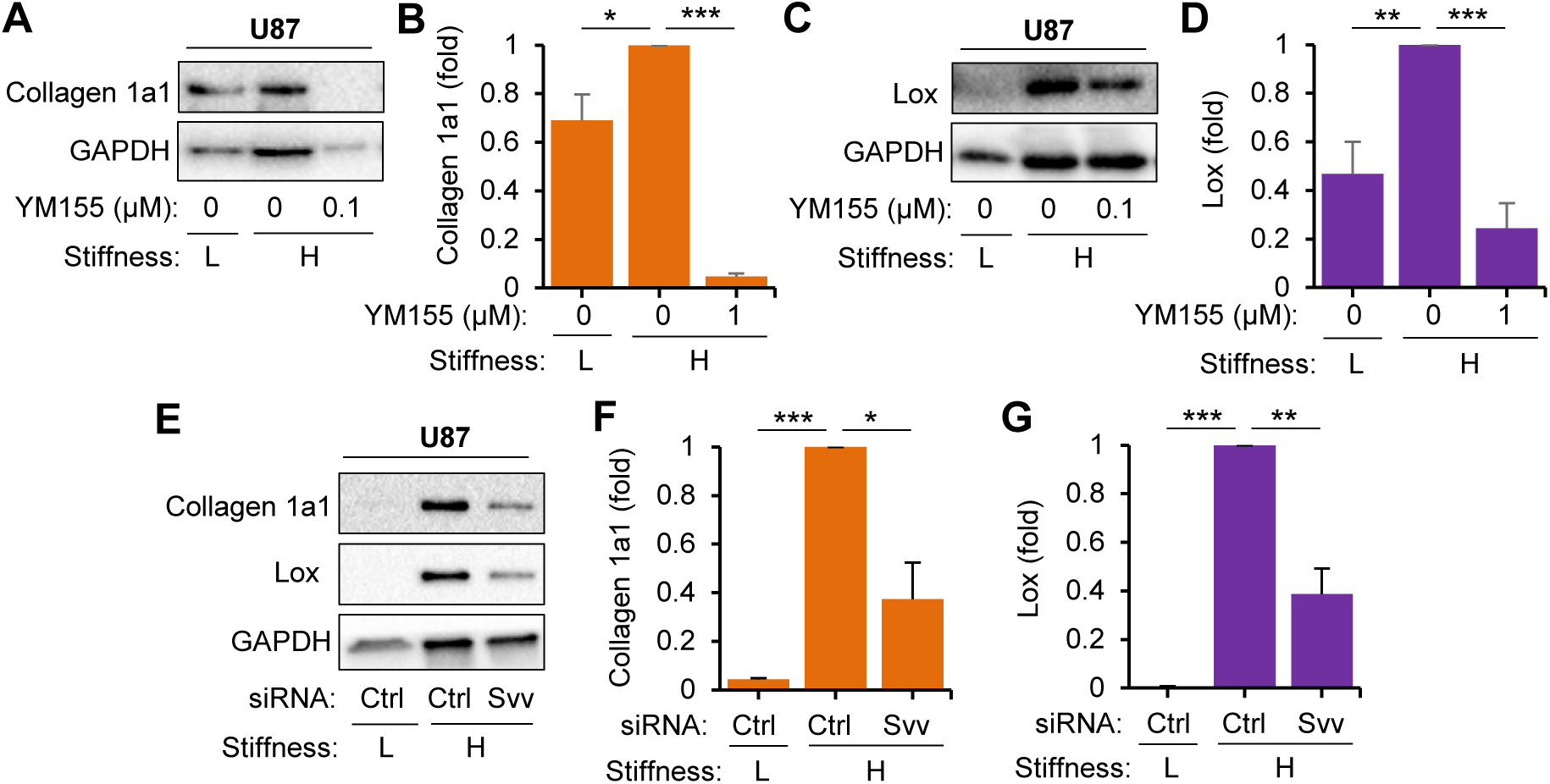
Survivin inhibition reduces stiffness-induced ECM synthesis in U87 Cells. Cells plated on low and high stiffness hydrogels were treated with DMSO or 0.1𝜇M YM155 for 24 hours. Western blots and representative images show changes in collagen 1a1 (**A−B**) and LOX (**C-D**). For siRNA knockdown, U87 cells were transfected with 200 nM control or survivin-targeting siRNA. Western blot analysis confirmed that survivin knockdown reduces stiffness-dependent expression of collagen 1a1 (**E−F**) and Lox (**E, G**). *n=4-6*; Significance: *\*p < 0.05*, *\*\*p < 0.01*, *\*\*\*p < 0.001*, and *NS*: not significant.

Collectively, these results indicate that survivin contributes to GBM progression through both its canonical regulation of the cell cycle and its novel role in ECM remodeling.

## DISCUSSION

Progressive stiffening of the ECM within the TME is a defining feature of GBM that activates mechanosensitive signaling pathways promoting tumor growth, invasion, and therapeutic resistance [13, 14, 17, 32–34]. As tumors develop, excessive ECM deposition and active remodeling further increase tissue stiffness, reinforcing malignant progression by enhancing tumor cell proliferation and migration [17] [35] [36]. Although the causality between TME stiffness and malignant cellular behavior remains under investigation, accumulating evidence indicates that mechanical and molecular changes co-develop to sustain tumorigenesis [37, 38]. However, the molecular mechanisms by which GBM cells sense mechanical stiffness and convert these cues into pro-tumorigenic signaling remain incompletely defined. In this study, we identified survivin as a mechanosensitive regulator that links ECM stiffness to GBM cell proliferation and ECM production, suggesting that survivin coordinates tumor growth and ECM remodeling in a stiff microenvironment.

Transcriptomic comparison of histologically defined GBM tumors and non-neoplastic brain tissue revealed coordinated activation of proliferative and ECM-remodeling programs, coupled with suppression of neuronal signaling genes. Broad upregulation of interphase and mitotic cell cycle regulators, together with altered checkpoint components, reflects an enhanced proliferative state consistent with the rapid expansion characteristic of GBM. Within this network, survivin emerged as a central node connecting cyclins, CDKs, mitotic regulators, supporting its role in cell cycle progression in GBM. These findings are consistent with previous studies demonstrating that survivin expression correlates with tumor grade, proliferation, and therapeutic resistance in gliomas [39, 40]. Matrisome profiling further revealed upregulation of both core ECM proteins, including collagens, fibronectin, as well as ECM-regulatory proteins such as matrix metalloproteinases and lysyl oxidase. These transcriptional patterns indicate that GBM cells actively remodel the extracellular environment rather than merely responding to it, contributing to the emergence of a mechanically stiff tumor niche that facilitates tumor expansion and invasion [41, 42].

A previous study in GBM patients reported significant upregulation and nuclear localization of survivin, promoting cell division and proliferation [43]. Consistent with both our transcriptomic analysis and this prior report, immunostaining of U87-implanted rat brains and human GBM specimens demonstrated strong nuclear survivin expression within tumor regions, coinciding with areas of elevated ECM deposition, indicating that survivin-expressing tumor cells reside within matrix-rich microenvironments. These observations support the transcriptomic data and the previous report, suggesting that survivin is associated with both proliferative activity and ECM remodeling within GBM tumors *in vivo*.

The mechanoregulation of ECM dynamics by survivin is particularly relevant given the pronounced stiffness differences between healthy brain tissue and GBM tumors and the established role ECM stiffness in promoting tumor progression. Stiffness-tunable hydrogels provide a physiologically relevant *in vitro* system for modeling tumor microenvironments and examining the effects of ECM mechanics on cellular behavior. Using this system, we observed that increasing hydrogel stiffness upregulated survivin expression, accompanied by elevated levels of key cell cycle regulators including cyclin D1 and cyclin A, corresponding to the G1 and S phases of the cell cycle. Pharmacologic inhibition of survivin with YM155 or siRNA-mediated knockdown significantly reduced stiffness-induced cyclin expression and S-phase entry, demonstrating that survivin functions as a mechanosensitive regulator of proliferation. In addition to its role in cell cycle regulation, survivin also influenced ECM-associated proteins, consistent with the upregulation of core matrisome and matrisome-affiliated genes observed in GBM tumors and in cells cultured on stiff hydrogels. Survivin inhibition attenuated stiffness-induced expression of these ECM components. These observations indicate that survivin contributes not only to proliferative signaling but also to active remodeling of the tumor microenvironment.

Together, these findings support a model in which ECM stiffening induces survivin expression, leading to coordinated activation of cell cycle progression and ECM production. This coupling may establish a feed-forward mechanism in which matrix stiffening promotes tumor proliferation while simultaneously reinforcing ECM remodeling that further increases tissue stiffness.

Although this study shows that survivin links mechanical cues to proliferative and ECM-remodeling responses in GBM cells, the mechanotransduction pathways underlying stiffness-dependent survivin regulation remain undefined. Future studies should investigate whether canonical mechanosensitive pathways, such as integrin-mediated cytoskeletal tension or YAP/TAZ signaling, contribute to survivin activation, and whether genes interconnected with survivin, such as BUB1, RB1, PLK1 (positive regulators) and CDKN2A, GLS2, PACSIN1, MAL2, KCNH3 (negative regulators), as well as ECM regulators like MMP2 and MMP7, mediate its proliferative and ECM-remodeling functions. Patient-derived GBM models and in vivo studies will further clarify survivin’s role in tumor progression and provide potential combinatorial therapeutic targets.

In summary, these findings identify survivin as a mechanosensitive regulator linking ECM stiffness to cell proliferation and ECM remodeling, providing insight into how mechanical cues contribute to GBM progression and suggesting survivin as a potential target for disrupting tumor growth within mechanically stiff tumor microenvironments.

## CONCLUSIONS

This study identifies survivin as a mechanosensitive regulator that links ECM stiffness to GBM cell proliferation and ECM remodeling. By coordinating cell cycle progression with ECM synthesis, survivin promotes a stiff, tumor-supportive microenvironment. Targeting survivin may suppress both proliferation and ECM remodeling, providing a potential therapeutic strategy to limit GBM aggressiveness.

## METHODS

### Differential Gene Expression Analysis

To investigate transcriptomic changes associated with GBM, a publicly available whole genome expression microarray dataset generated by de Tayrac et al. (GSE10878, 2008) was analyzed. This dataset compares 19 histologically defined GBM tumors with 4 non-neoplastic brain samples. Differential gene expression analysis was performed using Gene Expression Omnibus (GEO) web tool GEO2R with default settings. Statistical significance was determined using the Benjamini-Hochberg false discovery rate (B-H FDR). Differentially expressed genes (DEGs) were defined using significance cutoff of q-value ≤ 0.05 and absolute log_2_ fold change (log_2_FC) ≥ 0.75 (fold-change ≥ 1.68). Among the 8,887 genes in the dataset, filtering identified 1,858 DEGs, including 593 upregulated and 1,265 downregulated genes. UMAP and volcano plots were generated in GEO2R to visualize sample clustering and DEGs distribution, respectively.

### Functional Pathway Analysis

Upregulated and downregulated DEGs were analyzed separately using g:Profiler (https://biit.cs.ut.ee/gprofiler/gost) to perform Gene Ontology (GO) enrichment analysis. Biological Processes, Molecular Pathways, and Cellular Components were considered significant at a B-H FDR q-value ≤ 0.05. The top ten enriched terms in each category were visualized as bubble plots using SRplot.

### Network analysis

Network analysis was performed using Ingenuity Pathway Analysis (IPA). DEGs annotated in IPA as belonging to the canonical pathways “Cell Cycle Regulation” and “Extracellular matrix organization” were selected under canonical pathway analysis was utilized for further mechanistic predicted network analysis. Path Explorer feature under My Pathway was used to map out potential interactions between BIRC5, DEGs, and potential intermediate genes that were not previously revealed in the dataset for each network.

### Clustering Analysis

Protein-protein interaction (PPI) networks were generated using the STRING database. Upregulated and downregulated genes were analyzed separately, and nodes with a degree >2 were retained using Cytoscape filtering. Functionally distinct clusters within the PPI networks were identified using the STRING k-means clustering algorithm. Clusters were annotated based on the most enriched molecular function, biological process, and cellular component represented within each cluster.

### Matrisome Analysis

ECM-related transcriptional changes were further examined using the Matrisome AnalyzeR to classify DEGs into core matrisome categories (collagens, proteoglycans, and glycoproteins) and matrisome-associated categories (ECM-affiliated proteins, ECM regulators, and secreted factors). Expression patterns of annotated matrisome genes were visualized as heatmaps generated in SRplot, using hierarchical clustering with Euclidean distance and complete linkage.

### Cell Cycle Analysis

Cell cycle-related transcriptional changes were examined by annotating DEGs using the KEGG Cell Cycle gene list. Expression patterns of annotated cell cycle DEGs were visualized as heatmaps generated in SRplot using hierarchical clustering with Euclidean distance and complete linkage.

### Cell culture

U87 human GBM cells (RRID:CVCL_0022) and A1207 human GBM cells (RRID:CVCL_8481) were cultured in high-glucose Dulbecco’s modified Eagle’s medium (DMEM; 4.5 g/L glucose and L-glutamine without sodium pyruvate; 10-017-CV, Corning) supplemented with 1% HEPES buffer (25-060-CI, Corning), 1% penicillin–streptomycin (30-002-CI, Corning), and 10% fetal bovine serum (FBS; 35-010-CV, Corning). Cells were maintained at 37°C in a humidified incubator with 5% CO_2_. To synchronize cells at the quiescent (G_0_) phase of the cell cycle, cells were serum-starved for 48 hours in DMEM containing 1 mg/mL heat-inactivated, fatty acid-free bovine serum albumin (BSA). Synchronized cells were then detached with 0.05% trypsin-EDTA, centrifuged, resuspended, and seeded on soft or stiff hydrogels in DMEM containing 10% FBS for 24 hours prior to analysis.

### Preparation of fibronectin-infused polyacrylamide hydrogels

Polyacrylamide hydrogels with tunable stiffness were prepared as previously described [44–48]. Glass coverslips were treated with 0.1M NaOH solution (SS267, Fisher Scientific) for 3 minutes to increase surface reactivity, followed by coating with 3-(trimethoxysilyl)propyl methacrylate (440159, Sigma-Aldrich) to enable covalent bonding of fibronectin-infused polyacrylamide hydrogels. Soft (∼ 0.9 kPa; low-stiffness) and stiff (∼24 kPa; high-stiffness) hydrogels were prepared to mimic the stiffness of healthy brain tissue and glioblastoma-associated matrix, respectively. Hydrogels were generated by combining molecular biology-grade water, 1% bis-acrylamide, 40% acrylamide, and a tris-fibronectin solution containing acrylic acid N-Hydroxysuccunimide ester (A8060, Sigma-Aldrich) dissolved in DMSO (D2650, Sigma-Aldrich), followed by addition of 1 M tris-HCl (pH 8.4) with 0.075% fibronectin, 10% ammonium persulfate (A3678, Sigma-Aldrich), and TEMED (*N*,*N*,*N*′,*N*′-tetramethylethylenediamine; J63734.AC, Thermo Scientific) to initiate polymerization. The tris-fibronectin solution was incubated at 37°C for 1 hour prior to hydrogel preparation. Hydrogel solution (400 μL) was applied to 24x40-mm coverslips and 10 μL to 12-mm coverslips. Siliconized top coverslips were prepared using 20% Surfasil (TS42801, Thermo Scientific) in chloroform to prevent adhesion. The top coverslip was placed over the hydrogel solution on the bottom coverslips to ensure uniform spreading and removed after polymerization. Hydrogels were washed three times in 1x Dulbecco’s phosphate-buffered saline (DPBS; 21-031-CV, Corning) for 15 minutes and blocked in serum-free DMEM containing 1 mg/mL BSA for 1 hour before cell seeding. Coverslip sizes and cell-seeding density varied by application: 24x40-mm coverslips were seeded at 80-90% confluency for immunoblotting, whereas 12-mm coverslips were seeded to 50-60% confluency for immunostaining.

Hydrogel stiffness was quantified using atomic force microscopy (AFM). Hydrogels were indented using a circular symmetric AFM probe (QP-BioAC-Cl or qp-SCONT, NanoAndMore USA Corp) in contact mode on a NX12 AFM system (Park Systems) mounted on a Nikon ECLIPSE Ti2 inverted microscope. Ten measurements were obtained at different locations for each hydrogel, and the mean value was used to determine to hydrogel stiffness.

### Drug treatment and siRNA transfection

Serum-starved GBM cells were seeded on fibronectin-infused polyacrylamide hydrogels in DMEM containing 10% FBS and treated with 0.1 μM YM155 (a survivin inhibitor; 11490, Cayman Chemical) or DMSO (vehicle control) for 24−48 hours.

### siRNA transfection

GBM cells were transfected with 200 nM survivin siRNA (ID 121294; 5’-CCACUUCCAGGGUUUAUUCtt-3’ Ambion) or a negative control scramble siRNA (Silencer negative control siRNA; AM4636, Thermo Fisher Scientific) using Lipofectamine RNAiMAX (13778075, Thermo Fisher Scientific) according to the manufacturer’s instructions. Cells were transfected for 48 hours in Opti-MEM (31985-070, Gibco) and then seeded on hydrogels in medium containing 10% FBS for 24 hours before analysis.

### EdU incorporation

GBM cells cultured on hydrogels and treated with either 0.1 μM YM155 or DMSO were incubated with 20 μM EdU for 24 hours. Cells were then fixed in 3.7% formaldehyde for 1 hour. EdU incorporation was detected using the Click-iT EdU Alexa Fluor 488 Imaging Kit (C10337, Thermo Fisher Scientific) according to the manufacturer’s instructions. Cells on hydrogels were mounted on glass microscope slides with DAPI-containing mounting medium (P36962, Thermo Fisher Scientific). Five fields of view per hydrogel were analyzed to determine the percentage of EdU-positive cells among total DAPI-stained nuclei.

### Protein extraction and immunoblotting

GBM cells cultured on hydrogels for 24 hours were washed twice with cold 1X DPBS. Hydrogels were placed face-down on 200 μL of warm 5X sample buffer (250 mM tris-HCl, pH 6.8; 10% sodium dodecyl sulfate (SDS); 50% glycerol, 0.02% bromophenol blue; 10 mM 2-mercaptoethanol) for 2 minutes at room temperature. Cell lysates were denatured at 100°C for 5 minutes, fractionated on 8-12% SDS-PAGE gels, and transferred onto PVDF membranes using the Trans-Blot Turbo Transfer System (Bio-Rad). Membranes were blocked for 1 hour in either 6% milk or 5% BSA in 1X TBST, followed by incubation with primary antibodies for 2 hours at room temperature and overnight at 4°C: survivin (NB500-201, Novus Biologicals; 1:500), cyclin A (sc-271682, Santa Cruz Biotechnology; 1:200), cyclin D1 (sc-20044, Santa Cruz Biotechnology; 1:200), lysyl oxidase (NB100-2530, Novus Biologicals; 1:500), collagen 1a1 (C2456, Sigma-Aldrich; 1:200), and GAPDH (10494-1-AP, Proteintech; 1:4000). Membranes were washed three times in TBST for 15 minutes each and incubated with HRP-conjugated secondary antibodies (SA00001-2 or SA00001-1, Proteintech) for 1 hour. Signals were detected using Clarity Western ECL Substrate (170-5061, Bio-Rad) or Clarity Max Western ECL Substrate (1705062, Bio-Rad). Bands were imaged and quantified using Image Lab Software with normalization to GAPDH.

### Tumor Implantation and Controls

Human U87 GBM cells (100,000 cells in 5μL) were stereotactically implanted into the cortex of T-cell-deficient nude rats (NIH-Foxn1^rnu^; *n=3*), as previously described [49]. Rats were monitored daily for pain and neurological symptoms using the Grimace scale and Menzies scoring system. Rats exhibiting neurological deficits, inability to independently feed or ambulate, or > 15% weight loss underwent pre-euthanasia MRI (day 15) to confirm tumor development using a 9.4 T Bruker preclinical MRI system (Bruker, Billerica, MA, USA). T1-weighted images were acquired with and without contrast. Rats went euthanized on day 17 using 5% isoflurane anesthesia followed by decapitation and perfusion fixation with formalin. Healthy nude rat controls (*n=6*) and saline-injected controls (5 μL; *n=6*) were euthanized using the same procedure. All animal procedures were approved by the Institutional Animal Care and Use Committee (IACUC) at the University at Buffalo (#PROTO202200068).

### Human GBM and non-GBM tissue acquisition

Human GBM tissue specimens and associated data were obtained with approval from the Institutional Review Board (IRB #00009245) at the University at Buffalo. Non-GBM brain tissue specimens and data were obtained from the NIH NeuroBioBank.

### H&E Staining

Histology analysis of GBM tissue sections was performed at the Jacobs School of Medicine and Biomedical Sciences’ Histology Core. Formalin-fixed, paraffin-embedded (FFPE) tissue sections (4 µm in thickness) were deparaffinized in xylene, rehydrated through graded ethanol, stained with hematoxylin, differentiated with acetic acid, dehydrated with ascending graded ethanol, stained with eosin, cleared in xylene, and cover-slipped. Slides were scanned at 40X using the Aperio VERSA Brightfield, Fluorescence & FISH Digital Pathology Scanner (Leica Biosystems, Deer Park, IL, US).

### Immunostaining

Immunohistochemistry (IHC) was performed on 4-µm FFPE tissue sections using an IHC kit (AB64261, Abcam) according to the manufacturer’s instructions. Engraftment of human U87 cells into the nude rat brains was confirmed using STEM121 (Y40410, Takara Bio; 1:200). IHC staining for survivin (2808, Cell Signaling Technology; 1:1000) and collagen1a1 (NBP1-300054, Novus Biologicals; 1:500) was performed to assess their expression in U87-implanted nude rats. To examine survivin and ECM protein expression in human GBM tissues, FFPE human GBM sections (4-5 μm thick; *n=3*) obtained from surgical resections (University at Buffalo Neurosurgery, Buffalo General Hospital, Buffalo, NY) were analyzed by IHC. Survivin staining was performed on these human tissue sections. Brightfield slides were scanned at 40X using the Aperio VERSA Brightfield, Fluorescence & FISH Digital Pathology Scanner (Leica Biosystems).

### Digital Histology Analysis

Whole-slide images (WSI) from H&E and IHC staining were analyzed using QuPath 0.5.1 (P. Bankhead, 2017). Tumor and non-tumor regions were manually annotated prior to analysis. For H&E WSIs, the “cell detection” function was used to quantify overall cellularity (i.e., cell density). For IHC WSIs, images were thresholded before tumor and non-tumor annotation and classification. Positive staining and density maps were generated using the “positive cell detection” function based on nuclear or cytoplasmic DAB optical density (OD) depending on the stain pattern. Tumor regions were manually annotated using Aperio ImageScope digital histology viewer (v12.3.3, Leica Biosystems). Tumor annotation files (.xml) were converted to binary images, and tumor area was computed as the sum of the foreground pixels and converted to µm^2^ using the whole-slide image micron per pixel (MPP) ratio. Brain tissue was segmented in MATLAB using the *imgaussfilt* function with a Gaussian filter (σ = 3), followed by global mean-based thresholding. Total tissue area was calculated as the sum of foreground pixels in the binary image. Non-tumor tissue area was determined by subtracting tumor area from total brain tissue area. Both measures were converted to µm^2^ using the WSI MPP ratio. Nuclei in H&E images were identified based on Hematoxylin staining. Survivin-positive nuclei in IHC images were defined by QuPath classification as “positive 1+, 2+, or 3+.” Tumor and non-tumor nuclear density and survivin-positive cell density were computed by dividing the total nuclear or survivin-positive area by the corresponding tumor or non-tumor tissue area.

### Statistical analysis

Statistical significance was determined using a paired, two-tailed Student’s t-test. Significance thresholds were defined as *\*p<0.05*, *\*\*p<0.01*, *\*\*\*p<0.001*, and *\*\*\*\*p<0.0001*. Data are presented as mean + SEM from the indicated number of independent experiments.

## DATA AVAILABILITY

The data that support the findings of this study are available from the corresponding author upon reasonable request.

## ACKNOWLEDGMENTS

This work was supported by the Dr. Louis Sklarow Award and NIH/NHLBI grant R01HL163168 to Y.B.

## CONTRIBUTIONS

Y.B. designed the whole project and supervised all experiments; G.I., S.B., J.J., and Y.B. wrote the manuscript; G.I., S.B., J.J., R.D., B.A.S., B.T., K.P., S.B., J.E.T., T.A.I., R.Z., J.L., S.K., A.H.S., B.C.D., V.M.T., and Y.B. reviewed the data and the manuscript; G.I., R.D., B.T., K.P., and S.B. performed the bioinformatic analysis; G.I. and J.J. performed biological and biochemical assays and analyzed the data; S.B., B.A.S., J.L., and V.M.T. performed animal experiments, histological assays and analyzed the data. All authors have read and approved the article.

## AUTHOR DECLARATIONS

### Ethical Approval

Ethics approval is not required.

