## Supplemental figures for "Mechanotherapeutic Potential of Survivin in Glioblastoma"

### Supplementary Figure 1

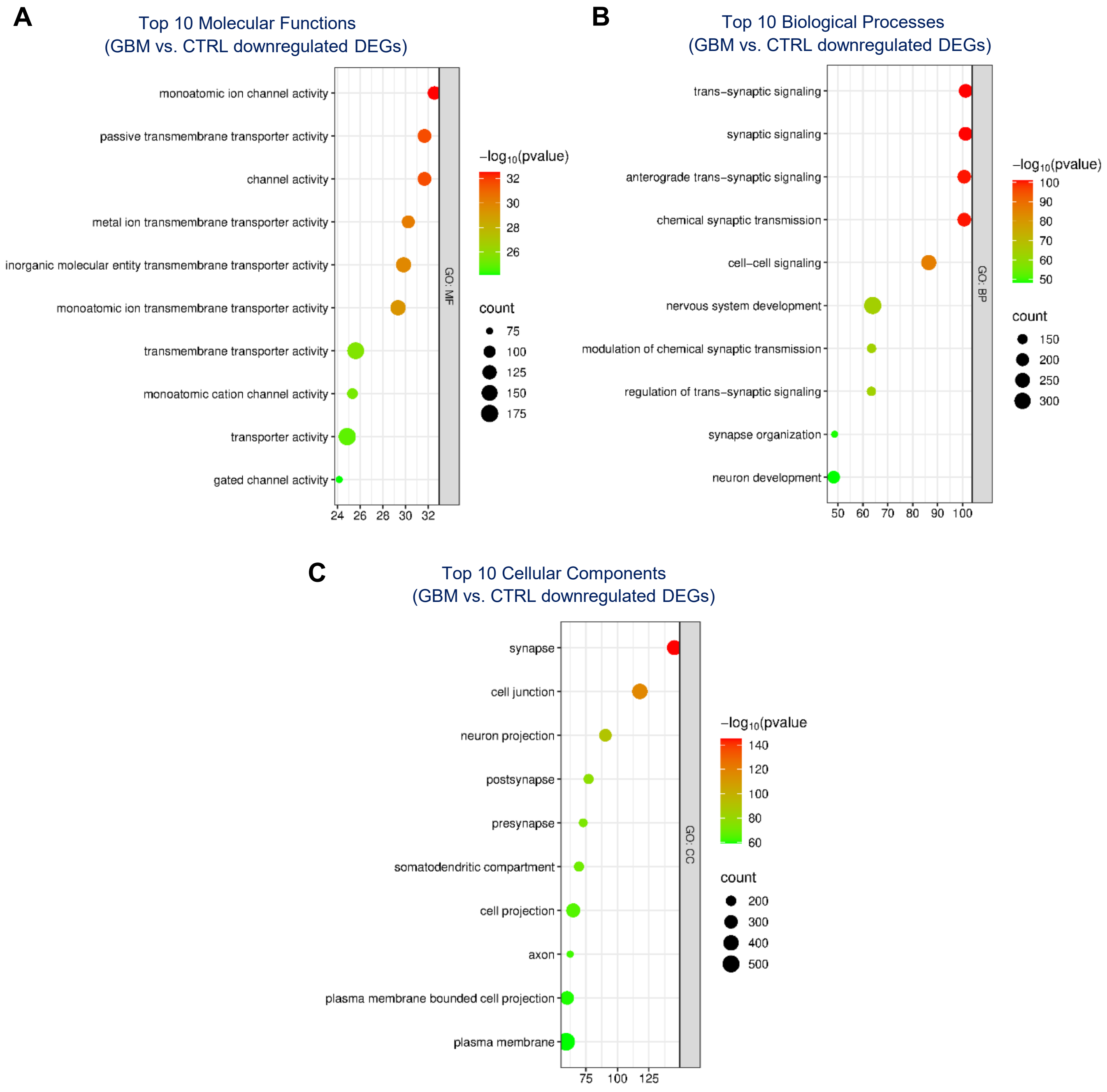


**Supplementary Figure 1. Downregulated DEGs are associated with transmembrane transport and synaptic activity.** Bubble plots depict the top 10 molecular functions (**A**), biological processes (**B**), and cellular components (**C**) for downregulated DEGs.

### Supplementary Figure 2

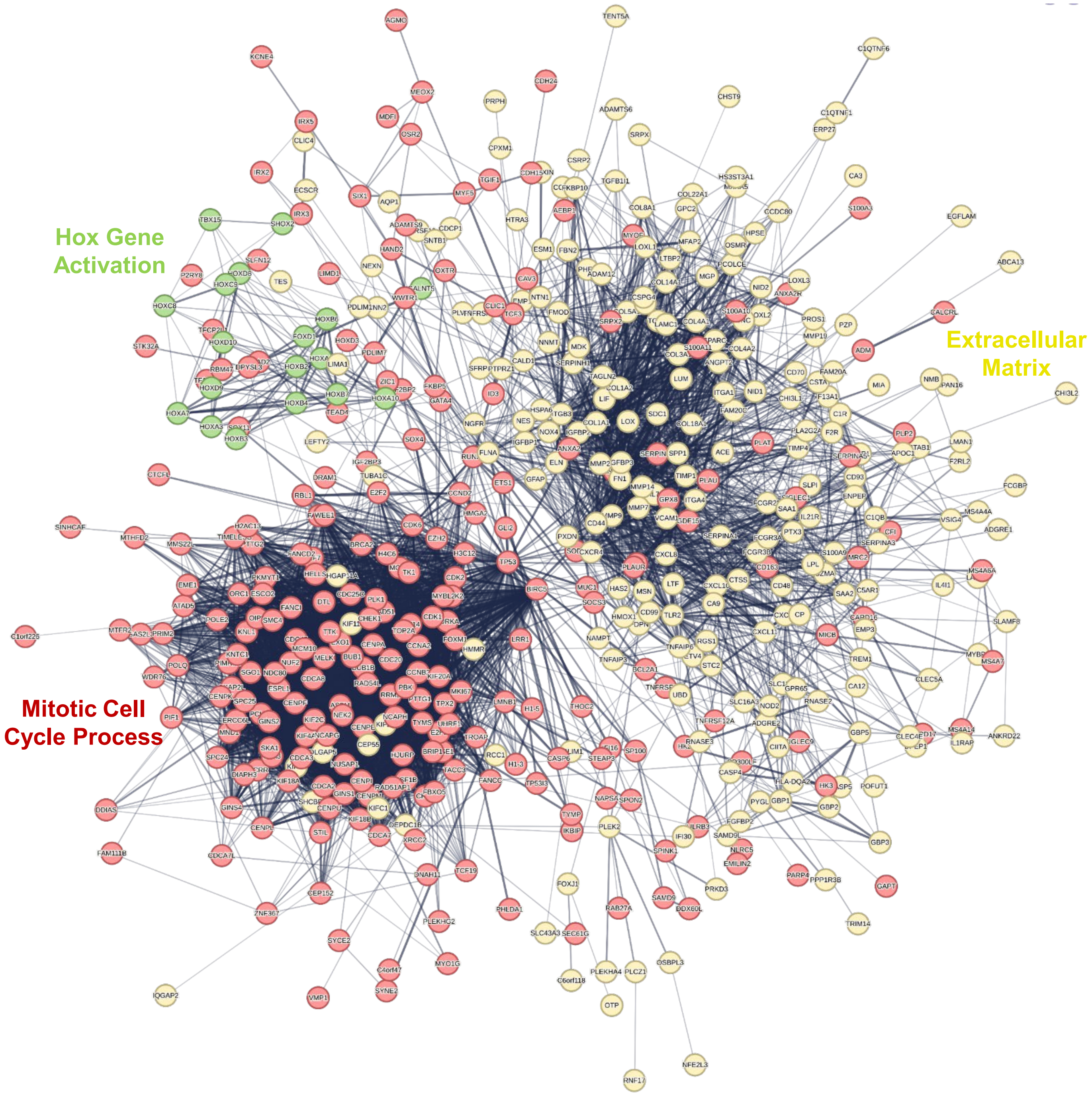


**Supplementary Figure 2. Protein-protein interaction network of upregulated DEGs reveals functional clusters related to cell cycle progression and ECM.** Protein-protein interaction network generated using the STRING database, showing functionally connected clusters among upregulated DEGs. K-means clustering grouped DEGs into three clusters, each named according to its most enriched molecular function, biological process, or cellular component.

### Supplementary Figure 3

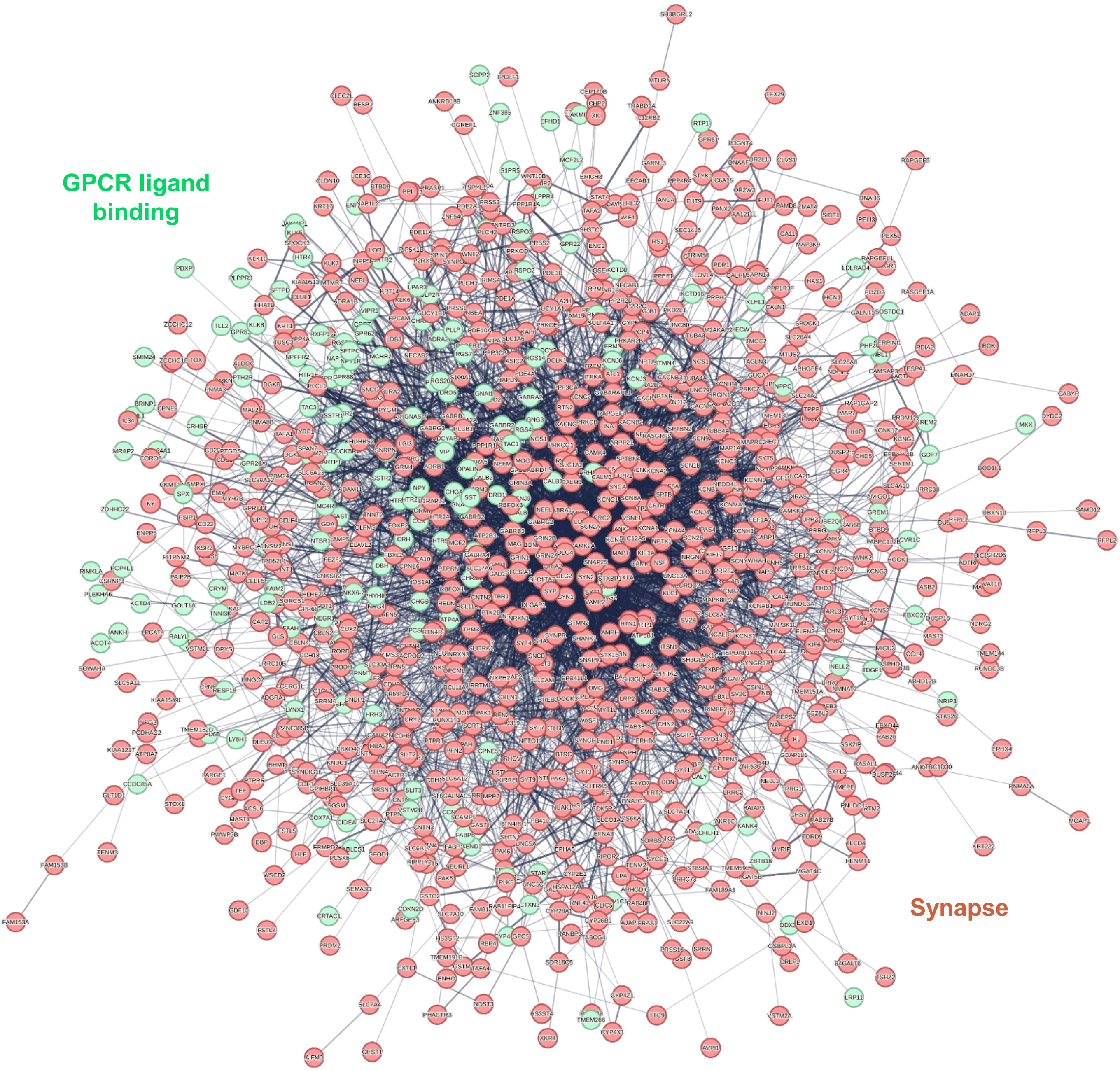


**Supplementary Figure 3. Protein-protein interaction network of downregulated DEGs reveals functional clusters related to GPCR signaling and synaptic processes.** Protein-protein interaction network generated using the STRING database, showing functionally connected clusters among downregulated DEGs. K-means clustering grouped these DEGs into two clusters, each named for its most enriched molecular function, biological process, or cellular component.

### Supplementary Figure 4

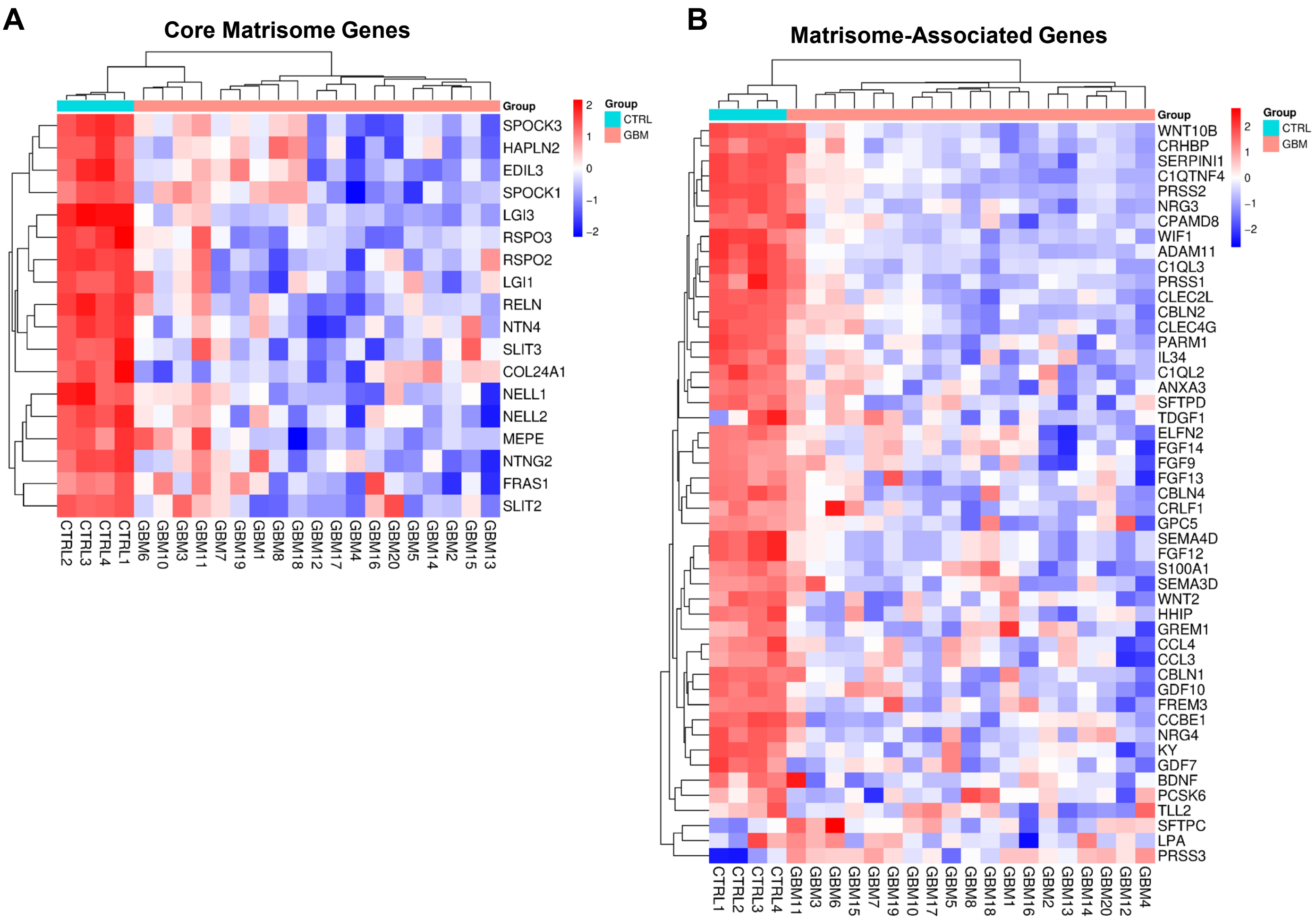


**Supplementary Figure 4. Matrisome analysis of downregulated DEGs.** Downregulated DEGs were uploaded to the Matrisome database to obtain gene annotations. Annotated genes were classified into core matrisome (**A**) and matrisome-associated (**B**) categories, and individual sample expression of each gene was visualized using SRplot in heatmaps.

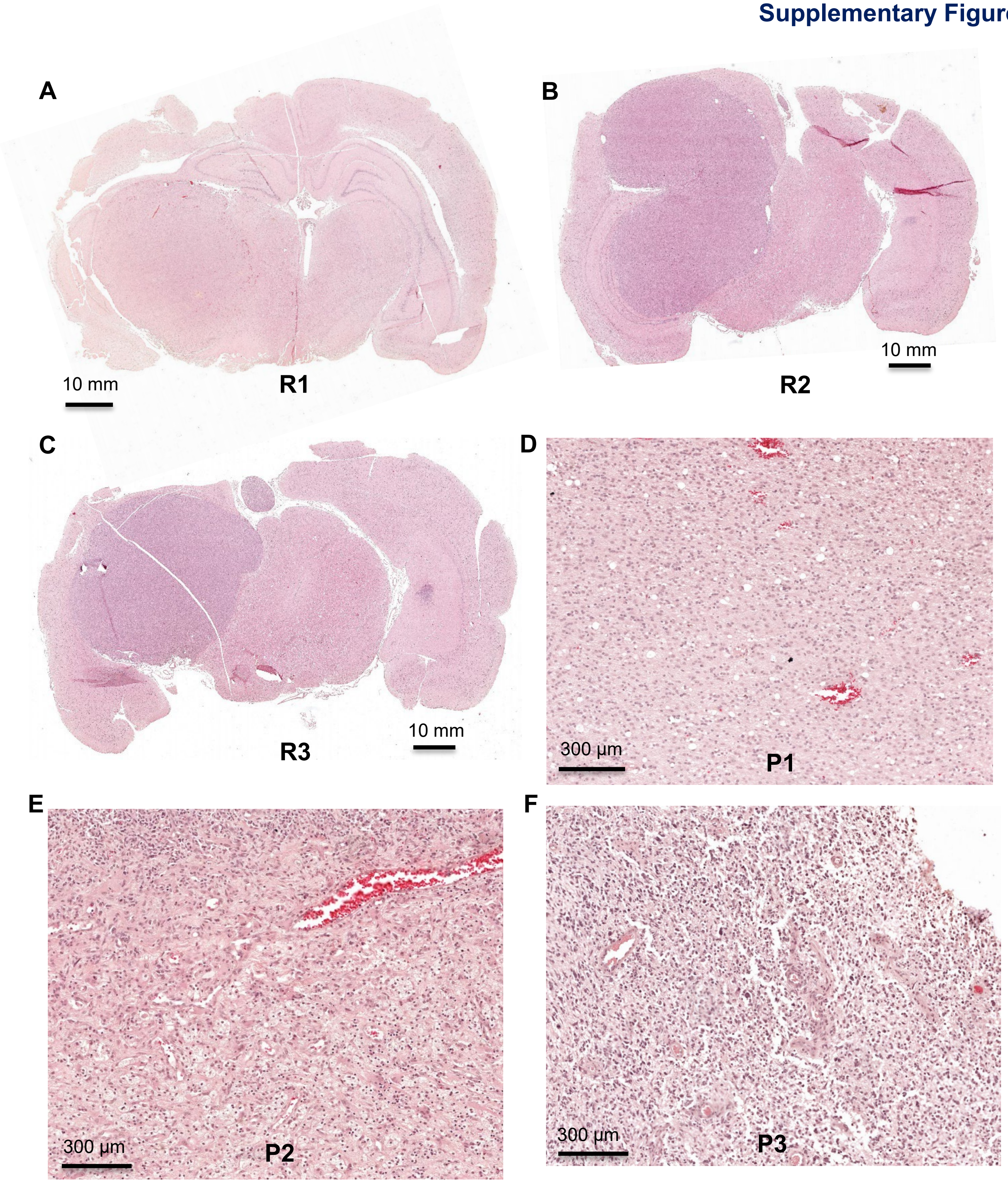


**Supplementary Figure 5. H&E staining of U87-implanted and human GBM tumor sections.** Representative hematoxylin and eosin (H&E) images are shown for U87-implanted tumors (**A–C**) and human GBM samples (**D–F**). U87-implanted tumor images are presented as whole-slide scans, while human GBM images are shown as selected regions of interest (ROIs). Whole-slide scanning acquired at 40x

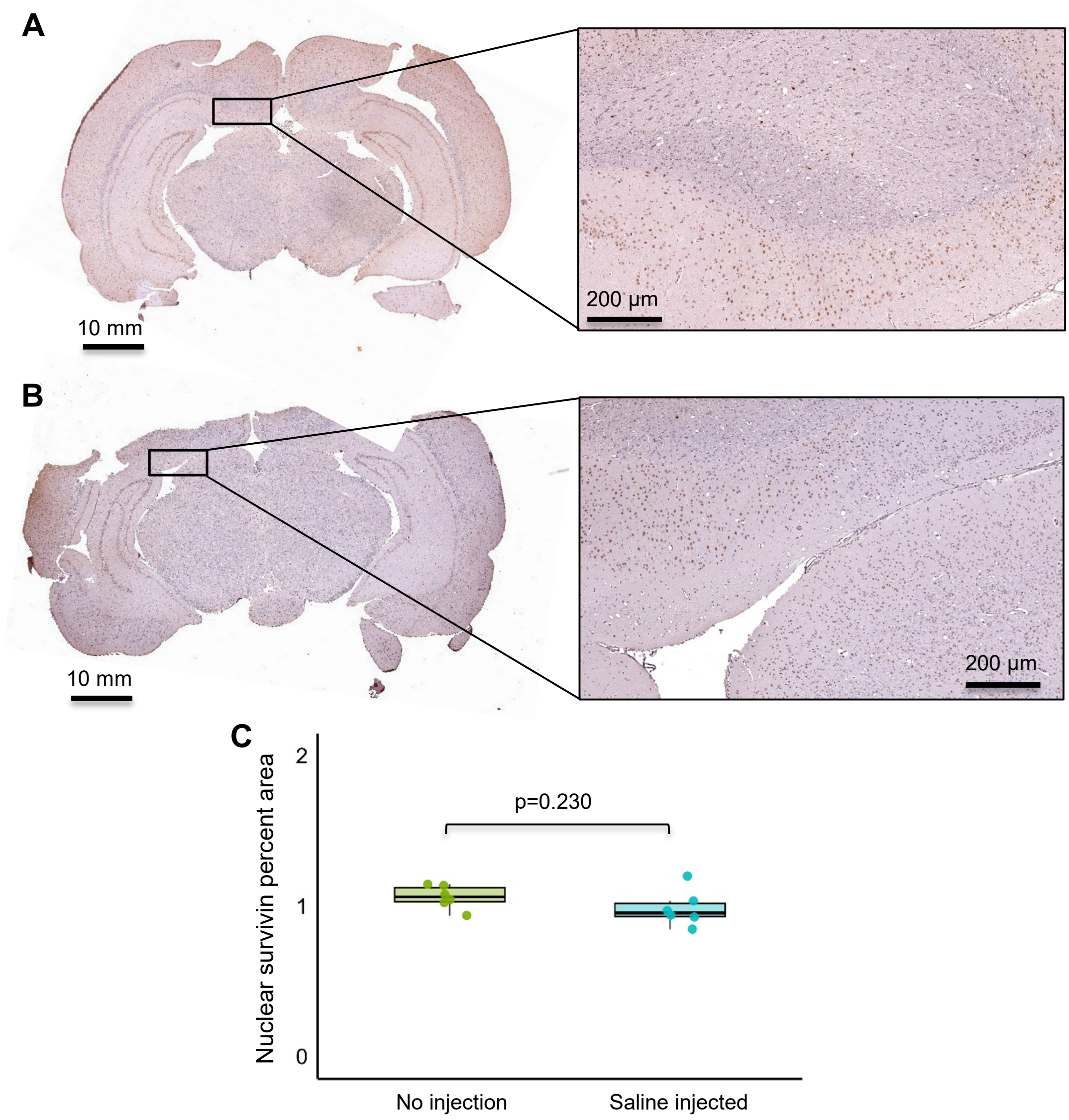


**Supplementary Figure 6. IHC survivin staining of control and saline-injected nude rats.** Control T-cell deficient nude rats (*n=6*) and saline-injected nude rats (*n=6*) were euthanized alongside U87-implanted rats. Following formalin-fixation, brain sections were subjected to IHC staining for survivin, and representative images are shown. Tissue sections and regions of interest are presented for control rats **(A)** and saline-injected rats **(B).** Whole-slide images were acquired at 40x magnification. Survivin expression was quantified as a percentage of survivin-positive nuclei relative to total nuclei in the brain section **(C).**

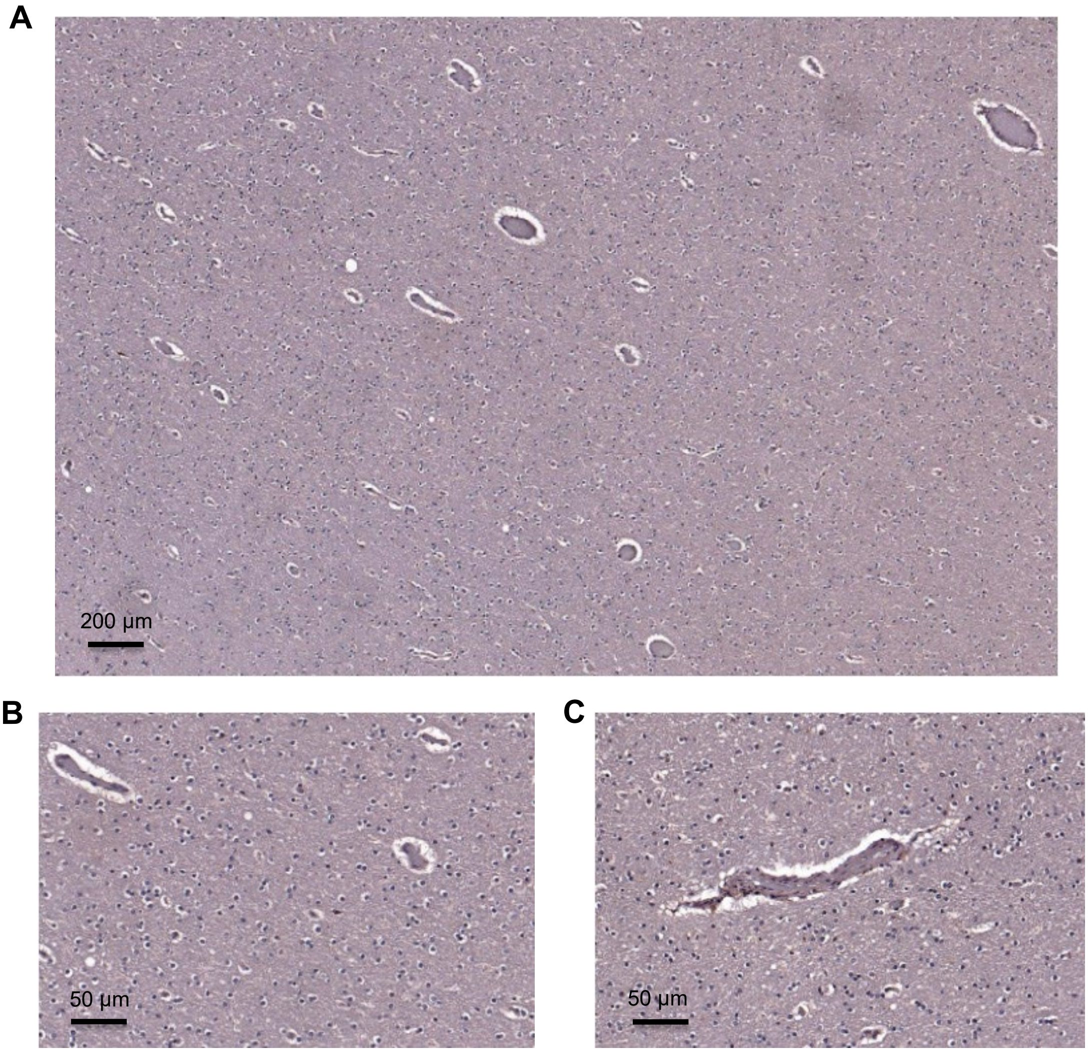


**Supplementary Figure 7. IHC survivin staining in non-GBM human brain.** Control human brain autopsies from non-GBM patients (*n=3*) were obtained from the NIH NeuroBioBank. Samples were immunohistochemically stained for survivin, and representative images (20x) are shown. Most regions exhibited negative staining (**A, B**), while occasional vascular areas showed positive survivin staining (**C**). Whole-slide images were acquired at 20x.

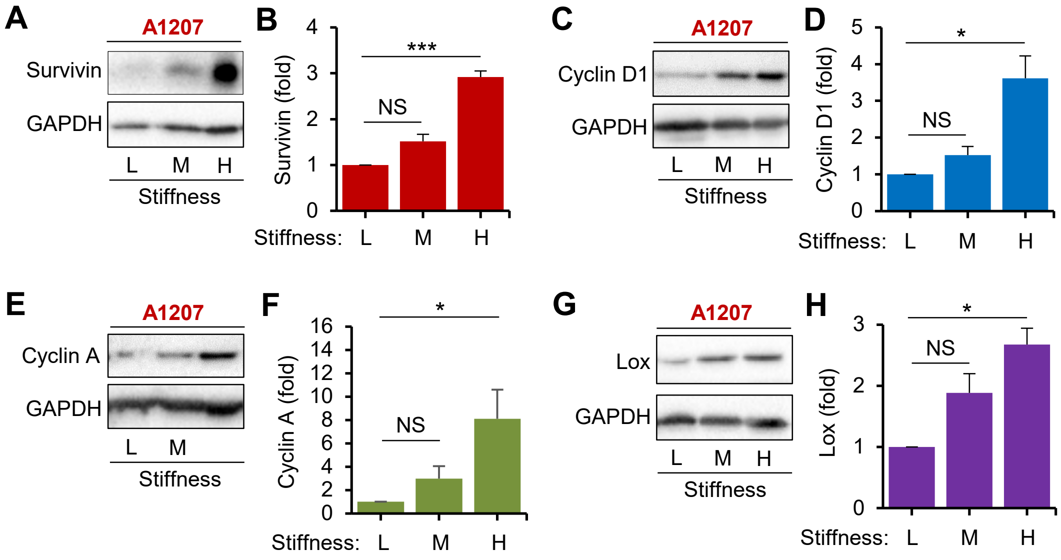


**Supplementary Figure 8. Stiffness-induced expression of survivin, cell cycle, and ECM proteins in A1207 cells.** Western blot analysis shows stiffness-induced changes in survivin (**A–B**), cyclin D1 (**C–D**), cyclin A (**E–F**), and lysyl oxidase (LOX) (**G–H**) expression in A1207 cells after 24 h of incubation on low (L), medium (M), and high (H) stiffness polyacrylamide hydrogels. Protein levels were normalized to GAPDH and expressed as fold change relative to low stiffness (control). Representative blot images are shown alongside quantification. Each analysis includes *n=3-6*; significance: **p < 0.05*, *p < 0.01*, *NS*: not significant.

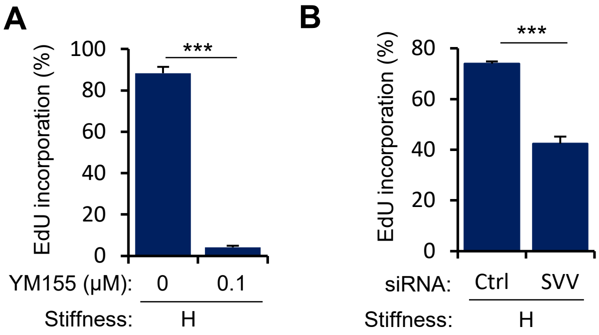


**Supplemental Figure 9. Survivin inhibition reduces S-phase entry in U87 Cells.** EdU incorporation was performed to assess newly synthesized DNA as a measure of proliferation. U87 cells cultured on high-stiffness hydrogels were treated with DMSO or 0.1 µM YM155 (**A**), or transfected with 200 nM control or BIRC5 siRNA (**B**), and incubated with EdU for 24 h prior to fixation. The EdU reaction was then performed, and slides were imaged with at least five fields of view (FOV) across four independent experiments. Each analysis includes $n = 4$; significance: $*p < 0.05$, $**p < 0.01$, $*p < 0.001$, *NS*: not significant.
