## Supplemental Table 1 for "Mechanotherapeutic Potential of Survivin in Glioblastoma"

**Supplementary Table 1. Transcriptomic analysis identifies genes significantly altered in GBM tumors**. RNA-sequencing analysis of publicly available GBM tumors and non-neoplastic brain samples was performed. Genes were filtered using q-value $\leq 0.05$ and $\log_2 FC \geq 0.75$. and ranked by $\log_2 FC$. Survivin (*BIRC5*) was identified as the third most upregulated gene in GBM tumors.

| Gene Name | adj. p Value | p Value | Fold Change | $\log_2 FC$ |
|---|---|---|---|---|
| NCAPG | 7.76E-11 | 5.43E-15 | 5.44742791 | 2.445575 |
| CCNB2 | 2.09E-10 | 2.54E-14 | 5.3586363 | 2.421866 |
| **BIRC5** | 1.98E-10 | 1.93E-14 | 4.84587066 | 2.276756 |
| PBK | 2.46E-09 | 1.08E-12 | 4.73741744 | 2.244101 |
| E2F2 | 2.28E-09 | 8.20E-13 | 4.36517563 | 2.12604 |
| FAM64A | 2.87E-09 | 1.36E-12 | 4.28267589 | 2.098513 |
| NDC80 | 7.31E-11 | 1.78E-15 | 4.1508477 | 2.053406 |
| TOP2A | 3.10E-08 | 7.42E-11 | 4.12254345 | 2.043535 |
| CENPA | 4.90E-09 | 3.23E-12 | 4.05972808 | 2.021383 |
| DLGAP5 | 1.13E-07 | 6.40E-10 | 3.97993178 | 1.992744 |
| SPOCD1 | 1.18E-05 | 5.24E-07 | 3.94427899 | 1.979762 |
| RRM2 | 1.17E-09 | 2.00E-13 | 3.92427026 | 1.972424 |
| MELK | 2.87E-09 | 1.60E-12 | 3.87367119 | 1.953702 |
| GTSE1 | 4.44E-09 | 2.82E-12 | 3.80325323 | 1.927234 |
| COL3A1 | 3.40E-06 | 9.69E-08 | 3.80011479 | 1.926043 |
| SERPINA3 | 1.42E-07 | 9.02E-10 | 3.74897083 | 1.906495 |
| KIF20A | 1.07E-08 | 1.21E-11 | 3.73090364 | 1.899525 |
| H19 | 0.00415 | 0.000844 | 3.72072148 | 1.895582 |
| ABCC3 | 3.80E-07 | 4.10E-09 | 3.66571164 | 1.874093 |
| UHRF1 | 1.26E-08 | 1.56E-11 | 3.61890516 | 1.855553 |
| ASPM | 1.34E-08 | 1.76E-11 | 3.60258433 | 1.849032 |
| CDCA2 | 9.16E-09 | 9.61E-12 | 3.57596273 | 1.838332 |
| HOXA7 | 0.00495 | 0.00105 | 3.57547967 | 1.838137 |
| GSC | 1.62E-06 | 3.44E-08 | 3.55297318 | 1.829027 |
| NUSAP1 | 6.30E-09 | 5.38E-12 | 3.54613584 | 1.826248 |
| C21orf62 | 6.79E-08 | 2.80E-10 | 3.49235254 | 1.804199 |
| UBD | 4.21E-05 | 2.71E-06 | 3.45601265 | 1.789109 |
| COL1A1 | 9.11E-05 | 7.35E-06 | 3.4449678 | 1.784491 |
| ANXA2 | 3.25E-06 | 9.11E-08 | 3.44183658 | 1.783179 |
| CRNDE | 3.43E-07 | 3.48E-09 | 3.43771084 | 1.781448 |
| TNC | 1.95E-07 | 1.44E-09 | 3.41554405 | 1.772115 |
| HOXD10 | 0.000432 | 5.25E-05 | 3.40874623 | 1.769241 |
| BUB1 | 5.72E-09 | 4.47E-12 | 3.30960077 | 1.726657 |
| KIFC1 | 3.34E-08 | 9.37E-11 | 3.30058969 | 1.722724 |
| SLC47A2 | 2.89E-05 | 1.67E-06 | 3.29588385 | 1.720665 |
| TNFRSF12A | 3.21E-07 | 3.12E-09 | 3.24491908 | 1.698183 |

| | | | | |
|---|---|---|---|---|
| SAA1 | 0.00513 | 0.0011 | 3.23814839 | 1.695169 |
| HJURP | 2.05E-08 | 3.40E-11 | 3.22463529 | 1.689136 |
| KIF18B | 1.33E-07 | 8.18E-10 | 3.21904066 | 1.686631 |
| CD44 | 6.53E-07 | 9.09E-09 | 3.21545456 | 1.685023 |
| CENPK | 3.35E-09 | 1.96E-12 | 3.19537068 | 1.675983 |
| CEP55 | 8.34E-09 | 7.55E-12 | 3.18354024 | 1.670632 |
| DTL | 1.14E-08 | 1.34E-11 | 3.16729336 | 1.663251 |
| F2R | 5.46E-08 | 1.97E-10 | 3.16427591 | 1.661875 |
| TICRR | 2.70E-08 | 5.79E-11 | 3.16181838 | 1.660755 |
| FAM20C | 7.96E-06 | 3.13E-07 | 3.14579945 | 1.653427 |
| CDC25C | 3.34E-08 | 9.06E-11 | 3.13621247 | 1.649023 |
| ANXA2P3 | 1.65E-06 | 3.53E-08 | 3.12238557 | 1.642649 |
| KIAA0101 | 5.24E-09 | 3.83E-12 | 3.09993636 | 1.632239 |
| TTK | 6.62E-08 | 2.65E-10 | 3.09923833 | 1.631914 |
| VMP1 | 5.56E-07 | 7.13E-09 | 3.07869233 | 1.622318 |
| HOTAIRM1 | 6.46E-06 | 2.37E-07 | 3.07565631 | 1.620894 |
| ESCO2 | 2.58E-08 | 5.09E-11 | 3.0569164 | 1.612077 |
| HOXB2 | 0.00167 | 0.000277 | 3.05376046 | 1.610587 |
| CENPU | 2.04E-08 | 3.34E-11 | 3.04081511 | 1.604458 |
| MKI67 | 5.40E-07 | 6.84E-09 | 3.03696188 | 1.602629 |
| KNL1 | 1.42E-07 | 8.95E-10 | 3.00439749 | 1.587076 |
| LTF | 0.0187 | 0.00533 | 3.00413303 | 1.586949 |
| ANKRD22 | 1.38E-07 | 8.62E-10 | 2.96561784 | 1.568333 |
| KIF4A | 1.72E-09 | 3.78E-13 | 2.92692788 | 1.549387 |
| E2F7 | 1.36E-08 | 1.82E-11 | 2.9263755 | 1.549115 |
| SLAMF8 | 2.12E-07 | 1.68E-09 | 2.91799316 | 1.544977 |
| SERPINA5 | 4.11E-06 | 1.26E-07 | 2.91028149 | 1.541159 |
| CDC45 | 3.34E-08 | 9.30E-11 | 2.89984744 | 1.535977 |
| CKAP2L | 3.41E-08 | 9.64E-11 | 2.89898667 | 1.535549 |
| IRX5 | 2.43E-05 | 1.33E-06 | 2.89209953 | 1.532117 |
| TMSB15A | 0.000324 | 3.66E-05 | 2.84858746 | 1.510247 |
| CHI3L1 | 2.96E-05 | 1.73E-06 | 2.83249282 | 1.502072 |
| CHI3L2 | 5.28E-05 | 3.64E-06 | 2.82237053 | 1.496907 |
| TRMU///GTSE1 | 1.15E-07 | 6.61E-10 | 2.8207582 | 1.496083 |
| LIF | 0.000883 | 0.000126 | 2.81997818 | 1.495684 |
| CD163 | 3.98E-06 | 1.20E-07 | 2.81983139 | 1.495609 |
| MMP19 | 0.00035 | 4.03E-05 | 2.78680257 | 1.478611 |
| CENPF | 1.59E-07 | 1.10E-09 | 2.78124936 | 1.475733 |
| SRPX2 | 6.76E-06 | 2.51E-07 | 2.77954589 | 1.474849 |
| FCGBP | 2.72E-07 | 2.34E-09 | 2.77760009 | 1.473839 |
| HOXD3 | 2.68E-05 | 1.52E-06 | 2.73915081 | 1.453729 |
| CDK1 | 5.99E-08 | 2.31E-10 | 2.73842107 | 1.453344 |
| CA9 | 0.000412 | 4.95E-05 | 2.73428647 | 1.451164 |

| | | | | |
|---|---|---|---|---|
| GPX8 | 8.16E-07 | 1.30E-08 | 2.73141419 | 1.449648 |
| IGFBP2 | 3.73E-08 | 1.09E-10 | 2.69731049 | 1.431522 |
| HOXB3 | 0.0227 | 0.00677 | 2.6964577 | 1.431065 |
| MMP7 | 0.0201 | 0.00582 | 2.69588097 | 1.430757 |
| KIF18A | 1.49E-08 | 2.22E-11 | 2.69562218 | 1.430618 |
| SPC25 | 3.94E-08 | 1.24E-10 | 2.69402045 | 1.429761 |
| CCNA2 | 6.96E-08 | 2.88E-10 | 2.69345059 | 1.429456 |
| MXRA5 | 5.31E-06 | 1.80E-07 | 2.6899581 | 1.427584 |
| MEX3A | 7.98E-07 | 1.26E-08 | 2.67230615 | 1.418085 |
| SGO1 | 9.48E-08 | 4.89E-10 | 2.66299607 | 1.41305 |
| HOXC9 | 0.00243 | 0.000437 | 2.65670474 | 1.409638 |
| OIP5 | 6.98E-08 | 2.94E-10 | 2.65448189 | 1.40843 |
| TROAP | 9.94E-08 | 5.36E-10 | 2.64857893 | 1.405219 |
| VEGFA | 1.06E-05 | 4.57E-07 | 2.64577211 | 1.403689 |
| LOXL2 | 7.50E-06 | 2.91E-07 | 2.63291692 | 1.396662 |
| EZH2 | 3.50E-07 | 3.62E-09 | 2.62937826 | 1.394722 |
| MMP9 | 0.00112 | 0.000169 | 2.62329418 | 1.39138 |
| IQGAP2 | 1.15E-07 | 6.62E-10 | 2.61984762 | 1.389483 |
| RAB42 | 1.38E-08 | 1.89E-11 | 2.60262256 | 1.379966 |
| CFI | 2.28E-06 | 5.56E-08 | 2.58834657 | 1.372031 |
| TGFBI | 6.37E-06 | 2.32E-07 | 2.57812515 | 1.366322 |
| IGF2BP3 | 0.000105 | 8.75E-06 | 2.55199199 | 1.351624 |
| CASC23 | 0.00834 | 0.00197 | 2.54258658 | 1.346297 |
| SKA3 | 8.34E-09 | 7.73E-12 | 2.54193335 | 1.345926 |
| KIF2C | 6.98E-08 | 2.92E-10 | 2.53933776 | 1.344452 |
| TUBA1C | 1.89E-06 | 4.32E-08 | 2.53284378 | 1.340758 |
| ANGPT2 | 1.33E-07 | 8.16E-10 | 2.52939509 | 1.338792 |
| ANXA2P1 | 1.12E-05 | 4.88E-07 | 2.52393928 | 1.335677 |
| E2F8 | 2.73E-06 | 7.07E-08 | 2.50932702 | 1.327301 |
| DDIAS | 4.25E-08 | 1.37E-10 | 2.50843769 | 1.326789 |
| TREM1 | 0.000396 | 4.70E-05 | 2.50697029 | 1.325945 |
| HOXA3 | 0.00747 | 0.00173 | 2.50307132 | 1.323699 |
| ESM1 | 0.00048 | 5.97E-05 | 2.50192319 | 1.323038 |
| RNF17 | 0.00501 | 0.00106 | 2.49596517 | 1.319598 |
| GFAP | 2.90E-06 | 7.74E-08 | 2.49226141 | 1.317455 |
| SERPINE1 | 0.000663 | 8.86E-05 | 2.48460033 | 1.313014 |
| CXCL10 | 0.00308 | 0.000585 | 2.48278305 | 1.311958 |
| PLEK2 | 2.22E-06 | 5.29E-08 | 2.47361693 | 1.306622 |
| SOX11 | 4.90E-07 | 5.91E-09 | 2.4681858 | 1.303451 |
| ARHGAP11A | 5.26E-07 | 6.60E-09 | 2.46766405 | 1.303146 |
| NES | 2.61E-08 | 5.21E-11 | 2.46737483 | 1.302977 |
| CD93 | 2.72E-07 | 2.35E-09 | 2.46516976 | 1.301687 |
| ELN | 1.10E-05 | 4.81E-07 | 2.46440352 | 1.301239 |

| | | | | |
|---|---|---|---|---|
| TIMP1 | 6.55E-05 | 4.79E-06 | 2.46388787 | 1.300937 |
| NOX4 | 3.29E-07 | 3.26E-09 | 2.46043073 | 1.298911 |
| CPXM1 | 2.88E-07 | 2.63E-09 | 2.45802655 | 1.297501 |
| MS4A6A | 2.80E-07 | 2.51E-09 | 2.43444111 | 1.283591 |
| ESPL1 | 7.37E-08 | 3.40E-10 | 2.42796816 | 1.27975 |
| PLCZ1 | 0.0148 | 0.004 | 2.42574366 | 1.278427 |
| PDPN | 9.56E-05 | 7.80E-06 | 2.42486445 | 1.277904 |
| CHEK1 | 1.07E-08 | 1.21E-11 | 2.42260299 | 1.276558 |
| METTL7B | 0.000142 | 1.28E-05 | 2.41772591 | 1.273651 |
| NUF2 | 2.97E-07 | 2.78E-09 | 2.41485238 | 1.271935 |
| GDF15 | 0.00021 | 2.13E-05 | 2.41420988 | 1.271551 |
| FOXM1 | 6.68E-08 | 2.72E-10 | 2.4078972 | 1.267774 |
| CDCA7 | 1.30E-08 | 1.65E-11 | 2.40376557 | 1.265296 |
| ADAM12 | 1.42E-05 | 6.68E-07 | 2.40211896 | 1.264308 |
| PLEKHA4 | 3.34E-08 | 9.25E-11 | 2.39934248 | 1.262639 |
| HMMR | 3.76E-07 | 4.03E-09 | 2.39898261 | 1.262423 |
| CENPM | 1.21E-07 | 7.08E-10 | 2.39219766 | 1.258337 |
| NCAPH | 1.24E-07 | 7.53E-10 | 2.39074258 | 1.257459 |
| CSTA | 2.23E-05 | 1.19E-06 | 2.38817027 | 1.255906 |
| KIF14 | 1.08E-05 | 4.71E-07 | 2.38119472 | 1.251686 |
| PTX3 | 0.00099 | 0.000145 | 2.37911218 | 1.250423 |
| SFRP4 | 5.55E-05 | 3.90E-06 | 2.37906156 | 1.250393 |
| ENPEP | 2.03E-07 | 1.56E-09 | 2.36811526 | 1.243739 |
| CA12 | 0.000753 | 0.000104 | 2.36758513 | 1.243416 |
| SKA1 | 1.21E-07 | 7.23E-10 | 2.36209322 | 1.240066 |
| EXO1 | 2.97E-07 | 2.76E-09 | 2.35335946 | 1.234722 |
| COL1A2 | 8.67E-05 | 6.90E-06 | 2.33546464 | 1.22371 |
| NEK2 | 4.17E-06 | 1.29E-07 | 2.32625701 | 1.218011 |
| SIX1 | 4.02E-05 | 2.55E-06 | 2.32537921 | 1.217466 |
| OSR2 | 0.000428 | 5.20E-05 | 2.32430243 | 1.216798 |
| CELSR1 | 0.000224 | 2.30E-05 | 2.31651384 | 1.211955 |
| WEE1 | 5.13E-09 | 3.51E-12 | 2.31298098 | 1.209753 |
| MMP14 | 1.18E-06 | 2.16E-08 | 2.30542095 | 1.20503 |
| COL8A1 | 0.00029 | 3.18E-05 | 2.30105968 | 1.202298 |
| HELLS | 9.81E-08 | 5.27E-10 | 2.29709178 | 1.199809 |
| ERCC6L | 2.27E-06 | 5.51E-08 | 2.29683067 | 1.199645 |
| LMNB1 | 4.13E-07 | 4.57E-09 | 2.29204902 | 1.196638 |
| LAMP3 | 5.52E-06 | 1.90E-07 | 2.29005953 | 1.195385 |
| ERP27 | 0.000103 | 8.60E-06 | 2.28722913 | 1.193601 |
| SERPINH1 | 1.58E-07 | 1.09E-09 | 2.28122145 | 1.189807 |
| STEAP3 | 1.88E-06 | 4.27E-08 | 2.27758816 | 1.187507 |
| RAD51 | 3.72E-08 | 1.07E-10 | 2.27666781 | 1.186924 |
| COL4A1 | 2.64E-06 | 6.78E-08 | 2.26678726 | 1.180649 |

| | | | | |
|---|---|---|---|---|
| SHOX2 | 7.51E-05 | 5.71E-06 | 2.2624067 | 1.177858 |
| CA3 | 0.00223 | 0.000394 | 2.25471971 | 1.172948 |
| GATA4 | 0.00816 | 0.00192 | 2.25335216 | 1.172073 |
| SRPX | 5.64E-06 | 1.96E-07 | 2.25190739 | 1.171148 |
| PLAU | 0.000483 | 6.03E-05 | 2.25055481 | 1.170281 |
| CCDC80 | 1.60E-07 | 1.12E-09 | 2.24643466 | 1.167637 |
| TACC3 | 4.92E-06 | 1.63E-07 | 2.23860952 | 1.162603 |
| BUB1B | 1.62E-06 | 3.44E-08 | 2.22099462 | 1.151206 |
| LOC154761 | 9.93E-05 | 8.18E-06 | 2.21999173 | 1.150554 |
| LUM | 0.00024 | 2.50E-05 | 2.21722811 | 1.148757 |
| WWTR1 | 4.04E-05 | 2.57E-06 | 2.20846658 | 1.143045 |
| CD70 | 0.0491 | 0.0176 | 2.20386339 | 1.140035 |
| HIST1H3B | 9.48E-08 | 4.90E-10 | 2.19917727 | 1.136964 |
| MCM10 | 2.78E-06 | 7.21E-08 | 2.19733357 | 1.135754 |
| STC2 | 0.00187 | 0.000319 | 2.19105606 | 1.131626 |
| RAD54L | 1.34E-06 | 2.58E-08 | 2.18632125 | 1.128505 |
| MEOX2 | 0.00102 | 0.00015 | 2.18558047 | 1.128017 |
| POLQ | 9.54E-07 | 1.62E-08 | 2.18043079 | 1.124613 |
| MYF5 | 0.0409 | 0.0141 | 2.18008971 | 1.124388 |
| GPR65 | 6.55E-07 | 9.13E-09 | 2.17879053 | 1.123528 |
| PTPN7 | 1.73E-06 | 3.78E-08 | 2.16910877 | 1.117102 |
| COL4A2 | 1.45E-07 | 9.23E-10 | 2.15572806 | 1.108175 |
| ADM | 0.000292 | 3.21E-05 | 2.1542551 | 1.107189 |
| NAPSB | 0.000191 | 1.88E-05 | 2.15309713 | 1.106413 |
| TBX15 | 4.73E-06 | 1.53E-07 | 2.15193545 | 1.105635 |
| IFI30 | 2.08E-06 | 4.86E-08 | 2.1485261 | 1.103347 |
| LEFTY2 | 0.000916 | 0.000132 | 2.14173116 | 1.098777 |
| HOXA10 | 0.0024 | 0.000431 | 2.13869418 | 1.09673 |
| IBSP | 0.00456 | 0.000946 | 2.13687897 | 1.095505 |
| LOX | 0.00288 | 0.000538 | 2.13598187 | 1.094899 |
| IRX2 | 0.03 | 0.00953 | 2.12872782 | 1.089992 |
| GBP2 | 4.93E-06 | 1.64E-07 | 2.12082121 | 1.084623 |
| GAS2L3 | 1.49E-06 | 3.04E-08 | 2.12007485 | 1.084115 |
| NNMT | 0.00123 | 0.000189 | 2.11616585 | 1.081453 |
| CP | 0.000387 | 4.56E-05 | 2.11523976 | 1.080821 |
| TFCP2L1 | 0.000506 | 6.38E-05 | 2.11240228 | 1.078885 |
| ADGRE1 | 2.60E-05 | 1.45E-06 | 2.11214679 | 1.07871 |
| COL22A1 | 0.000106 | 8.89E-06 | 2.11086747 | 1.077836 |
| CXCL8 | 0.0204 | 0.00592 | 2.10897369 | 1.076541 |
| ORC1 | 5.59E-07 | 7.20E-09 | 2.10774655 | 1.075701 |
| SAA2 | 0.0184 | 0.00523 | 2.10654218 | 1.074877 |
| IKBIP | 5.33E-08 | 1.87E-10 | 2.10335549 | 1.072693 |
| CLEC4E | 0.0015 | 0.000242 | 2.0982192 | 1.069165 |

| | | | | |
|---|---|---|---|---|
| GBP1 | 4.15E-05 | 2.66E-06 | 2.09656739 | 1.068029 |
| ASF1B | 3.56E-07 | 3.69E-09 | 2.09252056 | 1.065242 |
| CDCA8 | 1.44E-06 | 2.91E-08 | 2.0917066 | 1.064681 |
| LINC01206 | 0.0146 | 0.00393 | 2.09159569 | 1.064604 |
| OSMR | 7.26E-05 | 5.49E-06 | 2.09076629 | 1.064032 |
| IL1RAP | 2.57E-05 | 1.43E-06 | 2.08854264 | 1.062497 |
| MTFR2 | 7.44E-08 | 3.45E-10 | 2.08557992 | 1.060449 |
| SNTB1 | 1.71E-08 | 2.62E-11 | 2.08468788 | 1.059831 |
| HOXA5 | 0.0183 | 0.00518 | 2.08440266 | 1.059634 |
| RDH10 | 1.79E-05 | 9.07E-07 | 2.08115122 | 1.057382 |
| KIF15 | 5.39E-07 | 6.80E-09 | 2.08059044 | 1.056993 |
| EMP1 | 1.57E-05 | 7.60E-07 | 2.08041623 | 1.056872 |
| RNASE2 | 0.000111 | 9.37E-06 | 2.07960164 | 1.056307 |
| HTRA3 | 2.32E-05 | 1.25E-06 | 2.07514565 | 1.053213 |
| S100A9 | 0.00164 | 0.000271 | 2.07273359 | 1.051535 |
| CXCL9 | 0.00962 | 0.00235 | 2.07233337 | 1.051256 |
| SPC24 | 4.77E-06 | 1.56E-07 | 2.07137218 | 1.050587 |
| ADGRE2 | 1.02E-05 | 4.34E-07 | 2.07080887 | 1.050194 |
| SHCBP1 | 3.81E-08 | 1.14E-10 | 2.06900782 | 1.048939 |
| MMP2 | 4.44E-06 | 1.41E-07 | 2.06839224 | 1.04851 |
| TSPAN16 | 0.031 | 0.00992 | 2.06462219 | 1.045878 |
| ANXA1 | 2.74E-05 | 1.56E-06 | 2.06302914 | 1.044764 |
| PDLIM1 | 6.29E-06 | 2.27E-07 | 2.06188076 | 1.043961 |
| NOD2 | 8.78E-05 | 7.02E-06 | 2.06092114 | 1.043289 |
| KIF23 | 8.33E-07 | 1.35E-08 | 2.05939304 | 1.042219 |
| CRISPLD1 | 0.000327 | 3.70E-05 | 2.05796907 | 1.041221 |
| IGFBP3 | 0.00174 | 0.000292 | 2.05233096 | 1.037263 |
| HPSE | 0.00012 | 1.03E-05 | 2.05158155 | 1.036737 |
| ITGA1 | 1.70E-06 | 3.68E-08 | 2.05148414 | 1.036668 |
| CPS1-IT1 | 5.60E-05 | 3.95E-06 | 2.05076844 | 1.036165 |
| C15orf48 | 0.00114 | 0.000172 | 2.05038027 | 1.035892 |
| NTN1 | 5.44E-08 | 1.94E-10 | 2.04857413 | 1.03462 |
| FANCD2 | 6.30E-07 | 8.60E-09 | 2.04674178 | 1.033329 |
| ANXA2R | 0.00011 | 9.31E-06 | 2.0446969 | 1.031887 |
| TPX2 | 4.52E-07 | 5.26E-09 | 2.03869549 | 1.027646 |
| PLP2 | 4.25E-05 | 2.74E-06 | 2.03723202 | 1.02661 |
| MYBL2 | 1.21E-05 | 5.46E-07 | 2.03334807 | 1.023857 |
| CNGA3 | 0.00736 | 0.0017 | 2.03327295 | 1.023804 |
| TFPI | 2.43E-05 | 1.33E-06 | 2.0320238 | 1.022917 |
| CMTM3 | 2.28E-09 | 8.29E-13 | 2.02977329 | 1.021319 |
| FCGR2B | 0.00315 | 0.000601 | 2.02931384 | 1.020992 |
| HIST1H4K | 1.85E-07 | 1.36E-09 | 2.02898936 | 1.020761 |
| PXDN | 5.94E-08 | 2.28E-10 | 2.0279236 | 1.020003 |

| | | | | |
|---|---|---|---|---|
| PLK1 | 1.04E-06 | 1.80E-08 | 2.02761607 | 1.019785 |
| TCF19 | 7.36E-08 | 3.38E-10 | 2.02687132 | 1.019255 |
| HS3ST3A1 | 0.0451 | 0.0159 | 2.02539122 | 1.018201 |
| MYBPH | 0.00345 | 0.000672 | 2.02475633 | 1.017748 |
| HIST1H3D | 7.36E-08 | 3.33E-10 | 2.02035106 | 1.014606 |
| HOXD9 | 6.10E-05 | 4.40E-06 | 2.01889069 | 1.013563 |
| LILRB3 | 0.000428 | 5.20E-05 | 2.01666465 | 1.011971 |
| ZNF367 | 1.99E-05 | 1.04E-06 | 2.01544065 | 1.011095 |
| CXCL11 | 0.0258 | 0.00791 | 2.01438256 | 1.010338 |
| VCAM1 | 0.00596 | 0.00131 | 2.0137436 | 1.00988 |
| CDK2 | 2.61E-08 | 5.30E-11 | 2.01286609 | 1.009251 |
| FANCI | 1.31E-06 | 2.51E-08 | 2.01225676 | 1.008814 |
| BCL2A1 | 0.00085 | 0.00012 | 2.01223263 | 1.008797 |
| MS4A4A | 4.12E-05 | 2.64E-06 | 2.01222635 | 1.008793 |
| MND1 | 6.06E-06 | 2.17E-07 | 2.00828567 | 1.005965 |
| TK1 | 1.25E-06 | 2.36E-08 | 2.00730633 | 1.005261 |
| VSIG4 | 9.78E-06 | 4.11E-07 | 2.00224095 | 1.001616 |
| GBP3 | 0.000333 | 3.79E-05 | 2.0020962 | 1.001511 |
| FAM60A | 1.84E-07 | 1.34E-09 | 2.00191372 | 1.00138 |
| FKBP5 | 0.00018 | 1.75E-05 | 1.99594034 | 0.997069 |
| RUNX1 | 5.04E-06 | 1.68E-07 | 1.99174907 | 0.994036 |
| TNFAIP6 | 0.000344 | 3.95E-05 | 1.98467255 | 0.988901 |
| PLAC8 | 0.000431 | 5.25E-05 | 1.9832289 | 0.987851 |
| SLC11A1 | 9.66E-06 | 4.04E-07 | 1.98203948 | 0.986986 |
| PIF1 | 1.30E-06 | 2.48E-08 | 1.98157106 | 0.986645 |
| KLHDC8A | 9.53E-05 | 7.77E-06 | 1.98115932 | 0.986345 |
| PTTG3P | 7.43E-06 | 2.87E-07 | 1.97780611 | 0.983901 |
| KIF11 | 4.57E-06 | 1.47E-07 | 1.97721342 | 0.983469 |
| PLA2G2A | 0.0205 | 0.00597 | 1.97702991 | 0.983335 |
| MRC2 | 1.97E-06 | 4.52E-08 | 1.97649979 | 0.982948 |
| LINC00689 | 0.0456 | 0.0161 | 1.97516982 | 0.981977 |
| SMC4 | 4.06E-07 | 4.48E-09 | 1.97344704 | 0.980718 |
| OTP | 0.00184 | 0.000313 | 1.97266066 | 0.980143 |
| PARP4 | 7.48E-05 | 5.69E-06 | 1.96999286 | 0.97819 |
| TLR2 | 9.55E-06 | 3.98E-07 | 1.96818414 | 0.976865 |
| MTHFD2 | 7.91E-08 | 3.80E-10 | 1.96693325 | 0.975948 |
| DNAH11 | 7.00E-08 | 2.99E-10 | 1.96563902 | 0.974998 |
| TIMP4 | 0.000393 | 4.64E-05 | 1.96397586 | 0.973777 |
| PLEKHG2 | 1.19E-08 | 1.45E-11 | 1.96299895 | 0.973059 |
| TMEM45A | 5.86E-06 | 2.06E-07 | 1.95972891 | 0.970654 |
| MUC1 | 8.30E-06 | 3.32E-07 | 1.95897977 | 0.970103 |
| HK2 | 3.75E-05 | 2.33E-06 | 1.95688476 | 0.968559 |
| FKBP10 | 2.70E-08 | 5.78E-11 | 1.95503522 | 0.967195 |

| | | | | |
|---|---|---|---|---|
| GINS2 | 1.74E-07 | 1.26E-09 | 1.95492072 | 0.96711 |
| CENPE | 5.47E-06 | 1.88E-07 | 1.95464512 | 0.966907 |
| HAND2-AS1 | 0.00398 | 8.00E-04 | 1.95288419 | 0.965606 |
| MDFI | 1.23E-05 | 5.57E-07 | 1.95229193 | 0.965169 |
| FAM46A | 1.38E-06 | 2.71E-08 | 1.95219613 | 0.965098 |
| DIAPH3 | 7.67E-07 | 1.18E-08 | 1.95149058 | 0.964577 |
| TIMELESS | 2.19E-07 | 1.77E-09 | 1.95078015 | 0.964051 |
| HIST1H2AM | 1.39E-06 | 2.74E-08 | 1.94850212 | 0.962366 |
| AEBP1 | 0.000122 | 1.05E-05 | 1.94689759 | 0.961177 |
| MIA | 2.00E-04 | 2.00E-05 | 1.9467049 | 0.961034 |
| FAM111B | 0.000108 | 9.06E-06 | 1.94670247 | 0.961032 |
| HIST1H1B | 6.32E-06 | 2.29E-07 | 1.94534698 | 0.960028 |
| HOXA11-AS | 0.0132 | 0.00348 | 1.94513016 | 0.959867 |
| TRPM8 | 0.00452 | 0.000934 | 1.94483182 | 0.959645 |
| TFAP2B | 0.00477 | 0.000999 | 1.94322008 | 0.958449 |
| FMOD | 0.00298 | 0.000562 | 1.93788908 | 0.954486 |
| S100A10 | 0.000204 | 2.05E-05 | 1.9371056 | 0.953903 |
| RNASE3 | 0.000128 | 1.12E-05 | 1.93489101 | 0.952252 |
| DEPDC1B | 2.57E-05 | 1.43E-06 | 1.93356813 | 0.951266 |
| MSN | 4.34E-06 | 1.37E-07 | 1.93244171 | 0.950425 |
| IRX3 | 0.00204 | 0.000353 | 1.9301776 | 0.948734 |
| CLIC1 | 5.61E-07 | 7.25E-09 | 1.92981881 | 0.948465 |
| IGF2BP2 | 0.000765 | 0.000106 | 1.9289724 | 0.947833 |
| MYO1G | 5.02E-05 | 3.40E-06 | 1.92672437 | 0.94615 |
| FBLIM1 | 0.000322 | 3.62E-05 | 1.92547715 | 0.945216 |
| EGFLAM | 0.000461 | 5.69E-05 | 1.92530392 | 0.945086 |
| CKS2 | 1.47E-06 | 2.98E-08 | 1.92480301 | 0.944711 |
| S100A3 | 4.94E-05 | 3.33E-06 | 1.92384411 | 0.943992 |
| FN1 | 2.47E-05 | 1.36E-06 | 1.92376023 | 0.943929 |
| NGFR | 0.00328 | 0.000633 | 1.92328905 | 0.943576 |
| TYMP | 4.91E-05 | 3.30E-06 | 1.92288542 | 0.943273 |
| WISP1 | 0.00408 | 0.000827 | 1.91975885 | 0.940925 |
| RAB27A | 5.48E-05 | 3.83E-06 | 1.91928093 | 0.940566 |
| TYMS | 7.03E-06 | 2.64E-07 | 1.91733828 | 0.939105 |
| SPARC | 2.02E-08 | 3.25E-11 | 1.91532843 | 0.937592 |
| F2RL2 | 8.42E-05 | 6.65E-06 | 1.91461617 | 0.937055 |
| DPYSL3 | 6.30E-07 | 8.63E-09 | 1.9143176 | 0.93683 |
| TNFRSF19 | 2.00E-07 | 1.53E-09 | 1.91183127 | 0.934955 |
| AURKA | 5.18E-07 | 6.42E-09 | 1.90612892 | 0.930646 |
| LPL | 4.17E-06 | 1.29E-07 | 1.90602772 | 0.930569 |
| SAMD9 | 1.14E-05 | 5.04E-07 | 1.90443203 | 0.929361 |
| PHLDA1 | 3.02E-06 | 8.21E-08 | 1.90411564 | 0.929121 |
| MIR210HG | 3.93E-05 | 2.48E-06 | 1.89702738 | 0.923741 |

| | | | | |
|---|---|---|---|---|
| CLEC18B | 0.000197 | 1.95E-05 | 1.89620573 | 0.923116 |
| MARVELD3 | 0.00916 | 0.00221 | 1.89580739 | 0.922812 |
| HAS2 | 0.00119 | 0.000182 | 1.89512722 | 0.922295 |
| FAM20A | 8.56E-05 | 6.80E-06 | 1.89501452 | 0.922209 |
| DPEP1 | 0.00172 | 0.000287 | 1.89484285 | 0.922078 |
| FAM122B | 1.47E-07 | 9.50E-10 | 1.89359933 | 0.921131 |
| NMB | 0.000204 | 2.04E-05 | 1.89216959 | 0.920041 |
| COL5A1 | 0.00985 | 0.00242 | 1.89145716 | 0.919498 |
| NSUN7 | 0.00018 | 1.75E-05 | 1.88848248 | 0.917227 |
| CAPN15 | 0.000223 | 2.28E-05 | 1.88700638 | 0.916099 |
| HOXB6 | 0.0481 | 0.0172 | 1.88666778 | 0.91584 |
| PTTG1 | 4.83E-06 | 1.58E-07 | 1.88486396 | 0.91446 |
| ADAMTS9 | 0.000126 | 1.10E-05 | 1.88441223 | 0.914115 |
| CAV3 | 0.00299 | 0.000564 | 1.88192457 | 0.912209 |
| STK32A | 1.25E-05 | 5.67E-07 | 1.88101975 | 0.911515 |
| ITGA4 | 5.71E-05 | 4.05E-06 | 1.87878464 | 0.9098 |
| HOXC8 | 0.0188 | 0.00535 | 1.87568297 | 0.907416 |
| MAP3K7CL | 3.62E-06 | 1.05E-07 | 1.87327837 | 0.905565 |
| FGFBP2 | 0.0211 | 0.00615 | 1.87266196 | 0.905091 |
| STAB1 | 1.14E-05 | 5.02E-07 | 1.87190575 | 0.904508 |
| CASP4 | 6.06E-06 | 2.16E-07 | 1.87005422 | 0.90308 |
| ABCA13 | 0.00857 | 0.00204 | 1.86650995 | 0.900343 |
| AGMO | 0.0162 | 0.00446 | 1.86552449 | 0.899581 |
| CD48 | 0.00117 | 0.000178 | 1.86495394 | 0.89914 |
| PRPH | 0.000169 | 1.61E-05 | 1.86301592 | 0.89764 |
| MSR1 | 2.16E-05 | 1.15E-06 | 1.86223546 | 0.897036 |
| KLB | 0.000337 | 3.86E-05 | 1.85913083 | 0.894628 |
| NFE2L3 | 0.00437 | 0.000898 | 1.85890598 | 0.894454 |
| HAND2 | 0.00736 | 0.0017 | 1.85668156 | 0.892726 |
| WDR76 | 2.98E-05 | 1.75E-06 | 1.85423924 | 0.890827 |
| CDH24 | 1.08E-05 | 4.66E-07 | 1.84964384 | 0.887248 |
| OSBPL3 | 6.68E-06 | 2.47E-07 | 1.84851558 | 0.886367 |
| PCOLCE | 0.000165 | 1.55E-05 | 1.84685321 | 0.885069 |
| C1orf226 | 4.77E-06 | 1.55E-07 | 1.84646767 | 0.884768 |
| TES | 8.36E-06 | 3.35E-07 | 1.84636336 | 0.884687 |
| SERPINA1 | 0.000329 | 3.74E-05 | 1.84520947 | 0.883785 |
| HK3 | 0.000348 | 4.00E-05 | 1.84428933 | 0.883065 |
| EFR3A | 1.61E-05 | 7.84E-07 | 1.84326207 | 0.882261 |
| FOXJ1 | 0.00329 | 0.000634 | 1.8423254 | 0.881528 |
| S100A11 | 0.00014 | 1.26E-05 | 1.83924517 | 0.879114 |
| BEST3 | 0.00406 | 0.000822 | 1.83878602 | 0.878754 |
| HIST1H3F | 0.000884 | 0.000126 | 1.83723567 | 0.877537 |
| TGFB1I1 | 1.48E-06 | 3.02E-08 | 1.83611243 | 0.876654 |

| | | | | |
|---|---|---|---|---|
| HSPA6 | 0.00297 | 0.000561 | 1.83481715 | 0.875636 |
| C1R | 4.63E-05 | 3.05E-06 | 1.8344045 | 0.875312 |
| HOXB4 | 0.00186 | 0.000317 | 1.83101486 | 0.872644 |
| P2RY8 | 0.000139 | 1.24E-05 | 1.83095077 | 0.872593 |
| MS4A7 | 6.58E-05 | 4.82E-06 | 1.82860339 | 0.870742 |
| FLJ36840 | 2.90E-06 | 7.75E-08 | 1.82850998 | 0.870669 |
| CASP5 | 5.82E-06 | 2.04E-07 | 1.82795645 | 0.870232 |
| CDK6 | 0.000109 | 9.20E-06 | 1.82783419 | 0.870135 |
| LEF1-AS1 | 0.00441 | 0.000908 | 1.82726123 | 0.869683 |
| PYGL | 2.61E-05 | 1.46E-06 | 1.82650019 | 0.869082 |
| RBM47 | 7.95E-06 | 3.12E-07 | 1.82626738 | 0.868898 |
| LOC400043 | 4.94E-06 | 1.64E-07 | 1.8259155 | 0.86862 |
| STON1 | 7.76E-06 | 3.03E-07 | 1.82376482 | 0.86692 |
| IL7 | 0.000246 | 2.58E-05 | 1.82333317 | 0.866578 |
| LIMD1 | 7.76E-06 | 3.04E-07 | 1.82327402 | 0.866531 |
| FBN2 | 0.0327 | 0.0106 | 1.82219064 | 0.865674 |
| PHEX | 2.10E-05 | 1.11E-06 | 1.82001545 | 0.863951 |
| SLC16A3 | 0.000121 | 1.05E-05 | 1.81976594 | 0.863753 |
| SLC43A3 | 2.84E-06 | 7.46E-08 | 1.81976241 | 0.86375 |
| KRTAP8-1 | 0.000706 | 9.57E-05 | 1.81889051 | 0.863059 |
| CDC20 | 8.15E-06 | 3.24E-07 | 1.81822924 | 0.862534 |
| FANCA | 5.07E-06 | 1.69E-07 | 1.817053 | 0.861601 |
| HIST1H4J | 1.34E-06 | 2.59E-08 | 1.81701307 | 0.861569 |
| MGP | 0.0066 | 0.00149 | 1.81684079 | 0.861432 |
| LRR1 | 8.86E-07 | 1.48E-08 | 1.81683462 | 0.861427 |
| NID2 | 0.000105 | 8.76E-06 | 1.81558101 | 0.860431 |
| SIGLEC9 | 2.56E-05 | 1.42E-06 | 1.81537236 | 0.860266 |
| XRCC2 | 7.23E-05 | 5.45E-06 | 1.81481149 | 0.85982 |
| RAB13 | 6.60E-07 | 9.31E-09 | 1.81455753 | 0.859618 |
| KYNU | 0.00043 | 5.22E-05 | 1.81344099 | 0.85873 |
| SP140L | 2.02E-05 | 1.06E-06 | 1.81258757 | 0.858051 |
| CDH15 | 0.015 | 0.00407 | 1.8116881 | 0.857335 |
| PKMYT1 | 2.97E-07 | 2.76E-09 | 1.81114506 | 0.856902 |
| LOXL1 | 0.00235 | 0.00042 | 1.80970708 | 0.855756 |
| COL14A1 | 0.0102 | 0.00253 | 1.80851492 | 0.854806 |
| MFAP2 | 0.00111 | 0.000167 | 1.80827776 | 0.854616 |
| DRAM1 | 1.70E-05 | 8.44E-07 | 1.80481542 | 0.851851 |
| CLIC4 | 2.80E-05 | 1.61E-06 | 1.80481442 | 0.851851 |
| TEAD4 | 1.14E-05 | 5.04E-07 | 1.8025073 | 0.850005 |
| NAMPT | 0.000517 | 6.55E-05 | 1.79916774 | 0.84733 |
| METTL21B | 2.16E-05 | 1.15E-06 | 1.79904528 | 0.847232 |
| CIITA | 0.000932 | 0.000135 | 1.79736934 | 0.845887 |
| IL21R | 0.00245 | 0.000443 | 1.7963539 | 0.845072 |

| | | | | |
|---|---|---|---|---|
| CD300LF | 6.58E-05 | 4.82E-06 | 1.79595151 | 0.844748 |
| ZNF300 | 9.70E-05 | 7.95E-06 | 1.79531538 | 0.844237 |
| NLRC5 | 5.92E-05 | 4.23E-06 | 1.79525179 | 0.844186 |
| MMS22L | 1.91E-07 | 1.41E-09 | 1.79505706 | 0.84403 |
| PROS1 | 8.59E-06 | 3.49E-07 | 1.79399516 | 0.843176 |
| CLEC5A | 0.00154 | 0.00025 | 1.79356173 | 0.842827 |
| MCM2 | 1.88E-06 | 4.29E-08 | 1.79261975 | 0.84207 |
| TMEM255A | 0.000106 | 8.86E-06 | 1.79184855 | 0.841449 |
| C4orf47 | 2.19E-06 | 5.20E-08 | 1.79173317 | 0.841356 |
| CNN2 | 0.00198 | 0.000341 | 1.78886324 | 0.839043 |
| SEC61G | 0.0141 | 0.00379 | 1.78853742 | 0.83878 |
| FCGR3B | 0.000544 | 6.95E-05 | 1.78823879 | 0.838539 |
| SOX4 | 0.00107 | 0.00016 | 1.78641739 | 0.837069 |
| C1QB | 6.23E-05 | 4.52E-06 | 1.78634916 | 0.837014 |
| HIST1H3J | 0.00198 | 0.00034 | 1.78623847 | 0.836925 |
| C8orf88 | 3.24E-07 | 3.17E-09 | 1.78447811 | 0.835502 |
| ITGB3 | 0.00145 | 0.000231 | 1.78427761 | 0.83534 |
| RGS1 | 0.000174 | 1.66E-05 | 1.78319763 | 0.834467 |
| EMP3 | 0.00117 | 0.000177 | 1.78159795 | 0.833172 |
| FLNA | 2.59E-06 | 6.59E-08 | 1.78150212 | 0.833094 |
| PLAUR | 0.00127 | 0.000197 | 1.78135086 | 0.832972 |
| OXTR | 0.000419 | 5.04E-05 | 1.78081938 | 0.832541 |
| SNORA62///RPSA | 0.0497 | 0.0179 | 1.78020823 | 0.832046 |
| FAM129A | 5.53E-05 | 3.88E-06 | 1.77934763 | 0.831348 |
| TEAD2 | 2.11E-05 | 1.11E-06 | 1.77891725 | 0.830999 |
| SYNE2 | 1.37E-06 | 2.67E-08 | 1.77880726 | 0.83091 |
| LOXL3 | 1.35E-06 | 2.63E-08 | 1.77839315 | 0.830574 |
| CTSS | 0.00064 | 8.48E-05 | 1.77718775 | 0.829596 |
| PEX14 | 3.26E-06 | 9.18E-08 | 1.77689324 | 0.829357 |
| CSRP2 | 3.12E-05 | 1.86E-06 | 1.77605911 | 0.82868 |
| SDC1 | 0.00217 | 0.000381 | 1.77521345 | 0.827993 |
| RAD51AP1 | 2.42E-06 | 6.00E-08 | 1.77435121 | 0.827292 |
| SPON2 | 0.00119 | 0.000181 | 1.77381789 | 0.826858 |
| ADAMTS6 | 0.000852 | 0.000121 | 1.7733999 | 0.826518 |
| MDK | 0.000153 | 1.41E-05 | 1.77269791 | 0.825947 |
| PLVAP | 0.000142 | 1.28E-05 | 1.7723722 | 0.825682 |
| GBP5 | 0.00592 | 0.0013 | 1.76974193 | 0.823539 |
| NID1 | 3.71E-06 | 1.08E-07 | 1.76758979 | 0.821784 |
| C1QTNF6 | 4.48E-05 | 2.93E-06 | 1.7674119 | 0.821638 |
| TP53I3 | 5.97E-07 | 7.99E-09 | 1.76289824 | 0.817949 |
| GALNT5 | 0.00587 | 0.00129 | 1.76278069 | 0.817853 |
| CDCA3 | 4.31E-06 | 1.36E-07 | 1.76267036 | 0.817763 |
| PABPC1L | 1.69E-05 | 8.40E-07 | 1.76248796 | 0.817613 |

| | | | | |
|---|---|---|---|---|
| LIMA1 | 1.07E-06 | 1.88E-08 | 1.76093566 | 0.816342 |
| TAGLN2 | 4.37E-06 | 1.38E-07 | 1.75966842 | 0.815304 |
| PRKD3 | 1.39E-08 | 1.96E-11 | 1.75901941 | 0.814771 |
| APOC1 | 7.22E-06 | 2.75E-07 | 1.75896759 | 0.814729 |
| HOXD8 | 0.0366 | 0.0122 | 1.75728501 | 0.813348 |
| POM121L9P | 0.000875 | 0.000125 | 1.75537176 | 0.811777 |
| STIL | 2.87E-06 | 7.59E-08 | 1.75386342 | 0.810536 |
| RCC1 | 2.05E-07 | 1.59E-09 | 1.75311155 | 0.809918 |
| POFUT1 | 0.00718 | 0.00165 | 1.75255777 | 0.809462 |
| TCF3 | 2.81E-07 | 2.54E-09 | 1.75248282 | 0.8094 |
| C6orf118 | 0.0122 | 0.00314 | 1.75163308 | 0.808701 |
| HMGA2 | 0.0441 | 0.0154 | 1.75159168 | 0.808667 |
| CENPI | 4.02E-05 | 2.54E-06 | 1.75027353 | 0.80758 |
| ZNF107 | 7.81E-07 | 1.22E-08 | 1.74894097 | 0.806482 |
| ZIC1 | 7.33E-05 | 5.54E-06 | 1.74890509 | 0.806452 |
| RBL1 | 1.06E-07 | 5.84E-10 | 1.74668494 | 0.804619 |
| TNFRSF10B | 0.000318 | 3.57E-05 | 1.74647973 | 0.80445 |
| FAM111A | 1.17E-07 | 6.80E-10 | 1.74615327 | 0.80418 |
| NEXN | 2.63E-05 | 1.48E-06 | 1.74604023 | 0.804087 |
| CDCP1 | 0.00346 | 0.000673 | 1.74552522 | 0.803661 |
| PZP | 0.00583 | 0.00128 | 1.74438235 | 0.802716 |
| DDX60L | 3.63E-05 | 2.25E-06 | 1.74351177 | 0.801996 |
| SMIM3 | 0.000173 | 1.65E-05 | 1.74085507 | 0.799796 |
| ETV4 | 6.00E-04 | 7.85E-05 | 1.74011192 | 0.79918 |
| GPC2 | 0.00225 | 0.000399 | 1.73975506 | 0.798884 |
| THOC2 | 3.89E-08 | 1.20E-10 | 1.73973624 | 0.798869 |
| PRIM2 | 3.11E-08 | 7.56E-11 | 1.73944541 | 0.798627 |
| C5AR1 | 9.00E-04 | 0.000129 | 1.73909061 | 0.798333 |
| FCGR3A | 0.000136 | 1.22E-05 | 1.73879349 | 0.798087 |
| CASP6 | 5.46E-08 | 1.97E-10 | 1.73832942 | 0.797702 |
| HLA-DQA2 | 0.0254 | 0.00777 | 1.73550497 | 0.795356 |
| LMAN1 | 4.52E-05 | 2.97E-06 | 1.73482459 | 0.79479 |
| SPATA17 | 0.000136 | 1.21E-05 | 1.73428692 | 0.794343 |
| KDELR3 | 0.00117 | 0.000178 | 1.73374606 | 0.793893 |
| SPP1 | 3.10E-05 | 1.84E-06 | 1.73347484 | 0.793667 |
| PPP1R3B | 0.000533 | 6.78E-05 | 1.73260838 | 0.792946 |
| C1QTNF1 | 0.00219 | 0.000386 | 1.7322198 | 0.792622 |
| F13A1 | 0.0327 | 0.0106 | 1.7321151 | 0.792535 |
| SH2D4A | 0.000223 | 2.29E-05 | 1.73139644 | 0.791936 |
| ECSCR | 2.24E-05 | 1.19E-06 | 1.73123372 | 0.791801 |
| GAPT | 0.00287 | 0.000536 | 1.73074478 | 0.791393 |
| SOD2 | 0.000156 | 1.45E-05 | 1.73055705 | 0.791237 |
| CCND2 | 0.00048 | 5.98E-05 | 1.73035698 | 0.79107 |

| | | | | |
|---|---|---|---|---|
| GINS4 | 6.57E-06 | 2.42E-07 | 1.72972825 | 0.790545 |
| KCNE4 | 0.000169 | 1.61E-05 | 1.72907302 | 0.789999 |
| FANCC | 0.0395 | 0.0134 | 1.72858686 | 0.789593 |
| TGIF1 | 1.96E-06 | 4.49E-08 | 1.72767889 | 0.788835 |
| ACE | 0.000686 | 9.24E-05 | 1.72674016 | 0.788051 |
| SLFN12 | 7.69E-05 | 5.90E-06 | 1.72580696 | 0.787271 |
| CENPL | 7.63E-07 | 1.17E-08 | 1.72577203 | 0.787242 |
| FBXO5 | 6.73E-06 | 2.49E-07 | 1.72520775 | 0.78677 |
| ATAD5 | 2.40E-05 | 1.31E-06 | 1.72504536 | 0.786634 |
| CHRNA5 | 0.000166 | 1.57E-05 | 1.72468609 | 0.786334 |
| ITPRIPL1 | 1.55E-05 | 7.52E-07 | 1.72186699 | 0.783974 |
| MS4A14 | 0.000227 | 2.34E-05 | 1.7217628 | 0.783886 |
| TP53 | 4.59E-06 | 1.48E-07 | 1.72135195 | 0.783542 |
| HLA-DPB2 | 2.00E-04 | 1.99E-05 | 1.72082489 | 0.7831 |
| NEAT1 | 0.00121 | 0.000185 | 1.72057443 | 0.78289 |
| C8orf4 | 0.000744 | 0.000102 | 1.71979452 | 0.782236 |
| CARD17 | 0.000365 | 4.26E-05 | 1.71787909 | 0.780629 |
| TMEM100 | 0.0139 | 0.00371 | 1.7177699 | 0.780537 |
| MYOF | 0.000183 | 1.78E-05 | 1.71700495 | 0.779894 |
| CD99P1 | 7.08E-06 | 2.68E-07 | 1.71686845 | 0.77978 |
| TNFAIP3 | 0.00231 | 0.00041 | 1.7157962 | 0.778878 |
| HLA-L | 6.89E-05 | 5.12E-06 | 1.71562079 | 0.778731 |
| SOAT1 | 1.40E-06 | 2.77E-08 | 1.71555312 | 0.778674 |
| ETS1 | 3.40E-06 | 9.66E-08 | 1.71508348 | 0.778279 |
| USP30-AS1 | 0.00486 | 0.00102 | 1.71495402 | 0.77817 |
| SLPI | 0.00317 | 0.000606 | 1.71468635 | 0.777945 |
| IL4I1 | 0.000397 | 4.70E-05 | 1.71375277 | 0.777159 |
| PTTG2 | 4.99E-05 | 3.37E-06 | 1.71297572 | 0.776505 |
| CALCRL | 0.00163 | 0.000268 | 1.71279929 | 0.776356 |
| GLI2 | 0.000292 | 3.21E-05 | 1.71240921 | 0.776028 |
| LOC100128922 | 0.000189 | 1.86E-05 | 1.71212247 | 0.775786 |
| IGFBP1 | 0.00872 | 0.00208 | 1.71186271 | 0.775567 |
| FOXD1 | 0.000387 | 4.56E-05 | 1.71176743 | 0.775487 |
| SPINK1 | 0.0262 | 0.00807 | 1.71139182 | 0.77517 |
| C17orf53 | 3.12E-06 | 8.61E-08 | 1.71068651 | 0.774575 |
| MICB | 0.00031 | 3.46E-05 | 1.70996631 | 0.773968 |
| PTPRZ1 | 8.13E-06 | 3.21E-07 | 1.70984708 | 0.773867 |
| BRIP1 | 0.00181 | 0.000305 | 1.709773 | 0.773805 |
| DZIP1L | 3.76E-07 | 4.00E-09 | 1.70958304 | 0.773645 |
| HIST1H1D | 7.45E-07 | 1.13E-08 | 1.70892964 | 0.773093 |
| POLE2 | 1.74E-05 | 8.73E-07 | 1.70638076 | 0.77094 |
| HIST1H3H | 0.000143 | 1.30E-05 | 1.70614552 | 0.770741 |
| CSPG4 | 8.20E-05 | 6.42E-06 | 1.70612045 | 0.77072 |

| | | | | |
|---|---|---|---|---|
| EME1 | 8.51E-05 | 6.75E-06 | 1.70606226 | 0.77067 |
| LAMC1 | 2.72E-07 | 2.39E-09 | 1.70570825 | 0.770371 |
| SYCE2 | 0.000182 | 1.77E-05 | 1.70491664 | 0.769701 |
| PPP1R1C | 0.0362 | 0.012 | 1.70373127 | 0.768698 |
| LTBP2 | 0.0141 | 0.00378 | 1.70266357 | 0.767793 |
| PLAT | 0.00104 | 0.000155 | 1.70238979 | 0.767561 |
| CDCA7L | 7.47E-07 | 1.14E-08 | 1.70030788 | 0.765796 |
| NAPSA | 0.000894 | 0.000128 | 1.69972565 | 0.765302 |
| SP100 | 4.19E-06 | 1.31E-07 | 1.69938413 | 0.765012 |
| TNFRSF11B | 0.00048 | 5.98E-05 | 1.69680997 | 0.762825 |
| HOXB7 | 0.000121 | 1.04E-05 | 1.69620389 | 0.76231 |
| CEP152 | 2.86E-06 | 7.56E-08 | 1.69590435 | 0.762055 |
| CHST9 | 0.0254 | 0.00777 | 1.69548486 | 0.761698 |
| CHEK2 | 3.12E-06 | 8.60E-08 | 1.69531799 | 0.761556 |
| TRIM14 | 1.57E-05 | 7.59E-07 | 1.69520107 | 0.761456 |
| CYTL1 | 0.00049 | 6.15E-05 | 1.69441234 | 0.760785 |
| RP2 | 7.91E-08 | 3.76E-10 | 1.69424358 | 0.760641 |
| ATL3 | 1.60E-07 | 1.11E-09 | 1.69326961 | 0.759812 |
| CXCR4 | 0.000791 | 0.00011 | 1.69287248 | 0.759473 |
| CD99 | 0.000254 | 2.69E-05 | 1.69276558 | 0.759382 |
| SIGLEC1 | 0.0205 | 0.00597 | 1.69223239 | 0.758928 |
| AQP1 | 0.0384 | 0.013 | 1.69128103 | 0.758116 |
| MIR155HG | 0.000263 | 2.81E-05 | 1.68965348 | 0.756727 |
| ID3 | 2.73E-05 | 1.56E-06 | 1.68929455 | 0.756421 |
| EMILIN2 | 0.00222 | 0.000392 | 1.68792418 | 0.75525 |
| HMOX1 | 0.000838 | 0.000118 | 1.68789867 | 0.755228 |
| DRAXIN | 0.000483 | 6.03E-05 | 1.68771196 | 0.755069 |
| SOCS3 | 0.00256 | 0.000466 | 1.68692625 | 0.754397 |
| LINC01224 | 0.00092 | 0.000132 | 1.68665172 | 0.754162 |
| GINS1 | 4.11E-06 | 1.27E-07 | 1.68655761 | 0.754082 |
| LOC105369230///HLA-DRB5///HLA-DRB4///HLA-DRB3///HLA-DRB1 | 0.00479 | 0.00101 | 1.68629039 | 0.753853 |
| PDLIM7 | 0.000474 | 5.89E-05 | 1.68622085 | 0.753794 |
| MEST | 0.000696 | 9.41E-05 | 1.68584979 | 0.753476 |
| SAMD9L | 0.000351 | 4.05E-05 | 1.6858117 | 0.753443 |
| IFI16 | 3.12E-05 | 1.86E-06 | 1.68548244 | 0.753162 |
| CARD16 | 0.000359 | 4.15E-05 | 1.68525441 | 0.752966 |
| TOB2P1 | 1.89E-06 | 4.31E-08 | 1.68492853 | 0.752687 |
| KNTC1 | 8.83E-07 | 1.48E-08 | 1.68445128 | 0.752279 |
| BRCA2 | 2.76E-05 | 1.58E-06 | 1.68235341 | 0.750481 |
| CALD1 | 9.54E-07 | 1.62E-08 | 1.68232087 | 0.750453 |

| | | | | |
|---|---|---|---|---|
| GZMA | 0.00973 | 0.00239 | 1.68225172 | 0.750394 |
| TRIM49 | 0.0406 | 0.0139 | 0.5945921 | -0.75003 |
| EIF4A2 | 7.14E-08 | 3.16E-10 | 0.59435686 | -0.7506 |
| GAD1 | 0.00149 | 0.00024 | 0.59402024 | -0.75142 |
| CHSY3 | 0.00283 | 0.000526 | 0.59361918 | -0.75239 |
| STX1B | 0.00031 | 3.45E-05 | 0.59330379 | -0.75316 |
| NEBL | 0.00448 | 0.000926 | 0.59291886 | -0.75409 |
| KL | 0.0012 | 0.000184 | 0.5929159 | -0.7541 |
| SCAI | 4.40E-06 | 1.39E-07 | 0.5924314 | -0.75528 |
| HCRTR2 | 0.00137 | 0.000216 | 0.59242737 | -0.75529 |
| ADGRA1 | 0.00491 | 0.00104 | 0.59237292 | -0.75542 |
| GUCY1A3 | 0.0017 | 0.000283 | 0.59229541 | -0.75561 |
| HBA2 | 0.00521 | 0.00112 | 0.59189132 | -0.7566 |
| XKR4 | 0.015 | 0.00408 | 0.59184406 | -0.75671 |
| DTNB | 2.01E-08 | 3.19E-11 | 0.59176841 | -0.7569 |
| LOC283454 | 0.022 | 0.0065 | 0.59173712 | -0.75697 |
| EMX2 | 0.0219 | 0.00646 | 0.5916238 | -0.75725 |
| PRRT2 | 0.000166 | 1.57E-05 | 0.59160744 | -0.75729 |
| DLGAP1 | 0.00428 | 0.000876 | 0.59153712 | -0.75746 |
| EFHD1 | 0.0062 | 0.00137 | 0.59119518 | -0.75829 |
| ADGRB3 | 0.00105 | 0.000156 | 0.59083279 | -0.75918 |
| HID1 | 0.000139 | 1.24E-05 | 0.59082173 | -0.75921 |
| LAMP5 | 0.000422 | 5.10E-05 | 0.59062274 | -0.75969 |
| LINC00473 | 0.0343 | 0.0113 | 0.59057419 | -0.75981 |
| TSHZ2 | 5.86E-06 | 2.06E-07 | 0.59023566 | -0.76064 |
| NTNG2 | 0.000866 | 0.000123 | 0.59022985 | -0.76065 |
| CPEB1 | 1.43E-06 | 2.86E-08 | 0.59022135 | -0.76067 |
| LOC100506725 | 3.90E-06 | 1.17E-07 | 0.59016759 | -0.7608 |
| NCALD | 0.00181 | 0.000305 | 0.59010889 | -0.76095 |
| AMN1 | 2.59E-06 | 6.58E-08 | 0.59009155 | -0.76099 |
| DUSP8 | 4.10E-06 | 1.25E-07 | 0.58998407 | -0.76125 |
| BCL11B | 0.00623 | 0.00139 | 0.58994996 | -0.76134 |
| SLC22A15 | 0.000715 | 9.73E-05 | 0.58982342 | -0.76164 |
| ARHGEF4 | 0.000642 | 8.51E-05 | 0.5897045 | -0.76194 |
| BOK | 0.000223 | 2.28E-05 | 0.58948001 | -0.76249 |
| KCNAB2 | 1.98E-05 | 1.03E-06 | 0.58934454 | -0.76282 |
| LARGE1 | 1.01E-05 | 4.31E-07 | 0.58912942 | -0.76334 |
| PDE4A | 1.91E-05 | 9.91E-07 | 0.58885862 | -0.76401 |
| SHTN1 | 8.23E-05 | 6.45E-06 | 0.58862801 | -0.76457 |
| CABYR | 0.000181 | 1.76E-05 | 0.58827874 | -0.76543 |
| LOC105378732 | 9.78E-06 | 4.11E-07 | 0.58824025 | -0.76552 |
| NKAPL | 3.60E-05 | 2.22E-06 | 0.58819972 | -0.76562 |
| CDKN2D | 2.57E-07 | 2.16E-09 | 0.58815565 | -0.76573 |

| | | | | |
|---|---|---|---|---|
| PALM | 0.000581 | 7.54E-05 | 0.58813477 | -0.76578 |
| SLC39A10 | 3.82E-07 | 4.14E-09 | 0.58797259 | -0.76618 |
| BTRC | 3.56E-08 | 1.02E-10 | 0.58780409 | -0.76659 |
| MEG3 | 0.00458 | 0.00095 | 0.58770452 | -0.76684 |
| NIPA1 | 2.26E-06 | 5.49E-08 | 0.58737843 | -0.76764 |
| NINJ2 | 0.000193 | 1.90E-05 | 0.58715951 | -0.76818 |
| STMN4 | 0.0135 | 0.00357 | 0.5870909 | -0.76834 |
| PRDM12 | 0.000565 | 7.30E-05 | 0.58696643 | -0.76865 |
| PNLDC1 | 0.000552 | 7.08E-05 | 0.58671684 | -0.76926 |
| PTPN4 | 7.99E-07 | 1.26E-08 | 0.58671082 | -0.76928 |
| CCL3 | 0.0106 | 0.00265 | 0.58647524 | -0.76986 |
| RAB37 | 2.54E-05 | 1.41E-06 | 0.58627555 | -0.77035 |
| KLHL14 | 0.00173 | 0.000288 | 0.58621842 | -0.77049 |
| PITPNM2 | 1.60E-06 | 3.35E-08 | 0.58596736 | -0.77111 |
| CMTM4 | 4.78E-05 | 3.18E-06 | 0.58596053 | -0.77112 |
| OSBPL1A | 1.23E-05 | 5.57E-07 | 0.58573317 | -0.77168 |
| PLLP | 0.000507 | 6.39E-05 | 0.58554027 | -0.77216 |
| KCNIP3 | 0.00192 | 0.000328 | 0.58533255 | -0.77267 |
| SMG8 | 0.00119 | 0.000181 | 0.5852119 | -0.77297 |
| PLK5 | 0.00262 | 0.000479 | 0.58521117 | -0.77297 |
| TPD52L1 | 0.0174 | 0.00487 | 0.58508089 | -0.77329 |
| TDGF1 | 0.000675 | 9.06E-05 | 0.58504123 | -0.77339 |
| CCL4 | 0.00189 | 0.000323 | 0.58500003 | -0.77349 |
| ADD2 | 4.79E-06 | 1.57E-07 | 0.58499537 | -0.7735 |
| PTK2B | 1.80E-05 | 9.15E-07 | 0.58446409 | -0.77481 |
| PAH | 0.000195 | 1.92E-05 | 0.58439191 | -0.77499 |
| PCBP3 | 0.000439 | 5.35E-05 | 0.58418969 | -0.77549 |
| FRMD4A | 0.00338 | 0.000655 | 0.58417685 | -0.77552 |
| PPP3CA | 9.49E-08 | 4.94E-10 | 0.58358734 | -0.77698 |
| TEX29 | 1.31E-06 | 2.51E-08 | 0.58347595 | -0.77725 |
| TSPAN7 | 0.000123 | 1.07E-05 | 0.58331489 | -0.77765 |
| MAP3K10 | 6.01E-07 | 8.12E-09 | 0.58317638 | -0.778 |
| PSD3 | 4.99E-06 | 1.66E-07 | 0.58260715 | -0.7794 |
| SLIT3 | 0.00075 | 0.000103 | 0.5823169 | -0.78012 |
| MAP7 | 0.000221 | 2.26E-05 | 0.58205174 | -0.78078 |
| MCTP1 | 7.26E-05 | 5.48E-06 | 0.58184731 | -0.78129 |
| PRKAR2B | 0.000593 | 7.74E-05 | 0.58150331 | -0.78214 |
| CRY2 | 1.14E-06 | 2.07E-08 | 0.58149633 | -0.78216 |
| ELOVL4 | 2.90E-06 | 7.78E-08 | 0.58145176 | -0.78227 |
| ZBTB18 | 0.00043 | 5.22E-05 | 0.58133811 | -0.78255 |
| MAPRE3 | 1.07E-05 | 4.61E-07 | 0.58131228 | -0.78261 |
| BTBD16 | 7.91E-05 | 6.10E-06 | 0.58089753 | -0.78364 |
| ELMO1 | 0.00369 | 0.000728 | 0.58076511 | -0.78397 |

| | | | | |
|---|---|---|---|---|
| SCRT1 | 1.45E-05 | 6.87E-07 | 0.58068425 | -0.78417 |
| PDP1 | 2.04E-06 | 4.76E-08 | 0.58054677 | -0.78452 |
| C4orf45 | 2.55E-06 | 6.43E-08 | 0.58035662 | -0.78499 |
| RND1 | 2.65E-05 | 1.50E-06 | 0.58034258 | -0.78502 |
| NPPC | 0.00179 | 0.000301 | 0.58032283 | -0.78507 |
| NAV2 | 0.00165 | 0.000272 | 0.58009975 | -0.78563 |
| PPFIA3 | 1.41E-05 | 6.60E-07 | 0.57966677 | -0.7867 |
| BOD1L1 | 0.000235 | 2.44E-05 | 0.57955951 | -0.78697 |
| INPP4A | 4.57E-08 | 1.54E-10 | 0.57939635 | -0.78738 |
| FBXO27 | 0.00201 | 0.000346 | 0.57923163 | -0.78779 |
| GPR158 | 0.00357 | 7.00E-04 | 0.57920229 | -0.78786 |
| LPA | 3.77E-05 | 2.35E-06 | 0.57907576 | -0.78818 |
| DLG4 | 1.42E-06 | 2.82E-08 | 0.578896 | -0.78862 |
| NAP1L5 | 2.52E-06 | 6.36E-08 | 0.5787819 | -0.78891 |
| ZDHHC22 | 0.0087 | 0.00208 | 0.57849969 | -0.78961 |
| DBP | 6.91E-07 | 9.90E-09 | 0.57848799 | -0.78964 |
| EFNA3 | 8.69E-06 | 3.54E-07 | 0.57837264 | -0.78993 |
| IQSEC1 | 2.09E-08 | 3.51E-11 | 0.5782216 | -0.79031 |
| ENHO | 0.000211 | 2.13E-05 | 0.5781802 | -0.79041 |
| PLPPR4 | 0.000107 | 8.95E-06 | 0.57815735 | -0.79047 |
| UNC79 | 0.000866 | 0.000123 | 0.57811047 | -0.79058 |
| GAS7 | 0.000769 | 0.000107 | 0.57785475 | -0.79122 |
| C12orf54 | 0.00035 | 4.04E-05 | 0.57775871 | -0.79146 |
| LRP11 | 1.24E-06 | 2.33E-08 | 0.57774845 | -0.79149 |
| GDF7 | 0.000486 | 6.08E-05 | 0.57711515 | -0.79307 |
| BSCL2 | 6.92E-06 | 2.59E-07 | 0.57699159 | -0.79338 |
| SYNDIG1L | 0.0168 | 0.00466 | 0.57693656 | -0.79352 |
| SNTG2 | 0.0372 | 0.0125 | 0.5766509 | -0.79423 |
| ZNF483 | 0.000123 | 1.06E-05 | 0.57652221 | -0.79455 |
| KCNG1 | 0.0101 | 0.0025 | 0.57613396 | -0.79552 |
| LINC01128 | 5.90E-07 | 7.82E-09 | 0.57608161 | -0.79565 |
| HHIP | 0.000274 | 2.97E-05 | 0.57597101 | -0.79593 |
| MCF2L2 | 0.000108 | 9.06E-06 | 0.57596789 | -0.79594 |
| ARHGEF9 | 1.04E-06 | 1.80E-08 | 0.57574505 | -0.7965 |
| FAM174B | 1.69E-06 | 3.64E-08 | 0.57556158 | -0.79696 |
| CDH13 | 0.00622 | 0.00138 | 0.57524247 | -0.79776 |
| NBEA | 0.0016 | 0.000263 | 0.57516596 | -0.79795 |
| ELAVL4 | 3.86E-05 | 2.42E-06 | 0.57514866 | -0.79799 |
| PRODH | 0.00634 | 0.00142 | 0.57502692 | -0.7983 |
| BEGAIN | 6.83E-05 | 5.05E-06 | 0.57501432 | -0.79833 |
| MTMR7 | 4.74E-05 | 3.14E-06 | 0.57463836 | -0.79927 |
| CAMK2N1 | 3.67E-07 | 3.85E-09 | 0.57428935 | -0.80015 |
| BCL2L2 | 2.72E-07 | 2.33E-09 | 0.57420265 | -0.80037 |

| | | | | |
|---|---|---|---|---|
| P2RX6P | 0.000764 | 0.000106 | 0.57335159 | -0.80251 |
| RPS6KA5 | 6.40E-06 | 2.34E-07 | 0.5733506 | -0.80251 |
| SLC6A20 | 0.0106 | 0.00265 | 0.57334396 | -0.80253 |
| FAM65B | 4.66E-05 | 3.08E-06 | 0.57310937 | -0.80312 |
| RAPGEF5 | 0.000381 | 4.47E-05 | 0.57286103 | -0.80374 |
| SCAMP5 | 1.14E-06 | 2.08E-08 | 0.57281866 | -0.80385 |
| PRSS1 | 1.80E-05 | 9.10E-07 | 0.572779 | -0.80395 |
| CLCN4 | 3.24E-06 | 9.06E-08 | 0.57274962 | -0.80402 |
| IGSF8 | 4.27E-07 | 4.86E-09 | 0.57266876 | -0.80423 |
| CYP2E1 | 1.95E-06 | 4.48E-08 | 0.57247262 | -0.80472 |
| GREM1 | 0.0213 | 0.00622 | 0.57229075 | -0.80518 |
| OPRL1 | 3.32E-07 | 3.32E-09 | 0.57198848 | -0.80594 |
| KLK8 | 4.61E-05 | 3.04E-06 | 0.57195304 | -0.80603 |
| FXYD4 | 3.74E-07 | 3.97E-09 | 0.57194983 | -0.80604 |
| ABCC8 | 0.0196 | 0.00563 | 0.57191902 | -0.80612 |
| MAP3K9 | 9.48E-07 | 1.61E-08 | 0.57185433 | -0.80628 |
| CAPN3 | 1.54E-05 | 7.40E-07 | 0.57168558 | -0.80671 |
| DNAJC12 | 5.36E-05 | 3.71E-06 | 0.57109581 | -0.8082 |
| LRP2 | 0.000322 | 3.63E-05 | 0.57088885 | -0.80872 |
| RNF41 | 2.17E-07 | 1.74E-09 | 0.5707031 | -0.80919 |
| FLT3 | 0.00204 | 0.000353 | 0.57064566 | -0.80933 |
| ANKRD33B | 0.00337 | 0.000652 | 0.57055683 | -0.80956 |
| RNF175 | 0.0019 | 0.000324 | 0.57039364 | -0.80997 |
| IQCF4 | 3.02E-05 | 1.77E-06 | 0.57003792 | -0.81087 |
| TMEM191B | 6.74E-07 | 9.55E-09 | 0.5696981 | -0.81173 |
| SYTL2 | 1.53E-05 | 7.31E-07 | 0.56924811 | -0.81287 |
| CAP2 | 2.12E-06 | 5.02E-08 | 0.56903129 | -0.81342 |
| CAMKK2 | 2.28E-09 | 9.44E-13 | 0.56901946 | -0.81345 |
| PPM1H | 7.89E-07 | 1.23E-08 | 0.56898945 | -0.81353 |
| FADS6 | 4.82E-05 | 3.22E-06 | 0.56898637 | -0.81353 |
| KY | 0.000119 | 1.03E-05 | 0.56896764 | -0.81358 |
| KIAA1456 | 2.00E-05 | 1.05E-06 | 0.56888411 | -0.81379 |
| ANKRD29 | 0.000173 | 1.65E-05 | 0.5688383 | -0.81391 |
| BTBD9 | 2.11E-07 | 1.67E-09 | 0.5686199 | -0.81446 |
| KCNA5 | 0.0129 | 0.00338 | 0.56814324 | -0.81567 |
| FAM71E1 | 2.95E-05 | 1.72E-06 | 0.56795211 | -0.81616 |
| IQSEC2 | 6.13E-05 | 4.43E-06 | 0.56785768 | -0.8164 |
| NDRG2 | 0.000874 | 0.000125 | 0.56783753 | -0.81645 |
| RS1 | 0.00187 | 0.000319 | 0.56759221 | -0.81707 |
| PLCB1 | 1.90E-05 | 9.80E-07 | 0.56755067 | -0.81718 |
| ROGDI | 1.19E-08 | 1.44E-11 | 0.56733631 | -0.81772 |
| PLEKHA6 | 0.00224 | 0.000397 | 0.5673273 | -0.81775 |
| CYGB | 4.87E-06 | 1.61E-07 | 0.56732015 | -0.81777 |

| | | | | |
|---|---|---|---|---|
| ZCCHC18 | 1.49E-05 | 7.08E-07 | 0.5671212 | -0.81827 |
| GNAO1 | 1.90E-05 | 9.81E-07 | 0.56692724 | -0.81876 |
| KLHL34 | 0.000114 | 9.79E-06 | 0.56632413 | -0.8203 |
| CDK5R2 | 3.49E-07 | 3.59E-09 | 0.56627008 | -0.82044 |
| LOC157562 | 2.79E-07 | 2.48E-09 | 0.56617506 | -0.82068 |
| WASF1 | 5.92E-06 | 2.09E-07 | 0.56614633 | -0.82075 |
| LINC00950 | 0.000314 | 3.51E-05 | 0.56599386 | -0.82114 |
| KIF1A | 0.000226 | 2.33E-05 | 0.56566425 | -0.82198 |
| CACNB1 | 5.60E-07 | 7.22E-09 | 0.56523139 | -0.82309 |
| TSPAN19 | 0.000838 | 0.000118 | 0.5649746 | -0.82374 |
| CNTNAP5 | 0.00723 | 0.00166 | 0.56493328 | -0.82385 |
| CACNB3 | 1.28E-07 | 7.81E-10 | 0.56481997 | -0.82414 |
| TBC1D26 | 5.11E-07 | 6.27E-09 | 0.56471005 | -0.82442 |
| CORO2A | 0.000517 | 6.55E-05 | 0.56459693 | -0.82471 |
| PABPC1L2B | 5.54E-08 | 2.01E-10 | 0.56455839 | -0.82481 |
| LCE3C | 5.99E-05 | 4.29E-06 | 0.56445274 | -0.82508 |
| TUSC3 | 0.000365 | 4.25E-05 | 0.56435048 | -0.82534 |
| ARHGAP32 | 1.01E-06 | 1.73E-08 | 0.56434547 | -0.82535 |
| PTER | 2.33E-05 | 1.27E-06 | 0.5640402 | -0.82613 |
| TRAM1L1 | 0.000511 | 6.46E-05 | 0.56394149 | -0.82638 |
| PEG3 | 0.00215 | 0.000377 | 0.56381998 | -0.82669 |
| GDAP1L1 | 0.0185 | 0.00525 | 0.56376929 | -0.82682 |
| PEG3-AS1 | 0.00132 | 0.000207 | 0.56360667 | -0.82724 |
| PYGM | 0.0051 | 0.00109 | 0.56347929 | -0.82757 |
| SORBS2 | 0.00312 | 0.000596 | 0.56335538 | -0.82788 |
| NPTX2 | 0.0456 | 0.0161 | 0.56309432 | -0.82855 |
| GRM4 | 3.69E-06 | 1.08E-07 | 0.56239474 | -0.83034 |
| ANKRD24 | 1.71E-09 | 3.33E-13 | 0.56238035 | -0.83038 |
| TOX | 0.000304 | 3.37E-05 | 0.5622642 | -0.83068 |
| AMIGO1 | 1.80E-06 | 3.97E-08 | 0.56189649 | -0.83162 |
| ATPAF1 | 1.24E-07 | 7.51E-10 | 0.56173987 | -0.83203 |
| RSPO3 | 1.61E-05 | 7.85E-07 | 0.56151832 | -0.8326 |
| LRRC2 | 0.0035 | 0.000685 | 0.5614585 | -0.83275 |
| PCDH7 | 0.000197 | 1.95E-05 | 0.56122804 | -0.83334 |
| GPIHBP1 | 4.77E-05 | 3.17E-06 | 0.56110633 | -0.83365 |
| PRKAG2-AS1 | 2.53E-05 | 1.40E-06 | 0.56090751 | -0.83417 |
| LOC440792 | 0.00468 | 0.000975 | 0.56075722 | -0.83455 |
| FBXO44 | 4.65E-07 | 5.50E-09 | 0.56016952 | -0.83606 |
| CORT | 0.000266 | 2.85E-05 | 0.55990156 | -0.83675 |
| NCOA7 | 4.43E-08 | 1.45E-10 | 0.55943794 | -0.83795 |
| VAT1L | 0.0348 | 0.0115 | 0.55942964 | -0.83797 |
| FLJ35700 | 0.00474 | 0.000991 | 0.55915668 | -0.83868 |
| SPRYD3 | 7.31E-08 | 3.28E-10 | 0.55906549 | -0.83891 |

| | | | | |
|---|---|---|---|---|
| YWHAH | 1.42E-07 | 8.92E-10 | 0.55898803 | -0.83911 |
| LINC00643 | 0.0179 | 0.00505 | 0.55884763 | -0.83947 |
| CALM1 | 1.33E-06 | 2.56E-08 | 0.55837223 | -0.8407 |
| RGS20 | 0.00222 | 0.000393 | 0.55833868 | -0.84079 |
| NTSR1 | 0.0139 | 0.00372 | 0.55788675 | -0.84196 |
| PHACTR1 | 4.85E-05 | 3.25E-06 | 0.55766248 | -0.84254 |
| AKR1C1 | 0.00405 | 0.000819 | 0.5576578 | -0.84255 |
| RUNDC3B | 0.000204 | 2.05E-05 | 0.55760577 | -0.84268 |
| TPRG1L | 2.46E-07 | 2.05E-09 | 0.55747175 | -0.84303 |
| GOLT1A | 0.00264 | 0.000485 | 0.55740197 | -0.84321 |
| NTN4 | 0.000179 | 1.73E-05 | 0.55729225 | -0.84349 |
| PDE11A | 0.000221 | 2.26E-05 | 0.55716514 | -0.84382 |
| PLEKHH1 | 0.000287 | 3.13E-05 | 0.55710706 | -0.84397 |
| ARFGEF3 | 0.000144 | 1.31E-05 | 0.5569781 | -0.84431 |
| EPHA7 | 0.00158 | 0.000258 | 0.55661496 | -0.84525 |
| SERP2 | 1.38E-06 | 2.70E-08 | 0.55653036 | -0.84547 |
| PTGDS | 0.00741 | 0.00171 | 0.5564462 | -0.84569 |
| TMEM56 | 0.000629 | 8.29E-05 | 0.55630863 | -0.84604 |
| DNAAF1 | 0.00111 | 0.000167 | 0.55621779 | -0.84628 |
| KNCN | 6.77E-06 | 2.52E-07 | 0.5559506 | -0.84697 |
| MAST1 | 0.000143 | 1.30E-05 | 0.55566174 | -0.84772 |
| C1orf204 | 0.000463 | 5.72E-05 | 0.55507788 | -0.84924 |
| CHST1 | 0.000259 | 2.76E-05 | 0.55486357 | -0.8498 |
| MAPT | 0.000512 | 6.47E-05 | 0.55479831 | -0.84996 |
| CREB3L3 | 9.91E-06 | 4.18E-07 | 0.55468119 | -0.85027 |
| UBXN10 | 4.49E-05 | 2.94E-06 | 0.55424963 | -0.85139 |
| NSF | 2.80E-07 | 2.52E-09 | 0.55413708 | -0.85169 |
| FGF14 | 0.0118 | 0.00303 | 0.55407386 | -0.85185 |
| ASB2 | 3.97E-05 | 2.51E-06 | 0.55391899 | -0.85225 |
| CPAMD8 | 0.000461 | 5.70E-05 | 0.55386363 | -0.8524 |
| AKAP6 | 1.07E-06 | 1.89E-08 | 0.55381126 | -0.85253 |
| PLCL1 | 0.00144 | 0.000229 | 0.55339323 | -0.85362 |
| FCHO1 | 3.10E-06 | 8.50E-08 | 0.55283743 | -0.85507 |
| SH3TC2 | 3.20E-05 | 1.92E-06 | 0.55220647 | -0.85672 |
| B4GALT6 | 0.000916 | 0.000132 | 0.55203177 | -0.85718 |
| NEBL-AS1 | 0.00211 | 0.000368 | 0.55199071 | -0.85728 |
| TTC9 | 1.83E-05 | 9.33E-07 | 0.55173614 | -0.85795 |
| LINC00403 | 0.0347 | 0.0114 | 0.55119932 | -0.85935 |
| KLK5 | 1.87E-06 | 4.24E-08 | 0.55113494 | -0.85952 |
| KCNC3 | 2.53E-06 | 6.38E-08 | 0.55106378 | -0.85971 |
| IL34 | 0.000142 | 1.29E-05 | 0.55093396 | -0.86005 |
| ITSN1 | 1.80E-05 | 9.12E-07 | 0.55090727 | -0.86012 |
| ALDOC | 0.00054 | 6.88E-05 | 0.55080272 | -0.86039 |

| | | | | |
|---|---|---|---|---|
| KANK4 | 0.000211 | 2.13E-05 | 0.55080135 | -0.8604 |
| CPNE7 | 8.39E-07 | 1.37E-08 | 0.550521 | -0.86113 |
| RAB40B | 8.24E-07 | 1.32E-08 | 0.55044305 | -0.86133 |
| NRG4 | 0.000133 | 1.18E-05 | 0.55043004 | -0.86137 |
| KIAA1217 | 7.22E-05 | 5.44E-06 | 0.55012227 | -0.86218 |
| APLP1 | 8.30E-06 | 3.32E-07 | 0.54992421 | -0.8627 |
| GPC5 | 0.0155 | 0.00423 | 0.54941957 | -0.86402 |
| CNTN2 | 0.000784 | 0.000109 | 0.54937817 | -0.86413 |
| NEDD4L | 0.000181 | 1.76E-05 | 0.54894831 | -0.86526 |
| PRKCQ | 0.000184 | 1.80E-05 | 0.54868776 | -0.86594 |
| FBXL2 | 0.00014 | 1.26E-05 | 0.54866848 | -0.86599 |
| RANBP3L | 2.96E-05 | 1.73E-06 | 0.54858094 | -0.86622 |
| LDHD | 3.94E-05 | 2.49E-06 | 0.54823996 | -0.86712 |
| VAMP2 | 2.36E-07 | 1.95E-09 | 0.54779826 | -0.86828 |
| LYPD6B | 4.90E-06 | 1.62E-07 | 0.5475218 | -0.86901 |
| PRKCQ-AS1 | 1.06E-05 | 4.58E-07 | 0.54703665 | -0.87029 |
| OMG | 0.025 | 0.00761 | 0.54697568 | -0.87045 |
| AVPI1 | 4.25E-08 | 1.37E-10 | 0.5468933 | -0.87067 |
| SH3BGRL2 | 1.04E-05 | 4.45E-07 | 0.54686537 | -0.87074 |
| GYPE | 7.63E-08 | 3.59E-10 | 0.54677971 | -0.87097 |
| NBL1 | 1.51E-06 | 3.09E-08 | 0.54652652 | -0.87164 |
| TEF | 4.97E-07 | 6.01E-09 | 0.5463549 | -0.87209 |
| EFNA5 | 0.00299 | 0.000565 | 0.5463335 | -0.87215 |
| PCDHAC2 | 1.23E-05 | 5.56E-07 | 0.54613174 | -0.87268 |
| TMEM191A | 2.02E-06 | 4.69E-08 | 0.54580878 | -0.87353 |
| FAIM2 | 2.87E-07 | 2.60E-09 | 0.54575563 | -0.87367 |
| CAMK4 | 0.000247 | 2.60E-05 | 0.54563152 | -0.874 |
| IQSEC3 | 0.000183 | 1.79E-05 | 0.54552322 | -0.87429 |
| TMCC2 | 6.53E-06 | 2.41E-07 | 0.54543304 | -0.87453 |
| CPNE5 | 0.000129 | 1.13E-05 | 0.54514707 | -0.87528 |
| COL24A1 | 0.000311 | 3.48E-05 | 0.54501589 | -0.87563 |
| CHN2 | 0.00014 | 1.26E-05 | 0.54499685 | -0.87568 |
| RIPPLY2 | 0.0133 | 0.00351 | 0.54482239 | -0.87614 |
| DLG2 | 2.84E-08 | 6.38E-11 | 0.54464285 | -0.87662 |
| ZDHHC8P1 | 0.00704 | 0.00161 | 0.54461144 | -0.8767 |
| GARNL3 | 5.65E-09 | 4.27E-12 | 0.54437216 | -0.87733 |
| SPOCK1 | 0.00325 | 0.000624 | 0.54383654 | -0.87876 |
| STPG1 | 0.00219 | 0.000386 | 0.54381332 | -0.87882 |
| PTPN3 | 0.00162 | 0.000268 | 0.54378833 | -0.87888 |
| TMEM200A | 0.000222 | 2.27E-05 | 0.54364874 | -0.87925 |
| DNAJA4 | 0.000562 | 7.24E-05 | 0.54329271 | -0.8802 |
| HSD11B1 | 0.00871 | 0.00208 | 0.5431479 | -0.88058 |
| SFTPD | 0.000591 | 7.70E-05 | 0.54269093 | -0.8818 |

| | | | | |
|---|---|---|---|---|
| LOC100288911 | 9.58E-06 | 3.99E-07 | 0.54261909 | -0.88199 |
| AMER2 | 0.0156 | 0.00429 | 0.5425705 | -0.88212 |
| NRIP2 | 2.90E-05 | 1.68E-06 | 0.54251112 | -0.88228 |
| GPR68 | 0.00244 | 0.000439 | 0.54171239 | -0.8844 |
| CSRNP3 | 0.000258 | 2.75E-05 | 0.54144492 | -0.88511 |
| LRRC10B | 0.000303 | 3.36E-05 | 0.54142571 | -0.88516 |
| PPP2R2D | 7.28E-07 | 1.08E-08 | 0.54112553 | -0.88596 |
| GPRASP1 | 0.000206 | 2.07E-05 | 0.54074432 | -0.88698 |
| ARL3 | 4.14E-07 | 4.61E-09 | 0.5405088 | -0.88761 |
| RTN2 | 4.52E-05 | 2.97E-06 | 0.54048253 | -0.88768 |
| EPHA10 | 1.68E-06 | 3.62E-08 | 0.5403709 | -0.88798 |
| DOCK3 | 1.39E-05 | 6.48E-07 | 0.5403642 | -0.888 |
| GABARAPL3 | 3.36E-06 | 9.54E-08 | 0.54029738 | -0.88817 |
| KIF17 | 3.29E-08 | 8.28E-11 | 0.54020987 | -0.88841 |
| LRRTM4 | 0.000396 | 4.69E-05 | 0.54010226 | -0.8887 |
| DGKB | 0.023 | 0.00688 | 0.53971123 | -0.88974 |
| TARP | 5.28E-05 | 3.65E-06 | 0.53964823 | -0.88991 |
| SEZ6L2 | 0.00342 | 0.000664 | 0.53953412 | -0.89021 |
| MOAP1 | 5.47E-07 | 6.98E-09 | 0.53945772 | -0.89042 |
| CAMTA1 | 5.34E-06 | 1.81E-07 | 0.53895346 | -0.89177 |
| KRT1 | 0.000992 | 0.000145 | 0.53879184 | -0.8922 |
| RHOV | 3.26E-06 | 9.17E-08 | 0.5387639 | -0.89227 |
| BAIAP3 | 0.0106 | 0.00264 | 0.5385531 | -0.89284 |
| SCN9A | 0.00316 | 0.000604 | 0.53848083 | -0.89303 |
| ATP2C2 | 0.0012 | 0.000183 | 0.53836977 | -0.89333 |
| CLUL1 | 0.00155 | 0.000253 | 0.53800586 | -0.89431 |
| ZFP28 | 1.73E-05 | 8.65E-07 | 0.53780061 | -0.89486 |
| RIMS4 | 0.0124 | 0.00321 | 0.53780043 | -0.89486 |
| SMIM10L2B | 6.28E-05 | 4.56E-06 | 0.53779673 | -0.89487 |
| PPEF1 | 0.000403 | 4.79E-05 | 0.53771533 | -0.89509 |
| CGREF1 | 0.00137 | 0.000216 | 0.5376545 | -0.89525 |
| ANO5 | 0.00549 | 0.00119 | 0.5375862 | -0.89543 |
| CALM3 | 3.73E-08 | 1.10E-10 | 0.53751142 | -0.89563 |
| PGM2L1 | 2.98E-06 | 8.05E-08 | 0.53694953 | -0.89714 |
| MC4R | 0.0228 | 0.0068 | 0.53694369 | -0.89716 |
| GNAS | 2.35E-05 | 1.28E-06 | 0.53663807 | -0.89798 |
| ZNF702P | 2.42E-06 | 6.00E-08 | 0.53660403 | -0.89807 |
| SUGT1P3 | 4.79E-06 | 1.57E-07 | 0.53659503 | -0.89809 |
| DNAH6 | 0.013 | 0.0034 | 0.53657941 | -0.89814 |
| FAM131A | 2.29E-08 | 4.07E-11 | 0.53644152 | -0.89851 |
| ATP1B1 | 0.000168 | 1.60E-05 | 0.53627169 | -0.89896 |
| CAMK2G | 7.05E-06 | 2.66E-07 | 0.53589951 | -0.89997 |
| LOC101928377 | 1.62E-06 | 3.46E-08 | 0.53555688 | -0.90089 |

| | | | | |
|---|---|---|---|---|
| PRRG3 | 2.33E-06 | 5.72E-08 | 0.53542956 | -0.90123 |
| SPRN | 4.27E-07 | 4.86E-09 | 0.53532091 | -0.90152 |
| CELF3 | 0.0153 | 0.00417 | 0.53510985 | -0.90209 |
| SPTBN2 | 4.83E-06 | 1.59E-07 | 0.53489495 | -0.90267 |
| AP3B2 | 7.23E-06 | 2.76E-07 | 0.53486344 | -0.90276 |
| NUAK1 | 7.14E-08 | 3.19E-10 | 0.53478415 | -0.90297 |
| COX7A1 | 0.00119 | 0.000182 | 0.53478192 | -0.90298 |
| LINC00282 | 0.00908 | 0.00219 | 0.53476024 | -0.90304 |
| RAB27B | 0.00152 | 0.000246 | 0.53454555 | -0.90362 |
| CABLES1 | 1.15E-06 | 2.09E-08 | 0.53443952 | -0.9039 |
| SLC6A13 | 0.000724 | 9.87E-05 | 0.53419512 | -0.90456 |
| IQCA1 | 0.02 | 0.00577 | 0.53400294 | -0.90508 |
| PHACTR3 | 0.0216 | 0.00636 | 0.53396649 | -0.90518 |
| RNF43 | 0.0054 | 0.00116 | 0.53377236 | -0.9057 |
| LINC01260 | 7.36E-08 | 3.36E-10 | 0.53326679 | -0.90707 |
| CEP170B | 5.11E-07 | 6.25E-09 | 0.53320998 | -0.90722 |
| DNM3 | 7.00E-05 | 5.22E-06 | 0.53314102 | -0.90741 |
| GPR143 | 0.0039 | 0.000782 | 0.53310802 | -0.9075 |
| WNT2 | 0.0012 | 0.000183 | 0.53277386 | -0.9084 |
| PAK1 | 7.06E-08 | 3.07E-10 | 0.53257023 | -0.90896 |
| CFTR | 6.14E-05 | 4.45E-06 | 0.53238709 | -0.90945 |
| MICU3 | 1.32E-07 | 8.09E-10 | 0.53218885 | -0.90999 |
| RAB26 | 1.08E-06 | 1.92E-08 | 0.53164204 | -0.91147 |
| CCDC110 | 0.000228 | 2.35E-05 | 0.53124509 | -0.91255 |
| ENPP5 | 0.0228 | 0.00678 | 0.53088961 | -0.91352 |
| KIAA1324 | 0.00022 | 2.24E-05 | 0.5303141 | -0.91508 |
| EHD3 | 0.000528 | 6.71E-05 | 0.53024628 | -0.91527 |
| ANKH | 1.07E-05 | 4.64E-07 | 0.5301224 | -0.9156 |
| FAM153A | 1.19E-06 | 2.20E-08 | 0.52990792 | -0.91619 |
| RTP1 | 0.00192 | 0.000328 | 0.52983355 | -0.91639 |
| PSIP1 | 0.000273 | 2.95E-05 | 0.52982815 | -0.9164 |
| SLC6A12 | 2.04E-06 | 4.74E-08 | 0.52980182 | -0.91648 |
| NAP1L3 | 6.34E-06 | 2.31E-07 | 0.52967734 | -0.91681 |
| KCNA2 | 9.84E-05 | 8.09E-06 | 0.52894411 | -0.91881 |
| B3GALT2 | 0.000505 | 6.37E-05 | 0.52884073 | -0.91909 |
| GABRB1 | 0.00235 | 0.000419 | 0.52880433 | -0.91919 |
| LPAR3 | 5.03E-05 | 3.42E-06 | 0.52853661 | -0.91992 |
| PELI3 | 4.05E-06 | 1.24E-07 | 0.52826784 | -0.92066 |
| ZNF540 | 1.81E-06 | 4.00E-08 | 0.52825913 | -0.92068 |
| CLSTN3 | 6.37E-07 | 8.76E-09 | 0.52804468 | -0.92127 |
| ADRB1 | 2.63E-05 | 1.48E-06 | 0.52797298 | -0.92146 |
| CACNG8 | 6.87E-07 | 9.77E-09 | 0.52794586 | -0.92154 |
| GSTM5 | 0.0026 | 0.000475 | 0.52732931 | -0.92322 |

| | | | | |
|---|---|---|---|---|
| PP12613 | 1.21E-06 | 2.26E-08 | 0.5271995 | -0.92358 |
| KIF25 | 0.00677 | 0.00153 | 0.52695603 | -0.92425 |
| SHANK1 | 6.89E-05 | 5.12E-06 | 0.52694402 | -0.92428 |
| MRAP2 | 3.82E-05 | 2.39E-06 | 0.52681207 | -0.92464 |
| EFCAB1 | 0.0376 | 0.0126 | 0.52649601 | -0.92551 |
| GABBR1 | 1.90E-05 | 9.77E-07 | 0.52636377 | -0.92587 |
| SNPH | 7.28E-07 | 1.08E-08 | 0.52602122 | -0.92681 |
| NRXN1 | 0.00182 | 0.000309 | 0.52579401 | -0.92743 |
| CYP26A1 | 0.00404 | 0.000817 | 0.5252258 | -0.92899 |
| LOC440300 | 5.84E-09 | 4.70E-12 | 0.52519228 | -0.92908 |
| OXGR1 | 3.06E-05 | 1.80E-06 | 0.52475801 | -0.93028 |
| STOX1 | 8.00E-05 | 6.19E-06 | 0.52475514 | -0.93028 |
| SFTPC | 0.000168 | 1.60E-05 | 0.52453269 | -0.9309 |
| PCSK6 | 0.00174 | 0.000291 | 0.5243312 | -0.93145 |
| PANX2 | 5.46E-07 | 6.95E-09 | 0.52396171 | -0.93247 |
| NPAS4 | 0.000705 | 9.55E-05 | 0.52363887 | -0.93336 |
| KCNMA1 | 8.53E-06 | 3.44E-07 | 0.52338693 | -0.93405 |
| FOSB | 0.000118 | 1.01E-05 | 0.52321696 | -0.93452 |
| PPP1R3F | 1.98E-07 | 1.48E-09 | 0.52314602 | -0.93471 |
| ATP6V1C1 | 5.93E-07 | 7.91E-09 | 0.52293908 | -0.93529 |
| LPCAT4 | 7.63E-08 | 3.59E-10 | 0.52280299 | -0.93566 |
| PCP4 | 0.0073 | 0.00168 | 0.52267107 | -0.93602 |
| IDS | 2.79E-07 | 2.49E-09 | 0.52227766 | -0.93711 |
| UNC5A | 1.82E-06 | 4.06E-08 | 0.52197022 | -0.93796 |
| TBC1D30 | 7.14E-08 | 3.14E-10 | 0.52186205 | -0.93826 |
| TNIP3 | 0.000936 | 0.000135 | 0.52165092 | -0.93884 |
| GABARAPL1 | 8.16E-07 | 1.30E-08 | 0.52162781 | -0.93891 |
| BAIAP2 | 8.38E-08 | 4.11E-10 | 0.52118628 | -0.94013 |
| OLA1 | 3.31E-06 | 9.35E-08 | 0.52091244 | -0.94089 |
| EPB41L3 | 0.000129 | 1.13E-05 | 0.52057567 | -0.94182 |
| ANKS1B | 0.00345 | 0.000671 | 0.5205069 | -0.94201 |
| EGR3 | 5.68E-05 | 4.01E-06 | 0.52045777 | -0.94215 |
| ACTR3C | 0.00155 | 0.000253 | 0.52045715 | -0.94215 |
| DUSP26 | 0.00205 | 0.000354 | 0.52036802 | -0.9424 |
| FXYD7 | 1.75E-05 | 8.76E-07 | 0.52029249 | -0.94261 |
| LDB3 | 4.79E-06 | 1.57E-07 | 0.52026162 | -0.94269 |
| LGI3 | 3.10E-08 | 7.35E-11 | 0.52021334 | -0.94282 |
| SLC25A27 | 2.10E-05 | 1.11E-06 | 0.52016182 | -0.94297 |
| HENMT1 | 9.14E-07 | 1.54E-08 | 0.51991641 | -0.94365 |
| C9orf129 | 0.00375 | 0.000742 | 0.5198483 | -0.94384 |
| ISLR2 | 0.00025 | 2.63E-05 | 0.51973564 | -0.94415 |
| ZNF215 | 0.000382 | 4.48E-05 | 0.51973524 | -0.94415 |
| SCN1B | 9.65E-06 | 4.03E-07 | 0.51969767 | -0.94426 |

| | | | | |
|---|---|---|---|---|
| CALB2 | 0.00753 | 0.00174 | 0.51893804 | -0.94637 |
| CCNA1 | 0.00357 | 0.000699 | 0.51878109 | -0.9468 |
| DOCK9-AS2 | 0.000302 | 3.35E-05 | 0.51871694 | -0.94698 |
| LOC153811 | 2.62E-05 | 1.47E-06 | 0.51863548 | -0.94721 |
| SLC7A10 | 0.0235 | 0.00704 | 0.5184783 | -0.94764 |
| ENC1 | 2.04E-05 | 1.07E-06 | 0.51814928 | -0.94856 |
| IDI2-AS1 | 0.000963 | 0.00014 | 0.51810611 | -0.94868 |
| CEND1 | 8.59E-06 | 3.49E-07 | 0.51805566 | -0.94882 |
| MEPE | 0.00319 | 0.000612 | 0.51753 | -0.95029 |
| PRDM2 | 6.98E-08 | 2.94E-10 | 0.5175079 | -0.95035 |
| S1PR5 | 8.57E-06 | 3.47E-07 | 0.51670086 | -0.9526 |
| GRIA2 | 0.00563 | 0.00123 | 0.51635168 | -0.95357 |
| CD200 | 1.08E-05 | 4.67E-07 | 0.51632813 | -0.95364 |
| UNC80 | 0.000322 | 3.63E-05 | 0.51538863 | -0.95627 |
| ACTL6B | 9.68E-05 | 7.93E-06 | 0.5153867 | -0.95627 |
| EPHX4 | 1.24E-05 | 5.64E-07 | 0.51528739 | -0.95655 |
| RFPL3S | 5.54E-08 | 2.04E-10 | 0.51504607 | -0.95723 |
| CACNB2 | 2.72E-07 | 2.38E-09 | 0.51500006 | -0.95736 |
| PDXP | 7.12E-07 | 1.04E-08 | 0.51473286 | -0.9581 |
| PALM2 | 0.000598 | 7.81E-05 | 0.51441278 | -0.959 |
| TMEM59L | 0.000139 | 1.25E-05 | 0.51438729 | -0.95907 |
| SSX2IP | 3.48E-07 | 3.56E-09 | 0.51431677 | -0.95927 |
| ELOVL7 | 1.43E-06 | 2.85E-08 | 0.51405709 | -0.96 |
| TSPYL1 | 2.43E-08 | 4.48E-11 | 0.51349075 | -0.96159 |
| TRPM6 | 8.76E-05 | 7.00E-06 | 0.51335456 | -0.96197 |
| BDNF | 0.0143 | 0.00383 | 0.51298936 | -0.963 |
| CACNA2D2 | 9.75E-05 | 8.00E-06 | 0.51297453 | -0.96304 |
| GFOD1 | 5.02E-07 | 6.09E-09 | 0.51296312 | -0.96307 |
| PIP5K1B | 0.000297 | 3.28E-05 | 0.51285411 | -0.96338 |
| ADIRF | 0.00449 | 0.000928 | 0.51283375 | -0.96344 |
| SPTBN4 | 5.57E-06 | 1.93E-07 | 0.51279429 | -0.96355 |
| AARD | 0.000333 | 3.79E-05 | 0.51239322 | -0.96468 |
| MTURN | 0.000138 | 1.23E-05 | 0.51223769 | -0.96511 |
| GUCY1B3 | 1.62E-06 | 3.46E-08 | 0.51219636 | -0.96523 |
| RUNX1T1 | 0.000375 | 4.38E-05 | 0.51214676 | -0.96537 |
| FBXW7 | 5.86E-08 | 2.22E-10 | 0.51178423 | -0.96639 |
| CES5A | 6.30E-06 | 2.28E-07 | 0.51177958 | -0.96641 |
| RAPGEFL1 | 1.63E-07 | 1.15E-09 | 0.51157203 | -0.96699 |
| KLC1 | 6.22E-07 | 8.46E-09 | 0.51127734 | -0.96782 |
| SLC1A2 | 0.00116 | 0.000177 | 0.51108441 | -0.96837 |
| PDIA2 | 0.00035 | 4.04E-05 | 0.51105076 | -0.96846 |
| LINC00599 | 0.00859 | 0.00204 | 0.51094046 | -0.96877 |
| GALNT14 | 0.00012 | 1.03E-05 | 0.5106449 | -0.96961 |

| | | | | |
|---|---|---|---|---|
| NAV3 | 3.49E-05 | 2.13E-06 | 0.51053597 | -0.96992 |
| RIMKLA | 9.48E-06 | 3.93E-07 | 0.5104417 | -0.97018 |
| PLCH2 | 3.62E-05 | 2.24E-06 | 0.51039663 | -0.97031 |
| TP53TG3 | 0.0363 | 0.0121 | 0.51027183 | -0.97066 |
| ANXA3 | 0.00208 | 0.000361 | 0.51002954 | -0.97135 |
| LYNX1 | 1.37E-05 | 6.33E-07 | 0.50983517 | -0.9719 |
| GNAI1 | 0.000757 | 0.000105 | 0.50977626 | -0.97206 |
| SLC26A4 | 0.000117 | 1.00E-05 | 0.50971923 | -0.97223 |
| ST18 | 0.00798 | 0.00187 | 0.50965243 | -0.97241 |
| RBM24 | 0.00126 | 0.000195 | 0.50949793 | -0.97285 |
| SLC17A7 | 2.97E-07 | 2.81E-09 | 0.50936912 | -0.97322 |
| HAPLN2 | 0.00161 | 0.000264 | 0.50886328 | -0.97465 |
| HTR5A | 8.67E-06 | 3.53E-07 | 0.50884494 | -0.9747 |
| EXD1 | 1.85E-07 | 1.36E-09 | 0.50884194 | -0.97471 |
| PRB2 | 4.51E-06 | 1.44E-07 | 0.50869002 | -0.97514 |
| POPDC3 | 0.0027 | 0.000497 | 0.50862621 | -0.97532 |
| DUSP2 | 5.33E-08 | 1.88E-10 | 0.50840436 | -0.97595 |
| TINCR | 1.16E-05 | 5.17E-07 | 0.50832624 | -0.97617 |
| SLIT2 | 0.00275 | 0.000508 | 0.50810463 | -0.9768 |
| CALHM1 | 9.16E-09 | 9.39E-12 | 0.50798239 | -0.97715 |
| NDFIP2 | 1.27E-06 | 2.41E-08 | 0.50777279 | -0.97774 |
| CCBE1 | 2.64E-05 | 1.48E-06 | 0.50719267 | -0.97939 |
| KIAA1211L | 2.56E-06 | 6.51E-08 | 0.50718613 | -0.97941 |
| SYT3 | 7.73E-06 | 3.02E-07 | 0.50713997 | -0.97954 |
| NXPH2 | 0.000325 | 3.67E-05 | 0.50694274 | -0.98011 |
| CHP2 | 2.98E-06 | 8.06E-08 | 0.50682922 | -0.98043 |
| CDKL5 | 3.19E-07 | 3.08E-09 | 0.50636535 | -0.98175 |
| IL1RAPL1 | 0.0105 | 0.00263 | 0.50555361 | -0.98406 |
| CPNE4 | 0.0193 | 0.00553 | 0.50534046 | -0.98467 |
| BFSP1 | 2.04E-05 | 1.07E-06 | 0.50507891 | -0.98542 |
| PLCH1 | 2.21E-05 | 1.17E-06 | 0.50501337 | -0.98561 |
| OR2L13 | 3.09E-05 | 1.83E-06 | 0.50488485 | -0.98597 |
| MYBPC1 | 0.0322 | 0.0104 | 0.50457989 | -0.98685 |
| ITPR1 | 2.73E-08 | 5.99E-11 | 0.50434281 | -0.98752 |
| TUBA8 | 1.08E-06 | 1.92E-08 | 0.50401483 | -0.98846 |
| PDE6H | 8.63E-06 | 3.51E-07 | 0.50391674 | -0.98874 |
| C10orf35 | 0.000383 | 4.50E-05 | 0.50383593 | -0.98897 |
| OR2W3 | 1.56E-07 | 1.05E-09 | 0.50371224 | -0.98933 |
| MGAT4C | 0.0243 | 0.00733 | 0.50361945 | -0.98959 |
| SLC39A12 | 0.00652 | 0.00147 | 0.503563 | -0.98976 |
| UNC5C | 0.00544 | 0.00118 | 0.50303082 | -0.99128 |
| KCNIP4 | 0.00224 | 0.000396 | 0.50301283 | -0.99133 |
| LOC101929384 | 2.70E-05 | 1.53E-06 | 0.50242195 | -0.99303 |

| | | | | |
|---|---|---|---|---|
| ADRA1B | 0.000221 | 2.26E-05 | 0.50233221 | -0.99329 |
| SPINT2 | 1.41E-05 | 6.57E-07 | 0.50193898 | -0.99442 |
| MAP1A | 1.38E-08 | 1.91E-11 | 0.50172899 | -0.99502 |
| NALCN | 7.64E-05 | 5.84E-06 | 0.50146527 | -0.99578 |
| RIIAD1 | 0.000441 | 5.39E-05 | 0.50135402 | -0.9961 |
| TDRD9 | 0.00179 | 0.000301 | 0.5013494 | -0.99611 |
| LOC100129973 | 2.63E-05 | 1.48E-06 | 0.50088264 | -0.99746 |
| CORO6 | 6.91E-07 | 9.88E-09 | 0.50087569 | -0.99748 |
| CLIC5 | 1.15E-05 | 5.08E-07 | 0.5008475 | -0.99756 |
| LOC105376360 | 1.98E-07 | 1.49E-09 | 0.50071699 | -0.99793 |
| DIRAS2 | 3.34E-08 | 9.29E-11 | 0.50058578 | -0.99831 |
| RUFY2 | 7.43E-06 | 2.87E-07 | 0.50054962 | -0.99842 |
| SIDT1 | 0.000848 | 0.00012 | 0.50047378 | -0.99863 |
| MPP7 | 1.82E-06 | 4.03E-08 | 0.50027536 | -0.99921 |
| P2RX5 | 2.89E-06 | 7.67E-08 | 0.50020206 | -0.99942 |
| DNM1P46 | 7.05E-06 | 2.67E-07 | 0.50005182 | -0.99985 |
| ANKRD18A | 0.00102 | 0.00015 | 0.49998964 | -1.00003 |
| PAK3 | 0.000236 | 2.45E-05 | 0.49966719 | -1.00096 |
| SGIP1 | 4.37E-05 | 2.84E-06 | 0.4993778 | -1.0018 |
| ATL1 | 2.72E-07 | 2.38E-09 | 0.49934541 | -1.00189 |
| BICDL2 | 7.20E-07 | 1.06E-08 | 0.49924341 | -1.00218 |
| STXBP5 | 3.10E-07 | 2.95E-09 | 0.49923871 | -1.0022 |
| LOR | 0.000183 | 1.78E-05 | 0.49918058 | -1.00237 |
| CASKIN1 | 7.98E-06 | 3.14E-07 | 0.49905126 | -1.00274 |
| STXBP1 | 2.92E-07 | 2.68E-09 | 0.49837407 | -1.0047 |
| FAM133A | 0.00805 | 0.00189 | 0.49826823 | -1.00501 |
| EPHA5 | 0.000704 | 9.54E-05 | 0.49783239 | -1.00627 |
| ATP4A | 1.21E-05 | 5.45E-07 | 0.49762608 | -1.00687 |
| METTL21C | 0.00216 | 0.000378 | 0.49762439 | -1.00687 |
| CSMD3 | 0.0421 | 0.0145 | 0.49750951 | -1.0072 |
| TSPYL5 | 1.67E-05 | 8.30E-07 | 0.49705686 | -1.00852 |
| FAAH | 6.60E-07 | 9.31E-09 | 0.49705269 | -1.00853 |
| LRRC8B | 1.03E-05 | 4.39E-07 | 0.4968498 | -1.00912 |
| ACSL6 | 2.14E-05 | 1.13E-06 | 0.49673324 | -1.00946 |
| PRSS16 | 0.000216 | 2.20E-05 | 0.49659523 | -1.00986 |
| PFN2 | 6.33E-05 | 4.60E-06 | 0.49585407 | -1.01201 |
| PRRT1 | 1.08E-07 | 6.03E-10 | 0.49583372 | -1.01207 |
| RORB | 0.00589 | 0.00129 | 0.49537453 | -1.01341 |
| LOC285812 | 5.91E-07 | 7.87E-09 | 0.49515266 | -1.01405 |
| JAKMIP3 | 8.55E-05 | 6.78E-06 | 0.49406244 | -1.01723 |
| LOC441666 | 0.000704 | 9.54E-05 | 0.49379673 | -1.01801 |
| CUL3 | 6.41E-08 | 2.49E-10 | 0.49375299 | -1.01814 |
| SCN8A | 1.39E-07 | 8.74E-10 | 0.4934253 | -1.0191 |

| | | | | |
|---|---|---|---|---|
| TUBA4A | 3.71E-07 | 3.92E-09 | 0.49341809 | -1.01912 |
| FBXO41 | 9.95E-07 | 1.71E-08 | 0.49250846 | -1.02178 |
| RNF128 | 0.00295 | 0.000554 | 0.49248063 | -1.02186 |
| GABRG3 | 8.07E-05 | 6.27E-06 | 0.49183991 | -1.02374 |
| PLP1 | 0.0193 | 0.00554 | 0.49175189 | -1.024 |
| SLC5A11 | 0.00025 | 2.64E-05 | 0.49168873 | -1.02418 |
| LOC285696 | 7.22E-06 | 2.75E-07 | 0.49156493 | -1.02455 |
| FGF9 | 0.0135 | 0.00357 | 0.49093764 | -1.02639 |
| LINC00622 | 5.13E-07 | 6.30E-09 | 0.49048391 | -1.02772 |
| NOS1 | 0.00734 | 0.00169 | 0.49003303 | -1.02905 |
| ZNF536 | 0.00126 | 0.000195 | 0.48993491 | -1.02934 |
| PRKCE | 4.51E-07 | 5.22E-09 | 0.48993369 | -1.02934 |
| PPP3CB | 1.49E-08 | 2.15E-11 | 0.48933416 | -1.03111 |
| CLVS1 | 0.000458 | 5.65E-05 | 0.48932992 | -1.03112 |
| SRCIN1 | 6.84E-05 | 5.06E-06 | 0.48893768 | -1.03228 |
| LRRTM1 | 0.000316 | 3.54E-05 | 0.488855 | -1.03252 |
| GRP | 0.000144 | 1.31E-05 | 0.4887333 | -1.03288 |
| SLC26A8 | 8.30E-06 | 3.32E-07 | 0.48864777 | -1.03313 |
| FGF12 | 0.00918 | 0.00222 | 0.48863703 | -1.03316 |
| GJB1 | 7.34E-05 | 5.55E-06 | 0.48860645 | -1.03326 |
| STEAP2 | 0.00125 | 0.000193 | 0.48844983 | -1.03372 |
| LRRC38 | 1.40E-05 | 6.55E-07 | 0.48844604 | -1.03373 |
| AGAP2 | 0.000249 | 2.63E-05 | 0.48827753 | -1.03423 |
| C10orf82 | 4.46E-06 | 1.42E-07 | 0.48799504 | -1.03506 |
| SLC25A41 | 3.14E-05 | 1.88E-06 | 0.48759661 | -1.03624 |
| LRRC73 | 1.16E-05 | 5.17E-07 | 0.48729077 | -1.03715 |
| FAM155A | 0.00945 | 0.0023 | 0.48718655 | -1.03745 |
| GLP2R | 6.67E-06 | 2.47E-07 | 0.48653591 | -1.03938 |
| LINC00982 | 0.000391 | 4.62E-05 | 0.48616681 | -1.04048 |
| BASP1 | 0.000322 | 3.63E-05 | 0.48578054 | -1.04162 |
| SYT5 | 8.37E-07 | 1.36E-08 | 0.48561144 | -1.04213 |
| NELL2 | 0.000304 | 3.38E-05 | 0.48559626 | -1.04217 |
| NOS1AP | 3.09E-05 | 1.83E-06 | 0.48552194 | -1.04239 |
| STK32C | 5.39E-06 | 1.84E-07 | 0.48551629 | -1.04241 |
| CFAP58-AS1 | 5.08E-05 | 3.47E-06 | 0.48545252 | -1.0426 |
| HTR1E | 2.46E-07 | 2.06E-09 | 0.48530267 | -1.04304 |
| CNTN4 | 2.87E-09 | 1.61E-12 | 0.48513989 | -1.04353 |
| CD22 | 3.23E-06 | 9.01E-08 | 0.48501225 | -1.04391 |
| GABRB2 | 3.73E-06 | 1.09E-07 | 0.48460803 | -1.04511 |
| ASPHD1 | 0.000595 | 7.77E-05 | 0.48452157 | -1.04537 |
| CENPVL2 | 0.00465 | 0.000967 | 0.484444 | -1.0456 |
| TENM3 | 0.00477 | 0.000999 | 0.48439867 | -1.04573 |
| TRIM53AP | 0.00935 | 0.00227 | 0.48422397 | -1.04625 |

| | | | | |
|---|---|---|---|---|
| TSPOAP1 | 1.73E-05 | 8.64E-07 | 0.48320082 | -1.04931 |
| KCNN1 | 6.40E-07 | 8.85E-09 | 0.48319549 | -1.04932 |
| LNX1 | 0.00125 | 0.000192 | 0.48274148 | -1.05068 |
| FAXC | 8.53E-06 | 3.44E-07 | 0.48239271 | -1.05172 |
| STXBP5L | 3.06E-06 | 8.37E-08 | 0.4820505 | -1.05274 |
| AKAP5 | 2.27E-07 | 1.86E-09 | 0.48145782 | -1.05452 |
| DNAH17 | 5.22E-06 | 1.76E-07 | 0.48145568 | -1.05453 |
| UPP2 | 1.03E-06 | 1.77E-08 | 0.48132061 | -1.05493 |
| LOC644189 | 7.22E-06 | 2.75E-07 | 0.4812719 | -1.05508 |
| KCTD8 | 0.0035 | 0.000683 | 0.48111704 | -1.05554 |
| FUT9 | 0.0238 | 0.00717 | 0.48092519 | -1.05612 |
| NCS1 | 2.27E-07 | 1.87E-09 | 0.48022912 | -1.05821 |
| DDX25 | 0.00412 | 0.000835 | 0.47960679 | -1.06008 |
| LDLRAD4 | 3.60E-05 | 2.22E-06 | 0.47959858 | -1.0601 |
| CPEB3 | 2.81E-06 | 7.37E-08 | 0.47950325 | -1.06039 |
| FNDC9 | 0.00141 | 0.000223 | 0.47921059 | -1.06127 |
| SATB2-AS1 | 0.000266 | 2.85E-05 | 0.47919103 | -1.06133 |
| FLRT2 | 0.00269 | 0.000496 | 0.47911912 | -1.06154 |
| ATP2B2 | 3.99E-06 | 1.21E-07 | 0.47897774 | -1.06197 |
| ZCCHC12 | 6.38E-06 | 2.33E-07 | 0.47856387 | -1.06322 |
| STXBP6 | 6.43E-05 | 4.69E-06 | 0.47801596 | -1.06487 |
| NPHS1 | 3.57E-05 | 2.20E-06 | 0.47772984 | -1.06573 |
| LINC00672 | 0.000681 | 9.17E-05 | 0.47766623 | -1.06593 |
| KLHL32 | 0.00411 | 0.000833 | 0.47704263 | -1.06781 |
| CA7 | 3.32E-07 | 3.31E-09 | 0.476722 | -1.06878 |
| GLS | 1.50E-06 | 3.06E-08 | 0.47663196 | -1.06905 |
| BSPRY | 1.82E-05 | 9.30E-07 | 0.47606636 | -1.07077 |
| LMO3 | 0.0037 | 0.000731 | 0.47601611 | -1.07092 |
| ADRA2A | 9.79E-06 | 4.12E-07 | 0.47579655 | -1.07158 |
| KCNH5 | 1.14E-07 | 6.56E-10 | 0.47579222 | -1.0716 |
| CBX7 | 2.29E-08 | 4.04E-11 | 0.47555625 | -1.07231 |
| GSTO2 | 3.83E-06 | 1.14E-07 | 0.47538128 | -1.07284 |
| ADRA2C | 0.000102 | 8.45E-06 | 0.47500105 | -1.074 |
| STAMBPL1 | 5.50E-06 | 1.89E-07 | 0.4749805 | -1.07406 |
| DMKN | 0.0011 | 0.000165 | 0.47486943 | -1.0744 |
| TMEM144 | 2.24E-05 | 1.19E-06 | 0.47486894 | -1.0744 |
| SLC13A5 | 0.000406 | 4.85E-05 | 0.47458273 | -1.07527 |
| CRLF1 | 0.0206 | 0.006 | 0.47402367 | -1.07697 |
| MAPK8IP2 | 1.21E-06 | 2.26E-08 | 0.4739529 | -1.07718 |
| ITPKA | 1.21E-07 | 7.25E-10 | 0.47375251 | -1.07779 |
| SEMA4D | 6.24E-07 | 8.51E-09 | 0.47373472 | -1.07785 |
| BHMT | 3.41E-05 | 2.07E-06 | 0.47366862 | -1.07805 |
| TNNI3K | 0.000103 | 8.56E-06 | 0.47354968 | -1.07841 |

| | | | | |
|---|---|---|---|---|
| DCTN1-AS1 | 4.92E-06 | 1.63E-07 | 0.47351611 | -1.07851 |
| FXYD1 | 0.0036 | 0.000707 | 0.4734719 | -1.07865 |
| JPH1 | 3.83E-05 | 2.39E-06 | 0.47319633 | -1.07949 |
| CBFA2T3 | 7.47E-06 | 2.89E-07 | 0.47277554 | -1.08077 |
| RGS14 | 9.62E-07 | 1.64E-08 | 0.47206699 | -1.08294 |
| KLK10 | 0.000251 | 2.65E-05 | 0.47176042 | -1.08387 |
| PNMA5 | 1.05E-06 | 1.83E-08 | 0.47174535 | -1.08392 |
| TUBB4A | 4.83E-06 | 1.59E-07 | 0.47161137 | -1.08433 |
| FABP3 | 0.000158 | 1.48E-05 | 0.4716014 | -1.08436 |
| L1CAM | 0.000928 | 0.000134 | 0.47142151 | -1.08491 |
| CA4 | 0.00155 | 0.000252 | 0.47140315 | -1.08497 |
| DCLK1 | 1.76E-05 | 8.88E-07 | 0.47119231 | -1.08561 |
| GUCA2B | 0.000208 | 2.10E-05 | 0.47103801 | -1.08608 |
| PRKCZ | 4.39E-05 | 2.86E-06 | 0.47089209 | -1.08653 |
| RESP18 | 2.52E-06 | 6.33E-08 | 0.47048724 | -1.08777 |
| NPTXR | 2.87E-06 | 7.58E-08 | 0.46989301 | -1.0896 |
| RGS7BP | 2.23E-06 | 5.34E-08 | 0.46969938 | -1.09019 |
| CDS1 | 3.35E-06 | 9.51E-08 | 0.46942592 | -1.09103 |
| CHRM2 | 0.000719 | 9.79E-05 | 0.4690023 | -1.09233 |
| ANO4 | 4.71E-05 | 3.12E-06 | 0.46859371 | -1.09359 |
| DMRTC1 | 0.00149 | 0.000239 | 0.46842358 | -1.09411 |
| EPCAM | 1.64E-05 | 8.08E-07 | 0.46829246 | -1.09452 |
| FAM19A4 | 3.41E-07 | 3.44E-09 | 0.46788261 | -1.09578 |
| FAM189A1 | 3.84E-05 | 2.40E-06 | 0.46769626 | -1.09636 |
| CAPN13 | 2.55E-08 | 4.97E-11 | 0.46750432 | -1.09695 |
| RAB3A | 8.45E-05 | 6.68E-06 | 0.46745371 | -1.0971 |
| SH2D5 | 4.52E-07 | 5.24E-09 | 0.46690651 | -1.09879 |
| INPP5F | 2.48E-08 | 4.73E-11 | 0.46677312 | -1.09921 |
| FABP1 | 0.000562 | 7.24E-05 | 0.46675167 | -1.09927 |
| GLRA2 | 1.74E-05 | 8.73E-07 | 0.46640304 | -1.10035 |
| ENPP2 | 9.18E-05 | 7.42E-06 | 0.46614542 | -1.10115 |
| TGFBR3L | 1.05E-06 | 1.85E-08 | 0.46610523 | -1.10127 |
| EDIL3 | 0.000219 | 2.24E-05 | 0.46609625 | -1.1013 |
| EPN3 | 0.000851 | 0.000121 | 0.46598351 | -1.10165 |
| CCSER1 | 2.46E-05 | 1.35E-06 | 0.46559162 | -1.10286 |
| CHRM3 | 2.29E-05 | 1.23E-06 | 0.4653749 | -1.10353 |
| GABRG1 | 1.40E-05 | 6.51E-07 | 0.46509228 | -1.10441 |
| ABHD12B | 3.34E-08 | 8.91E-11 | 0.46501588 | -1.10465 |
| ADAP1 | 2.10E-07 | 1.65E-09 | 0.46494572 | -1.10487 |
| CYP4Z1 | 8.57E-06 | 3.47E-07 | 0.46478413 | -1.10537 |
| EMX2OS | 0.00119 | 0.000182 | 0.46450438 | -1.10624 |
| CAMSAP3 | 0.00519 | 0.00111 | 0.46446726 | -1.10635 |
| UNC13A | 2.07E-05 | 1.09E-06 | 0.46442737 | -1.10648 |

| | | | | |
|---|---|---|---|---|
| TAC3 | 0.0108 | 0.0027 | 0.46439071 | -1.10659 |
| NETO1 | 0.000778 | 0.000108 | 0.46438877 | -1.1066 |
| PAIP2B | 2.24E-05 | 1.20E-06 | 0.46432549 | -1.10679 |
| MGAT5B | 5.08E-05 | 3.46E-06 | 0.46431011 | -1.10684 |
| TMEM151A | 1.02E-06 | 1.77E-08 | 0.464137 | -1.10738 |
| ARPP21 | 2.00E-04 | 1.99E-05 | 0.46397874 | -1.10787 |
| GOT1 | 3.29E-08 | 8.20E-11 | 0.46382851 | -1.10834 |
| RPH3A | 0.00136 | 0.000213 | 0.46313985 | -1.11048 |
| KCNS2 | 8.08E-05 | 6.28E-06 | 0.46261245 | -1.11212 |
| PTH2R | 0.000118 | 1.02E-05 | 0.46261181 | -1.11213 |
| DGCR5 | 0.000291 | 3.19E-05 | 0.4626112 | -1.11213 |
| NPFFR2 | 7.08E-06 | 2.68E-07 | 0.46261091 | -1.11213 |
| MAGEE2 | 0.00699 | 0.0016 | 0.46257385 | -1.11224 |
| KIRREL3 | 4.53E-06 | 1.45E-07 | 0.46232853 | -1.11301 |
| RAB6B | 9.62E-08 | 5.07E-10 | 0.46195336 | -1.11418 |
| EPB41L1 | 1.48E-06 | 3.01E-08 | 0.46139766 | -1.11592 |
| DBH | 1.50E-06 | 3.06E-08 | 0.46130729 | -1.1162 |
| HSPA12A | 7.03E-07 | 1.01E-08 | 0.46045013 | -1.11888 |
| KCNB1 | 0.000347 | 3.99E-05 | 0.46024578 | -1.11952 |
| ANK3 | 0.000227 | 2.34E-05 | 0.4602401 | -1.11954 |
| SGPP2 | 2.46E-05 | 1.35E-06 | 0.4601728 | -1.11975 |
| DNAJC6 | 1.17E-06 | 2.16E-08 | 0.46016297 | -1.11978 |
| SMIM6 | 0.000342 | 3.92E-05 | 0.45972378 | -1.12116 |
| NCDN | 8.43E-08 | 4.17E-10 | 0.45929723 | -1.1225 |
| CHGA | 1.20E-06 | 2.23E-08 | 0.45850307 | -1.125 |
| SMIM24 | 2.43E-08 | 4.51E-11 | 0.45814707 | -1.12612 |
| PRKAR1B | 7.92E-07 | 1.24E-08 | 0.45813227 | -1.12616 |
| GABBR2 | 0.000184 | 1.80E-05 | 0.45809077 | -1.12629 |
| ZNF385B | 0.0012 | 0.000184 | 0.45761242 | -1.1278 |
| NRG3 | 1.39E-05 | 6.43E-07 | 0.45726991 | -1.12888 |
| LGI1 | 0.00107 | 0.00016 | 0.45720954 | -1.12907 |
| MYRIP | 2.69E-05 | 1.53E-06 | 0.4570053 | -1.12972 |
| FUT1 | 3.94E-06 | 1.19E-07 | 0.45693983 | -1.12992 |
| SPX | 0.0249 | 0.00755 | 0.45671558 | -1.13063 |
| MACROD2 | 3.80E-06 | 1.12E-07 | 0.45664189 | -1.13086 |
| SCN2A | 5.50E-05 | 3.85E-06 | 0.45631045 | -1.13191 |
| CAMKK1 | 4.57E-07 | 5.35E-09 | 0.45603871 | -1.13277 |
| LINC01140 | 2.26E-05 | 1.21E-06 | 0.45579193 | -1.13355 |
| FRRS1L | 0.000403 | 4.80E-05 | 0.45528909 | -1.13515 |
| ACOT4 | 3.44E-06 | 9.85E-08 | 0.4547922 | -1.13672 |
| RCAN2 | 5.01E-05 | 3.39E-06 | 0.45466539 | -1.13712 |
| KIF12 | 8.38E-06 | 3.37E-07 | 0.45460224 | -1.13732 |
| KIAA0513 | 1.16E-06 | 2.14E-08 | 0.45452042 | -1.13758 |

| | | | | |
|---|---|---|---|---|
| LINC00320 | 0.000739 | 0.000101 | 0.45450126 | -1.13764 |
| PKD2L1 | 7.27E-06 | 2.78E-07 | 0.45409404 | -1.13894 |
| CNTN6 | 2.32E-06 | 5.67E-08 | 0.45377326 | -1.13996 |
| SYNPO2 | 5.33E-05 | 3.69E-06 | 0.4537732 | -1.13996 |
| S100A1 | 0.00347 | 0.000676 | 0.4536119 | -1.14047 |
| FA2H | 0.0162 | 0.00447 | 0.45334667 | -1.14131 |
| HS3ST5 | 9.53E-05 | 7.76E-06 | 0.45321314 | -1.14174 |
| GPR88 | 0.000749 | 0.000103 | 0.45321107 | -1.14175 |
| HAS1 | 6.73E-06 | 2.49E-07 | 0.4531343 | -1.14199 |
| ELFN2 | 0.00231 | 0.000411 | 0.45209583 | -1.1453 |
| ADCYAP1 | 0.00222 | 0.000392 | 0.45200743 | -1.14558 |
| RAB3B | 0.000159 | 1.49E-05 | 0.45175848 | -1.14638 |
| GPR61 | 1.50E-07 | 9.82E-10 | 0.4516463 | -1.14673 |
| KCNQ2 | 7.00E-06 | 2.63E-07 | 0.45162217 | -1.14681 |
| SNRPN | 1.89E-06 | 4.32E-08 | 0.45161681 | -1.14683 |
| SYT9 | 0.00155 | 0.000253 | 0.45149359 | -1.14722 |
| DGKE | 1.09E-05 | 4.72E-07 | 0.45142096 | -1.14745 |
| LDB2 | 2.27E-06 | 5.54E-08 | 0.45140603 | -1.1475 |
| PDZD7 | 0.000109 | 9.20E-06 | 0.45110632 | -1.14846 |
| RAPGEF4 | 2.97E-07 | 2.81E-09 | 0.45039777 | -1.15073 |
| TRIM17 | 0.000108 | 9.13E-06 | 0.44981585 | -1.15259 |
| VSTM2B | 0.000839 | 0.000118 | 0.44977404 | -1.15273 |
| ADTRP | 1.27E-05 | 5.77E-07 | 0.44953533 | -1.15349 |
| PPL | 3.16E-05 | 1.89E-06 | 0.44887825 | -1.1556 |
| RASGEF1A | 1.04E-05 | 4.46E-07 | 0.44864437 | -1.15636 |
| PDE1B | 1.56E-07 | 1.06E-09 | 0.44854773 | -1.15667 |
| CHN1 | 5.94E-08 | 2.26E-10 | 0.44847389 | -1.1569 |
| KCNH3 | 7.76E-11 | 5.67E-15 | 0.44825215 | -1.15762 |
| ERC2 | 8.29E-07 | 1.34E-08 | 0.44820769 | -1.15776 |
| LYZL4 | 0.000679 | 9.13E-05 | 0.44810875 | -1.15808 |
| SORCS1 | 0.00223 | 0.000394 | 0.44803496 | -1.15832 |
| GRM7 | 1.09E-06 | 1.95E-08 | 0.4475721 | -1.15981 |
| BRINP1 | 0.00259 | 0.000473 | 0.44754784 | -1.15989 |
| ABCG4 | 3.55E-06 | 1.02E-07 | 0.447381 | -1.16042 |
| KRT14 | 1.84E-06 | 4.15E-08 | 0.44647463 | -1.16335 |
| CA10 | 0.0204 | 0.00591 | 0.44608913 | -1.1646 |
| CLDN10 | 0.00284 | 0.000529 | 0.44530135 | -1.16715 |
| SYNGR1 | 2.72E-07 | 2.34E-09 | 0.44520506 | -1.16746 |
| ADARB2 | 7.03E-07 | 1.02E-08 | 0.44507353 | -1.16788 |
| PVALB | 2.48E-05 | 1.36E-06 | 0.44504256 | -1.16798 |
| NEUROD2 | 8.29E-07 | 1.34E-08 | 0.44494573 | -1.1683 |
| KIAA1549L | 2.34E-06 | 5.74E-08 | 0.44474053 | -1.16896 |
| MAPK10 | 8.48E-07 | 1.39E-08 | 0.44408659 | -1.17109 |

| | | | | |
|---|---|---|---|---|
| ZNF365 | 3.45E-07 | 3.51E-09 | 0.44398136 | -1.17143 |
| SLC35F3 | 5.37E-05 | 3.73E-06 | 0.44394572 | -1.17154 |
| NEGR1 | 0.00178 | 3.00E-04 | 0.44386194 | -1.17182 |
| SSTR2 | 1.29E-05 | 5.94E-07 | 0.44377761 | -1.17209 |
| SLC7A4 | 3.11E-08 | 7.59E-11 | 0.44348595 | -1.17304 |
| FAM81A | 1.54E-05 | 7.42E-07 | 0.44273818 | -1.17547 |
| ERICH3 | 1.23E-05 | 5.59E-07 | 0.44213731 | -1.17743 |
| CDH22 | 0.000988 | 0.000145 | 0.4420927 | -1.17758 |
| TAGLN3 | 0.000131 | 1.16E-05 | 0.44185438 | -1.17836 |
| TPTE2P1 | 0.000273 | 2.94E-05 | 0.44170948 | -1.17883 |
| ERICH5 | 5.37E-05 | 3.72E-06 | 0.44073195 | -1.18203 |
| TLL2 | 2.55E-08 | 4.96E-11 | 0.44011517 | -1.18405 |
| PTPRN | 1.91E-05 | 9.90E-07 | 0.43999029 | -1.18446 |
| DKFZP586K1520 | 2.86E-06 | 7.56E-08 | 0.43964318 | -1.18559 |
| SV2C | 0.00106 | 0.000157 | 0.43940765 | -1.18637 |
| RALYL | 0.000904 | 0.00013 | 0.43904615 | -1.18756 |
| LOC389332 | 3.91E-06 | 1.17E-07 | 0.43895611 | -1.18785 |
| HSPB3 | 0.00378 | 0.000749 | 0.43856218 | -1.18915 |
| OPCML | 0.00129 | 0.000201 | 0.43840635 | -1.18966 |
| PTPN20 | 0.00409 | 0.000828 | 0.43829788 | -1.19002 |
| PNMA6A | 1.81E-05 | 9.20E-07 | 0.43823606 | -1.19022 |
| OVOL2 | 4.33E-08 | 1.40E-10 | 0.43813598 | -1.19055 |
| SYT7 | 1.29E-05 | 5.93E-07 | 0.43777158 | -1.19175 |
| EPHA8 | 5.34E-06 | 1.82E-07 | 0.43722843 | -1.19354 |
| SMIM10L2A | 2.98E-06 | 8.08E-08 | 0.43711715 | -1.19391 |
| KIF6 | 2.86E-05 | 1.65E-06 | 0.43706189 | -1.19409 |
| TNNT2 | 4.41E-06 | 1.40E-07 | 0.43681412 | -1.19491 |
| INSM2 | 0.00364 | 0.000717 | 0.43673556 | -1.19517 |
| LOC100507140 | 6.25E-05 | 4.53E-06 | 0.43669006 | -1.19532 |
| AFF2 | 0.00127 | 0.000197 | 0.43603499 | -1.19748 |
| LOC100506411 | 0.000117 | 1.00E-05 | 0.43567191 | -1.19869 |
| NPY1R | 0.000664 | 8.88E-05 | 0.43556614 | -1.19904 |
| FBXL16 | 0.00037 | 4.31E-05 | 0.43546096 | -1.19938 |
| EEF1A2 | 0.00233 | 0.000415 | 0.43527275 | -1.20001 |
| ANKRD34C-AS1 | 3.81E-06 | 1.13E-07 | 0.43486527 | -1.20136 |
| TPM3 | 4.25E-06 | 1.33E-07 | 0.43442818 | -1.20281 |
| SLITRK5 | 0.00076 | 0.000105 | 0.43428234 | -1.20329 |
| CHGB | 7.94E-05 | 6.14E-06 | 0.4342799 | -1.2033 |
| PAK6 | 7.25E-06 | 2.77E-07 | 0.43420616 | -1.20355 |
| RASGRF2 | 2.23E-06 | 5.35E-08 | 0.43407517 | -1.20398 |
| GPR32P1 | 7.95E-05 | 6.15E-06 | 0.43369521 | -1.20525 |
| ANKRD19P | 5.40E-07 | 6.83E-09 | 0.43348074 | -1.20596 |
| GDF10 | 0.00054 | 6.89E-05 | 0.43255359 | -1.20905 |

| | | | | |
|---|---|---|---|---|
| HTR1A | 2.87E-09 | 1.41E-12 | 0.43240326 | -1.20955 |
| EXTL1 | 2.88E-07 | 2.63E-09 | 0.43163836 | -1.21211 |
| LOC375196 | 6.26E-09 | 5.20E-12 | 0.43161257 | -1.21219 |
| NRIP3 | 1.84E-06 | 4.14E-08 | 0.43156977 | -1.21233 |
| HECW1 | 3.86E-05 | 2.42E-06 | 0.43146747 | -1.21268 |
| PPP1R16B | 7.09E-08 | 3.10E-10 | 0.43089701 | -1.21459 |
| FOXP2 | 0.00388 | 0.000776 | 0.43081584 | -1.21486 |
| SPTB | 5.56E-06 | 1.92E-07 | 0.43031341 | -1.21654 |
| NTSR2 | 0.00652 | 0.00147 | 0.43025862 | -1.21672 |
| SNAI3-AS1 | 3.46E-06 | 9.95E-08 | 0.43020229 | -1.21691 |
| WBSCR17 | 1.02E-05 | 4.37E-07 | 0.42970432 | -1.21858 |
| CAMK2B | 4.33E-05 | 2.80E-06 | 0.42894991 | -1.22112 |
| SEMA3D | 0.009 | 0.00216 | 0.42784137 | -1.22485 |
| SYT16 | 9.26E-06 | 3.83E-07 | 0.42783188 | -1.22488 |
| ANKRD18B | 0.000168 | 1.59E-05 | 0.42775612 | -1.22514 |
| PDE1A | 6.92E-06 | 2.59E-07 | 0.42746242 | -1.22613 |
| PCDH20 | 0.00348 | 0.000678 | 0.42611082 | -1.2307 |
| GRIN2B | 2.59E-06 | 6.61E-08 | 0.42588189 | -1.23147 |
| PNMAL2 | 2.70E-08 | 5.87E-11 | 0.42573875 | -1.23196 |
| PNMT | 0.00263 | 0.000483 | 0.42569012 | -1.23212 |
| ZMAT4 | 0.000996 | 0.000146 | 0.42555671 | -1.23258 |
| KCNJ6 | 0.000191 | 1.87E-05 | 0.425481 | -1.23283 |
| NTM-AS1 | 0.000584 | 7.59E-05 | 0.42546068 | -1.2329 |
| DYDC2 | 0.00112 | 0.000168 | 0.42518299 | -1.23384 |
| DPYS | 4.02E-05 | 2.55E-06 | 0.42489633 | -1.23482 |
| MYH7B | 4.00E-07 | 4.39E-09 | 0.42479683 | -1.23516 |
| C9orf24 | 2.71E-05 | 1.55E-06 | 0.424735 | -1.23537 |
| SLCO1A2 | 2.73E-05 | 1.56E-06 | 0.4243195 | -1.23678 |
| PEX5L | 8.29E-07 | 1.34E-08 | 0.42410303 | -1.23751 |
| PARM1 | 2.07E-05 | 1.09E-06 | 0.42329821 | -1.24025 |
| NGB | 2.72E-07 | 2.37E-09 | 0.42215623 | -1.24415 |
| HLF | 2.57E-05 | 1.43E-06 | 0.42153116 | -1.24629 |
| ERMN | 0.000292 | 3.20E-05 | 0.42137768 | -1.24681 |
| XK | 0.00032 | 3.59E-05 | 0.42096553 | -1.24823 |
| DAW1 | 4.76E-06 | 1.55E-07 | 0.42093002 | -1.24835 |
| SLC27A2 | 1.43E-06 | 2.87E-08 | 0.42039727 | -1.25017 |
| PCDH11Y | 0.00324 | 0.000622 | 0.42020438 | -1.25084 |
| TRABD2A | 4.41E-06 | 1.40E-07 | 0.41995071 | -1.25171 |
| MAST3 | 5.22E-09 | 3.69E-12 | 0.41991007 | -1.25185 |
| PNMA3 | 2.88E-06 | 7.65E-08 | 0.41933051 | -1.25384 |
| KLK6 | 0.00164 | 0.00027 | 0.41910214 | -1.25463 |
| PRKCG | 2.28E-09 | 6.93E-13 | 0.41895439 | -1.25513 |
| NNAT | 0.0478 | 0.0171 | 0.41878475 | -1.25572 |

| | | | | |
|---|---|---|---|---|
| AMPH | 1.42E-05 | 6.67E-07 | 0.41870565 | -1.25599 |
| CBLN1 | 0.000531 | 6.75E-05 | 0.41857011 | -1.25646 |
| ABLIM2 | 3.43E-06 | 9.80E-08 | 0.41800477 | -1.25841 |
| SPHKAP | 0.000938 | 0.000136 | 0.41796395 | -1.25855 |
| B3GNT4 | 1.78E-05 | 8.98E-07 | 0.41766524 | -1.25958 |
| OGDHL | 0.000314 | 3.50E-05 | 0.417576 | -1.25989 |
| PRSS2 | 1.05E-06 | 1.83E-08 | 0.41753736 | -1.26002 |
| DOC2A | 5.97E-07 | 8.02E-09 | 0.41740969 | -1.26046 |
| FREM3 | 0.00299 | 0.000564 | 0.4173936 | -1.26052 |
| CBLN4 | 0.000245 | 2.57E-05 | 0.4173459 | -1.26068 |
| ATOH7 | 2.50E-06 | 6.26E-08 | 0.41687709 | -1.26231 |
| FBXO2 | 8.09E-05 | 6.29E-06 | 0.41670427 | -1.2629 |
| LOC100507387 | 8.03E-05 | 6.22E-06 | 0.41638569 | -1.26401 |
| CYP46A1 | 8.32E-07 | 1.35E-08 | 0.41596882 | -1.26545 |
| FRAS1 | 9.11E-05 | 7.36E-06 | 0.41546922 | -1.26719 |
| ASIC2 | 2.13E-07 | 1.70E-09 | 0.41531132 | -1.26773 |
| TRIM58 | 1.50E-07 | 9.94E-10 | 0.41493101 | -1.26906 |
| STMN2 | 0.0079 | 0.00185 | 0.41490346 | -1.26915 |
| FABP6 | 0.000859 | 0.000122 | 0.41478002 | -1.26958 |
| KCNJ12 | 2.81E-06 | 7.39E-08 | 0.41476501 | -1.26963 |
| GALNT9 | 0.000189 | 1.85E-05 | 0.41461051 | -1.27017 |
| CLCA4 | 3.90E-06 | 1.17E-07 | 0.41449578 | -1.27057 |
| 5-Sep | 2.87E-09 | 1.61E-12 | 0.41433302 | -1.27114 |
| BSN | 1.65E-05 | 8.11E-07 | 0.41419312 | -1.27162 |
| OLFM1 | 0.000119 | 1.03E-05 | 0.41390119 | -1.27264 |
| GRIN1 | 2.97E-08 | 6.89E-11 | 0.41291181 | -1.27609 |
| NMNAT2 | 4.37E-05 | 2.84E-06 | 0.41286651 | -1.27625 |
| TMEM266 | 2.46E-05 | 1.35E-06 | 0.41255618 | -1.27734 |
| ADAM11 | 7.92E-08 | 3.83E-10 | 0.41240525 | -1.27787 |
| CTXN3 | 0.00236 | 0.000422 | 0.41238813 | -1.27793 |
| CNTNAP2 | 4.03E-06 | 1.22E-07 | 0.41233356 | -1.27812 |
| DLEU7 | 0.000302 | 3.34E-05 | 0.41173908 | -1.2802 |
| FGF13 | 0.00071 | 9.64E-05 | 0.41168829 | -1.28038 |
| SLITRK1 | 0.00337 | 0.000654 | 0.41164962 | -1.28051 |
| RASAL1 | 3.09E-06 | 8.48E-08 | 0.41136011 | -1.28153 |
| GRIP1 | 3.76E-06 | 1.11E-07 | 0.41084827 | -1.28332 |
| ANKRD34A | 9.62E-06 | 4.01E-07 | 0.41058215 | -1.28426 |
| KCNK12 | 0.000219 | 2.23E-05 | 0.41011326 | -1.28591 |
| DRD5 | 8.14E-07 | 1.29E-08 | 0.40997112 | -1.28641 |
| ESYT3 | 2.61E-08 | 5.35E-11 | 0.40885486 | -1.29034 |
| PAK5 | 0.00404 | 0.000816 | 0.40872268 | -1.29081 |
| ATP6V1G2 | 7.96E-06 | 3.13E-07 | 0.40871353 | -1.29084 |
| UBE2QL1 | 0.000115 | 9.81E-06 | 0.4075547 | -1.29493 |

| | | | | |
|---|---|---|---|---|
| RIMS1 | 5.42E-06 | 1.85E-07 | 0.40733181 | -1.29572 |
| RFPL3 | 1.82E-06 | 4.03E-08 | 0.40713219 | -1.29643 |
| FSTL4 | 1.23E-05 | 5.56E-07 | 0.40678751 | -1.29765 |
| KCNG3 | 2.79E-06 | 7.31E-08 | 0.40672391 | -1.29788 |
| CNTNAP4 | 1.38E-06 | 2.69E-08 | 0.40636084 | -1.29917 |
| CYP26B1 | 2.64E-06 | 6.79E-08 | 0.40613802 | -1.29996 |
| MAG | 0.00165 | 0.000272 | 0.40609647 | -1.30011 |
| RAB11FIP4 | 9.48E-07 | 1.61E-08 | 0.4058765 | -1.30089 |
| PPFIA2 | 0.000885 | 0.000126 | 0.40586744 | -1.30092 |
| CNTN5 | 0.000133 | 1.17E-05 | 0.4057052 | -1.3015 |
| PPP1R1B | 0.00185 | 0.000313 | 0.40514836 | -1.30348 |
| NAPB | 2.56E-07 | 2.14E-09 | 0.40500783 | -1.30398 |
| BEX5 | 5.16E-05 | 3.54E-06 | 0.40494009 | -1.30422 |
| NRXN3 | 2.89E-05 | 1.67E-06 | 0.40387625 | -1.30801 |
| BCL11A | 8.99E-07 | 1.51E-08 | 0.40320793 | -1.3104 |
| NEFH | 9.77E-05 | 8.03E-06 | 0.40296806 | -1.31126 |
| ELAVL2 | 4.59E-06 | 1.47E-07 | 0.40266355 | -1.31235 |
| TMEM246 | 0.000358 | 4.15E-05 | 0.40258884 | -1.31262 |
| CACNA2D3 | 5.63E-05 | 3.98E-06 | 0.40251236 | -1.31289 |
| SGSM1 | 4.75E-05 | 3.15E-06 | 0.40214456 | -1.31421 |
| DNM1 | 2.92E-07 | 2.69E-09 | 0.40199284 | -1.31476 |
| HTR4 | 0.000817 | 0.000115 | 0.40175927 | -1.3156 |
| C3orf80 | 2.30E-05 | 1.23E-06 | 0.40165516 | -1.31597 |
| GPR22 | 5.91E-06 | 2.09E-07 | 0.40158425 | -1.31623 |
| HHATL | 0.000657 | 8.76E-05 | 0.40157061 | -1.31627 |
| GABRB3 | 0.000788 | 0.00011 | 0.40147738 | -1.31661 |
| KIAA0319 | 9.22E-05 | 7.46E-06 | 0.40142134 | -1.31681 |
| RGS7 | 3.32E-06 | 9.40E-08 | 0.4008522 | -1.31886 |
| CACNG2 | 7.73E-06 | 3.02E-07 | 0.4008307 | -1.31894 |
| NRSN1 | 0.000386 | 4.54E-05 | 0.40069719 | -1.31942 |
| MYADML2 | 5.18E-07 | 6.42E-09 | 0.40047103 | -1.32023 |
| MIR124-2HG | 0.00453 | 0.000938 | 0.40010676 | -1.32154 |
| SHANK2 | 0.000134 | 1.19E-05 | 0.40006785 | -1.32168 |
| STAT4 | 3.73E-08 | 1.10E-10 | 0.39934197 | -1.3243 |
| RPRML | 0.000445 | 5.46E-05 | 0.39922902 | -1.32471 |
| MOG | 0.00621 | 0.00138 | 0.39876929 | -1.32637 |
| KRT17 | 4.46E-06 | 1.42E-07 | 0.39864689 | -1.32682 |
| STAR | 7.55E-06 | 2.93E-07 | 0.39844514 | -1.32755 |
| GPR62 | 9.57E-06 | 3.98E-07 | 0.39809661 | -1.32881 |
| SLC24A2 | 1.54E-05 | 7.40E-07 | 0.39735461 | -1.3315 |
| HS3ST2 | 0.00757 | 0.00176 | 0.3968648 | -1.33328 |
| TPPP | 1.67E-06 | 3.60E-08 | 0.39648599 | -1.33466 |
| PPP1R1A | 7.36E-05 | 5.58E-06 | 0.39626175 | -1.33547 |

| | | | | |
|---|---|---|---|---|
| LINC01361 | 6.07E-05 | 4.37E-06 | 0.3960351 | -1.3363 |
| RTN4R | 2.10E-07 | 1.64E-09 | 0.3956838 | -1.33758 |
| TCERG1L | 7.23E-05 | 5.45E-06 | 0.39551006 | -1.33821 |
| CALB1 | 0.000153 | 1.42E-05 | 0.39510807 | -1.33968 |
| TENM2 | 0.00122 | 0.000187 | 0.39506017 | -1.33986 |
| AIFM3 | 0.001 | 0.000147 | 0.3947661 | -1.34093 |
| ENTPD3 | 1.02E-07 | 5.54E-10 | 0.39411463 | -1.34331 |
| PCLO | 1.31E-06 | 2.52E-08 | 0.39378856 | -1.34451 |
| MKX | 1.75E-05 | 8.77E-07 | 0.39321635 | -1.3466 |
| ARHGEF7 | 0.000576 | 7.47E-05 | 0.39315652 | -1.34682 |
| CRTAC1 | 0.000971 | 0.000142 | 0.39275926 | -1.34828 |
| RTN1 | 4.42E-06 | 1.40E-07 | 0.39237679 | -1.34969 |
| MAL | 6.06E-06 | 2.17E-07 | 0.39201028 | -1.35104 |
| SYNGR3 | 2.08E-06 | 4.89E-08 | 0.39199575 | -1.35109 |
| SYNPO | 1.99E-06 | 4.58E-08 | 0.39144833 | -1.35311 |
| KIAA1107 | 7.73E-06 | 3.02E-07 | 0.39143091 | -1.35317 |
| NAP1L2 | 3.66E-05 | 2.27E-06 | 0.39089171 | -1.35516 |
| KCNJ9 | 7.46E-06 | 2.88E-07 | 0.39072612 | -1.35577 |
| PRPH2 | 1.56E-07 | 1.07E-09 | 0.39055009 | -1.35642 |
| KCTD16 | 1.33E-06 | 2.55E-08 | 0.39030485 | -1.35733 |
| MIR7-3HG | 0.000112 | 9.51E-06 | 0.38961043 | -1.3599 |
| PRSS3 | 1.89E-06 | 4.34E-08 | 0.38959026 | -1.35997 |
| SLC22A9 | 2.87E-08 | 6.50E-11 | 0.38932951 | -1.36094 |
| SNCG | 2.28E-09 | 8.54E-13 | 0.38930757 | -1.36102 |
| CES4A | 1.57E-07 | 1.08E-09 | 0.38843758 | -1.36425 |
| TNNT1 | 0.000208 | 2.10E-05 | 0.38781486 | -1.36656 |
| RIMS3 | 1.24E-06 | 2.33E-08 | 0.3875033 | -1.36772 |
| KNDC1 | 5.48E-05 | 3.82E-06 | 0.3871663 | -1.36897 |
| SPOCK3 | 0.000153 | 1.41E-05 | 0.38681491 | -1.37028 |
| FAM153B | 0.000123 | 1.07E-05 | 0.38680896 | -1.37031 |
| PNCK | 7.20E-07 | 1.06E-08 | 0.38611246 | -1.37291 |
| JAKMIP1 | 7.92E-05 | 6.12E-06 | 0.38604759 | -1.37315 |
| KCNAB1 | 4.15E-06 | 1.28E-07 | 0.38598664 | -1.37338 |
| C1QL2 | 9.64E-05 | 7.88E-06 | 0.38540027 | -1.37557 |
| IL12RB2 | 9.27E-06 | 3.84E-07 | 0.38519498 | -1.37634 |
| NECAB2 | 7.05E-05 | 5.28E-06 | 0.38508905 | -1.37674 |
| CDKL2 | 0.000376 | 4.40E-05 | 0.38490134 | -1.37744 |
| PCP4L1 | 0.00157 | 0.000256 | 0.38470828 | -1.37816 |
| MUM1L1 | 0.000446 | 5.48E-05 | 0.3842152 | -1.38001 |
| RELN | 2.33E-05 | 1.26E-06 | 0.38363015 | -1.38221 |
| CAMK1G | 6.77E-07 | 9.61E-09 | 0.38355833 | -1.38248 |
| CPLX1 | 0.000168 | 1.60E-05 | 0.38348759 | -1.38275 |
| ACVR1C | 0.000185 | 1.82E-05 | 0.38331565 | -1.3834 |

| | | | | |
|---|---|---|---|---|
| MATK | 8.81E-07 | 1.47E-08 | 0.38299891 | -1.38459 |
| EPB41L4B | 0.000174 | 1.67E-05 | 0.38196777 | -1.38848 |
| SDR16C5 | 1.68E-07 | 1.20E-09 | 0.3817598 | -1.38926 |
| MCF2 | 0.00123 | 0.000188 | 0.38154163 | -1.39009 |
| DACH2 | 0.000956 | 0.000139 | 0.38087677 | -1.3926 |
| SNCB | 5.18E-07 | 6.39E-09 | 0.38056134 | -1.3938 |
| CDH12 | 4.26E-05 | 2.75E-06 | 0.37997663 | -1.39602 |
| PTPN5 | 0.000183 | 1.78E-05 | 0.37971358 | -1.39702 |
| NDST3 | 0.000167 | 1.58E-05 | 0.37959861 | -1.39745 |
| LOC101060524///DRD5P2 | 7.23E-05 | 5.45E-06 | 0.37928686 | -1.39864 |
| CA11 | 3.58E-07 | 3.72E-09 | 0.37868529 | -1.40093 |
| KCNS1 | 2.74E-05 | 1.56E-06 | 0.37850254 | -1.40163 |
| AJAP1 | 1.62E-05 | 7.94E-07 | 0.37843447 | -1.40188 |
| LY6H | 0.00031 | 3.45E-05 | 0.37841747 | -1.40195 |
| GAST | 0.000139 | 1.25E-05 | 0.37829982 | -1.4024 |
| SLC8A2 | 5.91E-06 | 2.08E-07 | 0.37803822 | -1.4034 |
| PLCXD3 | 0.000292 | 3.21E-05 | 0.37801912 | -1.40347 |
| KIF5A | 1.38E-07 | 8.60E-10 | 0.37757857 | -1.40515 |
| VSTM2L | 0.000205 | 2.06E-05 | 0.37754863 | -1.40527 |
| CALN1 | 4.12E-05 | 2.64E-06 | 0.37722482 | -1.4065 |
| FAM163B | 2.68E-05 | 1.52E-06 | 0.37690358 | -1.40773 |
| RFPL1 | 2.70E-08 | 5.75E-11 | 0.37655688 | -1.40906 |
| PSD | 3.07E-06 | 8.39E-08 | 0.37652198 | -1.40919 |
| CRHBP | 0.000232 | 2.40E-05 | 0.37567856 | -1.41243 |
| SH3GL3 | 1.84E-06 | 4.15E-08 | 0.37564385 | -1.41256 |
| PDYN | 0.0173 | 0.00484 | 0.37526959 | -1.414 |
| SCN3B | 6.31E-07 | 8.66E-09 | 0.37483675 | -1.41567 |
| CYP4X1 | 2.36E-06 | 5.79E-08 | 0.37435032 | -1.41754 |
| TAC1 | 0.014 | 0.00373 | 0.37428014 | -1.41781 |
| NGEF | 0.000116 | 9.95E-06 | 0.37375247 | -1.41985 |
| HMP19 | 0.00253 | 0.00046 | 0.37325397 | -1.42177 |
| ATRNL1 | 1.43E-06 | 2.88E-08 | 0.37314542 | -1.42219 |
| CPNE9 | 2.40E-05 | 1.31E-06 | 0.37260747 | -1.42427 |
| RIMS2 | 0.000265 | 2.84E-05 | 0.3725224 | -1.4246 |
| CACNG3 | 1.05E-07 | 5.73E-10 | 0.37226467 | -1.4256 |
| MTUS2 | 1.74E-05 | 8.67E-07 | 0.37189617 | -1.42703 |
| RUNDC3A | 1.16E-05 | 5.11E-07 | 0.37182663 | -1.4273 |
| KCNA1 | 2.33E-05 | 1.27E-06 | 0.37132197 | -1.42926 |
| LHX6 | 2.24E-05 | 1.20E-06 | 0.37130465 | -1.42932 |
| PRKCB | 3.01E-06 | 8.19E-08 | 0.37022723 | -1.43352 |
| SLC1A6 | 0.000426 | 5.16E-05 | 0.37010161 | -1.43401 |
| GRIN2A | 6.01E-07 | 8.12E-09 | 0.37001973 | -1.43433 |
| GABRD | 0.000194 | 1.91E-05 | 0.36995705 | -1.43457 |

| | | | | |
|---|---|---|---|---|
| SAMD12 | 1.09E-06 | 1.95E-08 | 0.36977969 | -1.43526 |
| PPP2R2C | 1.82E-06 | 4.03E-08 | 0.36968869 | -1.43562 |
| RAP1GAP2 | 5.14E-06 | 1.72E-07 | 0.36912803 | -1.43781 |
| KCNK1 | 0.00229 | 0.000406 | 0.36899398 | -1.43833 |
| SNCA | 2.40E-06 | 5.92E-08 | 0.36882969 | -1.43897 |
| INA | 3.34E-05 | 2.02E-06 | 0.36844945 | -1.44046 |
| REPS2 | 3.44E-06 | 9.84E-08 | 0.36800784 | -1.44219 |
| CARNS1 | 0.000266 | 2.85E-05 | 0.3675996 | -1.44379 |
| DOK6 | 5.92E-05 | 4.24E-06 | 0.36739299 | -1.4446 |
| LINC00889 | 1.49E-08 | 2.26E-11 | 0.36683431 | -1.4468 |
| UMODL1-AS1 | 1.22E-06 | 2.28E-08 | 0.36671866 | -1.44725 |
| CIDEA | 2.17E-06 | 5.14E-08 | 0.36634443 | -1.44873 |
| CELF5 | 3.96E-05 | 2.50E-06 | 0.36496348 | -1.45418 |
| PLPPR3 | 0.000218 | 2.22E-05 | 0.36494152 | -1.45426 |
| WNK2 | 4.86E-06 | 1.60E-07 | 0.36416357 | -1.45734 |
| ATP8A2 | 0.000742 | 0.000102 | 0.36402439 | -1.45789 |
| CLEC4G | 2.89E-05 | 1.67E-06 | 0.36381751 | -1.45871 |
| CELF4 | 1.72E-06 | 3.76E-08 | 0.36284817 | -1.46256 |
| PDE2A | 2.33E-05 | 1.26E-06 | 0.3625966 | -1.46356 |
| DPP10 | 0.00048 | 5.98E-05 | 0.36232358 | -1.46465 |
| KCNC1 | 3.74E-07 | 3.97E-09 | 0.36227225 | -1.46485 |
| SCN2B | 6.17E-07 | 8.36E-09 | 0.36213467 | -1.4654 |
| SRRM4 | 0.000391 | 4.62E-05 | 0.36030463 | -1.47271 |
| MAP7D2 | 4.77E-07 | 5.68E-09 | 0.36025042 | -1.47293 |
| PRMT8 | 1.82E-05 | 9.30E-07 | 0.36018575 | -1.47319 |
| ST8SIA3 | 3.62E-05 | 2.24E-06 | 0.35997005 | -1.47405 |
| LRFN2 | 0.000226 | 2.32E-05 | 0.35989434 | -1.47435 |
| NELL1 | 8.20E-05 | 6.43E-06 | 0.35971482 | -1.47507 |
| JPH3 | 1.50E-06 | 3.06E-08 | 0.35925763 | -1.47691 |
| RSPO2 | 9.46E-05 | 7.69E-06 | 0.35850966 | -1.47992 |
| SLC7A14 | 0.000739 | 0.000101 | 0.35829378 | -1.48079 |
| LOC441052 | 3.19E-07 | 3.05E-09 | 0.35818924 | -1.48121 |
| SLC4A10 | 1.54E-06 | 3.20E-08 | 0.35804333 | -1.48179 |
| LRRC7 | 1.48E-05 | 7.03E-07 | 0.35773972 | -1.48302 |
| CLEC2L | 4.07E-05 | 2.59E-06 | 0.35716829 | -1.48532 |
| ST6GALNAC5 | 8.37E-06 | 3.36E-07 | 0.35676038 | -1.48697 |
| AMER3 | 0.00122 | 0.000188 | 0.35615198 | -1.48944 |
| CNNM1 | 1.95E-07 | 1.45E-09 | 0.35545985 | -1.49224 |
| TMEM125 | 3.34E-05 | 2.02E-06 | 0.3549744 | -1.49421 |
| SYCE1 | 1.19E-06 | 2.21E-08 | 0.35475313 | -1.49511 |
| SLC17A6 | 0.00127 | 0.000198 | 0.35461132 | -1.49569 |
| SEC14L5 | 6.13E-06 | 2.20E-07 | 0.35425109 | -1.49716 |
| CDH8 | 2.03E-06 | 4.72E-08 | 0.35383368 | -1.49886 |

| | | | | |
|---|---|---|---|---|
| CNTN3 | 0.000412 | 4.94E-05 | 0.35377981 | -1.49908 |
| DMTN | 7.18E-07 | 1.05E-08 | 0.35357123 | -1.49993 |
| C1QTNF4 | 2.56E-06 | 6.51E-08 | 0.35346368 | -1.50037 |
| VIPR1 | 3.05E-06 | 8.34E-08 | 0.35321793 | -1.50137 |
| C1orf115 | 6.62E-08 | 2.66E-10 | 0.3528351 | -1.50293 |
| CPNE6 | 1.18E-05 | 5.28E-07 | 0.35281419 | -1.50302 |
| SYP | 3.70E-07 | 3.90E-09 | 0.35229502 | -1.50514 |
| GABRG2 | 2.18E-05 | 1.16E-06 | 0.35223613 | -1.50539 |
| DYNC1I1 | 0.000336 | 3.83E-05 | 0.351582 | -1.50807 |
| IPCEF1 | 9.62E-07 | 1.65E-08 | 0.35155841 | -1.50816 |
| HRH3 | 7.31E-06 | 2.80E-07 | 0.35100213 | -1.51045 |
| WIF1 | 4.03E-07 | 4.45E-09 | 0.35051529 | -1.51245 |
| CNKSR2 | 1.06E-07 | 5.86E-10 | 0.34975086 | -1.5156 |
| EPHB6 | 1.88E-06 | 4.26E-08 | 0.34934636 | -1.51727 |
| CNDP1 | 0.00763 | 0.00177 | 0.34923963 | -1.51771 |
| SOWAHA | 3.56E-06 | 1.03E-07 | 0.34800535 | -1.52282 |
| GRM3 | 1.29E-05 | 5.93E-07 | 0.34754507 | -1.52473 |
| SNAP91 | 3.10E-06 | 8.52E-08 | 0.34726683 | -1.52588 |
| LYPD8 | 2.97E-07 | 2.79E-09 | 0.34628982 | -1.52995 |
| LOC440040 | 0.000934 | 0.000135 | 0.34563334 | -1.53269 |
| MCHR2 | 7.52E-08 | 3.50E-10 | 0.34560653 | -1.5328 |
| FBXO40 | 1.34E-08 | 1.75E-11 | 0.34534387 | -1.53389 |
| HPCAL4 | 0.000172 | 1.65E-05 | 0.34469221 | -1.53662 |
| LRFN5 | 9.12E-06 | 3.76E-07 | 0.34455221 | -1.53721 |
| STX1A | 2.14E-07 | 1.71E-09 | 0.34395351 | -1.53971 |
| GPR83 | 8.41E-06 | 3.39E-07 | 0.34364117 | -1.54103 |
| CRH | 0.00793 | 0.00186 | 0.34347633 | -1.54172 |
| CCDC85A | 1.20E-05 | 5.40E-07 | 0.3416126 | -1.54957 |
| FLJ33534 | 7.89E-09 | 6.93E-12 | 0.34149174 | -1.55008 |
| PTPRT | 2.89E-05 | 1.67E-06 | 0.33997249 | -1.55651 |
| RGS4 | 1.15E-05 | 5.08E-07 | 0.33994324 | -1.55663 |
| RASGRF1 | 3.90E-06 | 1.17E-07 | 0.33931845 | -1.55929 |
| KHDRBS2 | 7.35E-05 | 5.56E-06 | 0.33929395 | -1.55939 |
| KCNB2 | 2.81E-06 | 7.37E-08 | 0.33860397 | -1.56233 |
| WNT10B | 7.34E-06 | 2.82E-07 | 0.33803499 | -1.56476 |
| NKX6-2 | 0.0122 | 0.00314 | 0.33798767 | -1.56496 |
| LOC100996385 | 2.72E-06 | 7.05E-08 | 0.33793369 | -1.56519 |
| ARHGAP44 | 1.08E-05 | 4.67E-07 | 0.3379195 | -1.56525 |
| NEURL1 | 9.27E-06 | 3.84E-07 | 0.33752281 | -1.56694 |
| KCNC2 | 4.79E-05 | 3.19E-06 | 0.33714603 | -1.56855 |
| NECAB1 | 2.08E-07 | 1.61E-09 | 0.3358326 | -1.57419 |
| C11orf97 | 3.77E-05 | 2.35E-06 | 0.33548452 | -1.57568 |
| SOSTDC1 | 5.64E-08 | 2.09E-10 | 0.3353881 | -1.5761 |

| | | | | |
|---|---|---|---|---|
| RIMBP2 | 0.00023 | 2.38E-05 | 0.33534486 | -1.57628 |
| CKMT1A | 2.91E-05 | 1.69E-06 | 0.33517787 | -1.577 |
| HMGCLL1 | 1.59E-05 | 7.77E-07 | 0.33501556 | -1.5777 |
| SLC32A1 | 0.00383 | 0.000763 | 0.33497239 | -1.57789 |
| KCNA4 | 0.000193 | 1.90E-05 | 0.33301953 | -1.58632 |
| EGR4 | 0.00048 | 5.98E-05 | 0.3320636 | -1.59047 |
| ZNF831 | 0.000381 | 4.47E-05 | 0.3313299 | -1.59366 |
| SOHLH1 | 2.08E-06 | 4.90E-08 | 0.33115001 | -1.59444 |
| OPRK1 | 1.02E-05 | 4.36E-07 | 0.33105867 | -1.59484 |
| SLC6A15 | 4.03E-06 | 1.23E-07 | 0.33100146 | -1.59509 |
| CACNA1I | 5.24E-06 | 1.77E-07 | 0.33058473 | -1.59691 |
| ARHGDIG | 6.42E-06 | 2.35E-07 | 0.32994199 | -1.59972 |
| ICAM5 | 1.78E-05 | 8.98E-07 | 0.3295027 | -1.60164 |
| OCA2 | 9.55E-07 | 1.63E-08 | 0.32941684 | -1.60201 |
| KCTD4 | 6.57E-05 | 4.81E-06 | 0.32842483 | -1.60636 |
| ABCC12 | 1.55E-06 | 3.23E-08 | 0.32816971 | -1.60749 |
| SH3GL2 | 0.00013 | 1.14E-05 | 0.32666564 | -1.61411 |
| DLGAP1-AS4 | 9.73E-06 | 4.08E-07 | 0.326578 | -1.6145 |
| FAM19A2 | 1.34E-05 | 6.19E-07 | 0.32514573 | -1.62084 |
| LRTM2 | 1.54E-05 | 7.45E-07 | 0.3246705 | -1.62295 |
| DPP10-AS1 | 0.0042 | 0.000855 | 0.32465565 | -1.62302 |
| FSTL5 | 2.82E-05 | 1.62E-06 | 0.32452036 | -1.62362 |
| TMEM155 | 7.14E-08 | 3.18E-10 | 0.32444178 | -1.62397 |
| GABRA4 | 0.000152 | 1.40E-05 | 0.32433803 | -1.62443 |
| RBM11 | 3.13E-06 | 8.63E-08 | 0.32413035 | -1.62535 |
| FEZF2 | 0.000373 | 4.35E-05 | 0.32409323 | -1.62552 |
| TMEM132D | 1.60E-06 | 3.37E-08 | 0.32398008 | -1.62602 |
| GABRA2 | 0.0015 | 0.000242 | 0.32396276 | -1.6261 |
| KSR2 | 2.31E-05 | 1.24E-06 | 0.32344547 | -1.62841 |
| KLHL1 | 1.34E-06 | 2.59E-08 | 0.32323018 | -1.62937 |
| DDN | 2.48E-08 | 4.67E-11 | 0.32231602 | -1.63345 |
| PHF24 | 6.84E-06 | 2.55E-07 | 0.32058288 | -1.64123 |
| WSCD2 | 1.86E-05 | 9.53E-07 | 0.31991357 | -1.64425 |
| TUNAR | 0.000512 | 6.47E-05 | 0.31987914 | -1.6444 |
| NPM2 | 2.97E-05 | 1.74E-06 | 0.31951368 | -1.64605 |
| SERPINI1 | 1.53E-06 | 3.16E-08 | 0.31944561 | -1.64636 |
| NRGN | 2.42E-06 | 6.02E-08 | 0.31929753 | -1.64703 |
| ETNPPL | 0.00296 | 0.000556 | 0.31929702 | -1.64703 |
| SLITRK4 | 1.00E-05 | 4.22E-07 | 0.31901817 | -1.64829 |
| FAM19A1 | 0.000164 | 1.54E-05 | 0.31879038 | -1.64932 |
| BHLHE22 | 0.000408 | 4.88E-05 | 0.31815601 | -1.65219 |
| PHYHIP | 1.62E-06 | 3.41E-08 | 0.31708598 | -1.65705 |
| SYN2 | 3.34E-06 | 9.47E-08 | 0.31691629 | -1.65783 |

| | | | | |
|---|---|---|---|---|
| NPTX1 | 0.000121 | 1.04E-05 | 0.31602106 | -1.66191 |
| GREM2 | 0.000142 | 1.29E-05 | 0.31529439 | -1.66523 |
| GLS2 | 2.97E-07 | 2.79E-09 | 0.31484549 | -1.66728 |
| HOOK1 | 1.10E-06 | 1.97E-08 | 0.31415199 | -1.67047 |
| RXFP1 | 1.22E-06 | 2.29E-08 | 0.31357525 | -1.67312 |
| TMEM130 | 0.000128 | 1.12E-05 | 0.31345473 | -1.67367 |
| TYRP1 | 2.62E-06 | 6.70E-08 | 0.31334618 | -1.67417 |
| SLC6A17 | 5.39E-06 | 1.84E-07 | 0.31256378 | -1.67778 |
| ANO3 | 0.000147 | 1.35E-05 | 0.31217894 | -1.67955 |
| CDH9 | 0.000103 | 8.58E-06 | 0.31174162 | -1.68158 |
| RFPL2 | 1.47E-07 | 9.51E-10 | 0.31079188 | -1.68598 |
| KCNJ4 | 0.000133 | 1.17E-05 | 0.31038846 | -1.68785 |
| CACNA1B | 1.19E-05 | 5.32E-07 | 0.31012072 | -1.6891 |
| MFSD4A | 2.00E-07 | 1.53E-09 | 0.30987522 | -1.69024 |
| RAB3C | 0.000436 | 5.31E-05 | 0.30980489 | -1.69057 |
| SYT1 | 9.47E-05 | 7.70E-06 | 0.30958228 | -1.69161 |
| CAMKV | 4.74E-05 | 3.15E-06 | 0.30764066 | -1.70068 |
| GLT1D1 | 4.23E-05 | 2.72E-06 | 0.30748849 | -1.7014 |
| DRD1 | 9.32E-08 | 4.73E-10 | 0.3074051 | -1.70179 |
| RBP4 | 5.99E-06 | 2.13E-07 | 0.30678573 | -1.7047 |
| GALNTL5 | 0.000303 | 3.36E-05 | 0.30639752 | -1.70652 |
| CPLX3 | 9.87E-07 | 1.70E-08 | 0.30632039 | -1.70689 |
| CHD5 | 1.06E-05 | 4.53E-07 | 0.30591692 | -1.70879 |
| SYT13 | 0.000299 | 3.30E-05 | 0.30539903 | -1.71123 |
| SYN1 | 2.43E-06 | 6.08E-08 | 0.30488194 | -1.71368 |
| VIP | 4.78E-06 | 1.56E-07 | 0.30431041 | -1.71638 |
| KIAA1644 | 0.000126 | 1.10E-05 | 0.30407224 | -1.71751 |
| C11orf87 | 1.90E-05 | 9.82E-07 | 0.30372132 | -1.71918 |
| SULT4A1 | 1.27E-06 | 2.41E-08 | 0.3033995 | -1.72071 |
| NPY | 5.08E-05 | 3.46E-06 | 0.3030277 | -1.72248 |
| GNG3 | 4.87E-05 | 3.27E-06 | 0.30292824 | -1.72295 |
| ADCY2 | 6.17E-06 | 2.21E-07 | 0.30269123 | -1.72408 |
| C1QL3 | 1.13E-06 | 2.04E-08 | 0.302631 | -1.72437 |
| CPLX2 | 7.12E-05 | 5.34E-06 | 0.3026244 | -1.7244 |
| RTN4RL1 | 3.78E-05 | 2.36E-06 | 0.302465 | -1.72516 |
| MPPED1 | 1.81E-06 | 3.99E-08 | 0.30061285 | -1.73402 |
| HTR2C | 1.24E-06 | 2.33E-08 | 0.30035592 | -1.73526 |
| LINC01106 | 5.04E-05 | 3.42E-06 | 0.29989415 | -1.73747 |
| GRIN3A | 2.22E-06 | 5.32E-08 | 0.29835803 | -1.74488 |
| AK5 | 3.15E-05 | 1.88E-06 | 0.29827488 | -1.74529 |
| HCN1 | 1.13E-05 | 4.97E-07 | 0.29567431 | -1.75792 |
| NEFL | 0.00132 | 0.000205 | 0.29553449 | -1.7586 |
| MYT1L | 0.000151 | 1.40E-05 | 0.29526407 | -1.75992 |

| | | | | |
|---|---|---|---|---|
| HS3ST4 | 7.81E-07 | 1.21E-08 | 0.29419369 | -1.76516 |
| SMPX | 4.68E-05 | 3.09E-06 | 0.2938132 | -1.76703 |
| PTPRR | 8.10E-05 | 6.31E-06 | 0.29335488 | -1.76928 |
| PCSK2 | 0.000316 | 3.54E-05 | 0.29254008 | -1.77329 |
| CUX2 | 0.000656 | 8.75E-05 | 0.2924666 | -1.77366 |
| TBR1 | 2.84E-08 | 6.36E-11 | 0.2915248 | -1.77831 |
| ATP2B3 | 1.44E-06 | 2.91E-08 | 0.29146953 | -1.77858 |
| SYT4 | 0.000124 | 1.08E-05 | 0.29131936 | -1.77933 |
| KCNJ3 | 1.25E-05 | 5.67E-07 | 0.29120869 | -1.77987 |
| PPP4R4 | 0.000135 | 1.20E-05 | 0.29113932 | -1.78022 |
| HTR2A | 0.000169 | 1.60E-05 | 0.29094801 | -1.78117 |
| RFPL1S | 1.43E-06 | 2.84E-08 | 0.290878 | -1.78151 |
| LINGO2 | 3.65E-05 | 2.26E-06 | 0.29079759 | -1.78191 |
| RBFOX1 | 7.04E-06 | 2.65E-07 | 0.28994792 | -1.78613 |
| SLC30A3 | 3.15E-06 | 8.72E-08 | 0.28946744 | -1.78853 |
| DLGAP2 | 2.69E-06 | 6.95E-08 | 0.28875808 | -1.79207 |
| SNAP25 | 2.61E-06 | 6.67E-08 | 0.28731137 | -1.79931 |
| VSTM2A | 4.51E-05 | 2.96E-06 | 0.28647177 | -1.80354 |
| STYK1 | 3.53E-05 | 2.16E-06 | 0.28627687 | -1.80452 |
| CALY | 2.57E-05 | 1.43E-06 | 0.28403939 | -1.81584 |
| TESPA1 | 2.28E-09 | 6.57E-13 | 0.28317152 | -1.82025 |
| MICAL2 | 2.22E-05 | 1.18E-06 | 0.28310735 | -1.82058 |
| CABP1 | 2.22E-06 | 5.30E-08 | 0.28142847 | -1.82916 |
| GUCA1B | 8.83E-06 | 3.62E-07 | 0.2791509 | -1.84088 |
| SLC26A4-AS1 | 3.47E-07 | 3.54E-09 | 0.27701152 | -1.85198 |
| GPR26 | 0.000215 | 2.18E-05 | 0.27542022 | -1.86029 |
| RYR2 | 2.77E-05 | 1.59E-06 | 0.27447543 | -1.86525 |
| CREG2 | 2.68E-05 | 1.52E-06 | 0.27235605 | -1.87643 |
| SERTM1 | 6.03E-06 | 2.15E-07 | 0.26678005 | -1.90628 |
| KCNV1 | 4.52E-06 | 1.45E-07 | 0.26526884 | -1.91447 |
| CBLN2 | 3.03E-05 | 1.78E-06 | 0.26514185 | -1.91516 |
| OLFM3 | 1.98E-07 | 1.49E-09 | 0.26479656 | -1.91704 |
| GDA | 0.00027 | 2.90E-05 | 0.26271276 | -1.92844 |
| CARTPT | 3.79E-07 | 4.08E-09 | 0.26047278 | -1.9408 |
| CDH18 | 0.00261 | 0.000478 | 0.25923372 | -1.94767 |
| CRYM | 7.98E-05 | 6.17E-06 | 0.25700375 | -1.96014 |
| GJB6 | 3.05E-05 | 1.80E-06 | 0.25661361 | -1.96233 |
| GABRA1 | 3.78E-05 | 2.36E-06 | 0.25649864 | -1.96298 |
| CCKBR | 3.73E-06 | 1.09E-07 | 0.25646615 | -1.96316 |
| SSTR1 | 4.13E-05 | 2.65E-06 | 0.25438179 | -1.97493 |
| CAMK2A | 1.84E-05 | 9.43E-07 | 0.25402893 | -1.97694 |
| GAD2 | 0.00132 | 0.000207 | 0.25392916 | -1.9775 |
| SST | 1.94E-05 | 1.01E-06 | 0.25169886 | -1.99023 |

| | | | | |
|---|---|---|---|---|
| RBFOX3 | 8.20E-05 | 6.42E-06 | 0.250728 | -1.99581 |
| NEFM | 0.000348 | 4.00E-05 | 0.24747258 | -2.01466 |
| NWD2 | 1.51E-06 | 3.11E-08 | 0.24373983 | -2.03659 |
| SV2B | 9.03E-05 | 7.27E-06 | 0.24132704 | -2.05094 |
| GABRA5 | 1.67E-05 | 8.29E-07 | 0.24006013 | -2.05853 |
| SVOP | 0.000212 | 2.15E-05 | 0.23881699 | -2.06602 |
| KRT222 | 3.63E-05 | 2.24E-06 | 0.23798055 | -2.07108 |
| SLC12A5 | 6.66E-05 | 4.90E-06 | 0.23750879 | -2.07395 |
| KLK7 | 3.83E-08 | 1.16E-10 | 0.23437978 | -2.09308 |
| LY86-AS1 | 2.79E-06 | 7.31E-08 | 0.23340182 | -2.09911 |
| NEUROD6 | 1.21E-07 | 7.09E-10 | 0.23176336 | -2.10928 |
| FRMPD4 | 4.86E-06 | 1.60E-07 | 0.22975368 | -2.12184 |
| CCK | 0.000916 | 0.000132 | 0.22650338 | -2.1424 |
| VSNL1 | 0.000857 | 0.000122 | 0.22498654 | -2.15209 |
| SYNPR | 0.00133 | 0.000208 | 0.22395313 | -2.15873 |
| PACSIN1 | 1.12E-05 | 4.89E-07 | 0.22249531 | -2.16815 |
| FRMPD2 | 1.89E-06 | 4.32E-08 | 0.21959982 | -2.18705 |
| OPALIN | 1.14E-07 | 6.54E-10 | 0.21742817 | -2.20139 |
| MAL2 | 5.06E-05 | 3.45E-06 | 0.20832544 | -2.26309 |
