## Supplemental Table 2 for "Mechanotherapeutic Potential of Survivin in Glioblastoma"

**Supplementary Table 2. GBM recruited patients' demographics.** Demographic variables, comorbidities, family history, and tumor characteristics of the study cohort. The average patient age was 59.3 years; all individuals identified as White/Caucasian, and 60% were male. Current smoking was reported in 30% of patients. Hypertension and arthritis were the most prevalent comorbidities (30% each), while no patients reported heart disease, stroke, high cholesterol, cancer, diabetes, or asthma. Family history included intracranial aneurysm and heart disease in 30% of patients. All tumors were primary GBM and IDH WT. *Except for age (mean), all values are percentages. Abbreviations: IDH WT, isocitrate dehydrogenase wild type.

| **Demographics** | |
|---|---|
| **Age** | 59.3 |
| **Race (White/ Caucasian)** | 100% |
| **Sex (Male)** | 60% |
| **Smoking (Current)** | 30% |
| **Comorbidities** | |
| **Hypertension** | 30% |
| **Heart Disease** | 0% |
| **Stroke** | 0% |
| **High Cholesterol** | 0% |
| **Cancer** | 0% |
| **Diabetes** | 0% |
| **Arthritis** | 30% |
| **Asthma** | 0% |
| **Family History** | |
| **Intracranial Aneurysm** | 30% |
| **Stroke** | 0% |
| **Heart Disease** | 30% |
| **Diabetes** | 0% |
| **Prior Brain Tumor** | 0% |
| **History of Cancer** | 0% |
| **Tumor Data** | |
| **Primary GBM** | 100% |
| **IDH WT** | 100% |
