## Supplemental Table 3 for "Mechanotherapeutic Potential of Survivin in Glioblastoma"

**Supplementary Table 3. NeuroBioBank control brain tissue donor demographics.** Control human brain autopsies from non-GBM donors (*n=3*) were obtained from the NIH NeuroBioBank. Demographic variables, comorbidities, relevant clinical history are provided. Tissue samples were collected from the dorsolateral prefrontal cortex, formalin-fixed, and paraffin embedded. All donors had no clinical or neuropathological brain diagnosis. *n=3.*

| Variable | Patients |
|---|---|
| **Age** | >50 |
| **Brain Region** | Dorsolateral Prefrontal cortex (Brodmann area 9) |
| **Preparation** | FFPE |
| **Hemisphere** | 66.6% Left |
| **Tissue Type** | Brain |
| **Manner of Death** | 66.6% Natural 33.3% Accidental |
| **Gender** | Male |
| **Race** | White |
| **Clinical Brain Diagnosis** | No clinical brain diagnosis found |
| **Neuropathology Diagnosis** | Diagnostic pathology not present |
